# Protein G-quadruplex interactions and their effects on phase transitions and protein aggregation

**DOI:** 10.1101/2023.09.21.558871

**Authors:** Bikash R. Sahoo, Vojč Kocman, Nathan Clark, Nikhil Myers, Xiexiong Deng, Ee L. Wong, Harry J. Yang, Anita Kotar, Bryan B. Guzman, Daniel Dominguez, Janez Plavec, James C.A. Bardwell

**Affiliations:** Howard Hughes Medical Institute; Department of Molecular, Cellular and Developmental Biology, University of Michigan, Ann Arbor, MI, USA; National Institute of Chemistry, Ljubljana, Slovenia; Department of Pharmacology, UNC Chapel Hill, USA

## Abstract

The SERF family of proteins were originally discovered for their ability to accelerate amyloid formation. Znf706 is an uncharacterized protein whose N-terminus is homologous to SERF proteins. We show here that human Znf706 can promote protein aggregation and amyloid formation. Unexpectedly, Znf706 specifically interacts with stable, non-canonical nucleic acid structures known as G-quadruplexes. G-quadruplexes can affect gene regulation and suppress protein aggregation; however, it is unknown if and how these two activities are linked. We find Znf706 binds preferentially to parallel G-quadruplexes with low micromolar affinity, primarily using its N-terminus, and upon interaction, its dynamics are constrained. G-quadruplex binding suppresses Znf706’s ability to promote protein aggregation. Znf706 in conjunction with G-quadruplexes therefore may play a role in regulating protein folding. RNAseq analysis shows that Znf706 depletion specifically impacts the mRNA abundance of genes that are predicted to contain high G-quadruplex density. Our studies give insight into how proteins and G-quadruplexes interact, and how these interactions affect both partners and lead to the modulation of protein aggregation and cellular mRNA levels. These observations suggest that the SERF family of proteins, in conjunction with G-quadruplexes, may have a broader role in regulating protein folding and gene expression than previously appreciated.

## INTRODUCTION

Structural biology and biochemistry have been used to classify a wide variety of protein families based on their three-dimensional structures and functions. The sequence properties of these proteins driving disorder and their overall importance in protein function have been the subject of innumerable reviews (1). However, a surprisingly sizable percentage of proteins, including those in broadly conserved families, still lack clearly defined functions. This is particularly an issue for proteins that contain regions of disorder (2–5). Many of these proteins have been reported to play roles in macromolecular interactions, liquid-liquid phase separations, subcellular localization, regulation of gene expression, aggregation, and disease, but many still lack clearly defined functions (2, 6–9). The inherent difficulty in studying these proteins is their inability to fold into an average 3D structure making them particularly challenging research subjects. While some details are emerging about how disordered proteins interact with each other (10), there is limited availability of high-resolution structural information of them in isolation which is a prerequisite for understanding how they interact with their binding partners.

We decided to study one such family of partially disordered proteins, namely the 4F5 family (Pfam: PF04419). The founding member of this family, the Small ERDK-rich Factor (SERF), was discovered for its ability to accelerate amyloid aggregation (11). These proteins form fuzzy complexes with amyloid precursors, which accelerate amyloid nucleation (12). SERF-related proteins are characterized as having high net charge, a high degree of conservation, and partial structural disorder, but their roles in normal cell physiology are unclear (11–16).

Around 40% of the 4F5 family of proteins are characterized by an N-terminal SERF-homologous domain and a C-terminal single C2H2 type zinc-finger domain (15). The most common zinc-finger proteins interact with nucleic acid structures to regulate gene expression using multiple zinc-finger domains (17). We focused our research on a protein containing a single zinc finger, the ∼8. 5 kDa human Znf706 protein. Though this protein has family members in a wide variety of eukaryotic organisms, little is known about their functions (18, 19)

Presented here is evidence that Znf706 specifically interacts with G-quadruplexes. G-quadruplexes are non-canonical, guanine-rich nucleic acid structures that form G-tetrads. These tetrads are formed by planar Hoogsteen base-paired arrangements of four guanine nucleotides stabilized by a monovalent cation (K^+^/Na^+^). They can stack over one another through π-π integration to form compact and ordered structures (20). G-quadruplexes have been detected in telomeres and the untranslated regions of many genes (21). They are thought to regulate telomere function and cellular processes such as gene regulation and mRNA stability, particularly under stress conditions (20, 22–24). Several proteins are known to regulate G-quadruplex function. Helicases, such as DHX36, DDX5, FANCJ, POT1, and RPA are capable of unfolding G-quadruplexes (25–29), and nucleolin, p53, DNMT1, along with several zinc-finger proteins, including Sp1 and MAZ, are known to bind to and stabilize G-quadruplexes (30). G-quadruplex structure and polymorphism have been extensively studied, and several high-resolution structures of G-quadruplexes complexed with proteins have been reported including with DHX36 helicase (26), yeast telomeric protein Rap1 (31), and telomere end binding protein in *Oxytricha nova* (32). However, the structural characteristics of G-quadruplexes that allow for protein recognition, the dynamic changes that occur in both the G-quadruplex and protein partners upon binding, and how binding affects the activity of both partners remain unclear. There are however intriguing links emerging between G-quadruplexes and certain protein folding diseases (33). G-quadruplexes have been found in amyloid aggregates and several amyloidogenic proteins appear to specifically bind G-quadruplexes (33, 34). Our research utilized a variety of biophysical tools, including NMR spectroscopy, to perform a detailed characterization of protein and G-quadruplex interactions and to determine the structural and dynamic changes that occur in both binding partners upon interaction. Based on these findings, we have proposed a model as to how these interactions may affect G-quadruplex and Znf706’s activities in controlling protein aggregation and in vivo mRNA levels.

## MATERIALS AND METHODS

### Recombinant human Znf706 and *α*-Synuclein protein expression and purification

The human Znf706 sequence was codon-optimized for *E. coli* expression and cloned into the pET28 derived vector containing an N-terminal His_6_-SUMO tag (pET-HisSumo) or an enhanced green fluorescent protein tag (eGFP). Wild-type Znf706 protein or variants, containing A2C or A76C substitutions were expressed as His_6_-SUMO fusion proteins. These His_6_-SUMO and eGFP tagged Znf706 proteins were expressed in *E. coli* BL21 DE3 using PEM media at 37 °C until the culture density reached OD600 of 1. 0 followed by an overnight induction using 0. 2 M IPTG at a culture temperature of 22 °C with shaking at 160 rpm. For recombinant expression of uniformly labeled ^15^N or ^15^N, ^13^C His_6_-SUMO tagged Znf706, M-9 minimal media supplemented with ^15^N and ^15^N, ^13^C isotope labeled ammonium chloride (^15^NH_4_Cl, 1 g/L) and glucose (D-Glucose-6-^13^C, 4 g/L) as sole nitrogen and carbon sources, respectively, were used instead of PEM media. Isotopes were purchased from Cambridge Isotope Laboratories. Tagged proteins were purified by passage through a nickel affinity HisTrap column (Cytiva, #17524802) followed by HisSUMO tag cleavage with ULP1 protease in 40 mM Tris, 10 mM sodium phosphate, 300 mM NaCl, pH 8. 0, and dialyzed overnight at 4 °C. The resulting tag-free Znf706 proteins were then purified by removing the cleaved His-tag containing fragments by absorption onto a nickel affinity resin and collecting the flow-through. Further purification occurred by subjecting the samples to an anion exchange column (HiTrap SP, Cytiva, 17115201) followed by size-exclusion chromatography (HiLoad 16/60 Superdex S75; GE Healthcare, 17106801) as previously described (12). The protein sample stocks were prepared either in 20 mM sodium phosphate (NaPi, pH 7. 4), 100 mM KCl, 0. 05% NaN_3,_ or 20 mM Tris-HCl (pH 7. 4), 100 mM LiCl, 0. 05% NaN_3_. For analytical ultracentrifugation (AUC), eGFP (A206K) and eGFP (A206K) fused Znf706 proteins were prepared using the pET-HisSumo construct. The expression and purification of wild-type and A90C α-Synuclein protein was as previously described (12). Lyophilized α-Synuclein protein was dissolved in 20 mM NaPi, 100 mM KCl, 0. 05% NaN_3_, pH 7. 4 for use in the amyloid aggregation assay. Protein concentrations were measured by UV absorption using a UV-1900i (Shimadzu) spectrophotometer and the molar extinction coefficients were calculated using the ExPASy ProtParam tool. A zinc-specific assay using the chromophoric chelator 4-(2-pyridylazo) resorcinol was used to measure zinc coordination to Znf706 and compared with a known concentration of ZnCl2 used as a standard in this assay. The C41 and C44 cysteines in Znf706 were reduced by para-chloromercuribenzoic acid and compared with Znf706 under a non-reducing condition.

### Fluorescent and nitroxide spin labeling

Znf706 A2C or A76C and α-Synuclein A90C cysteine mutant proteins were purified as described above, and used for Alexa Fluor 488 ((AF488), CAS-A10254) or 1-Oxyl-2, 2, 5, 5-tetramethyl-Δ3-pyrroline-3-methyl methanethiosulfonate ((MTSL), CAS: 81213-52-7) labeling. To perform the labeling, 200 µM of cysteine labeled Znf706 A2C (dissolved in 20 mM Tris-HCl, 0. 2 mM ZnCl_2_, pH 8. 0) or α-Synuclein A90C (dissolved in 20 mM Tris-HCl, 0. 1 mM EDTA, pH 8. 0) proteins were incubated with a 10-fold molar excess of AF488 overnight at 37 °C (Znf706 A2C) or 4 °C (α-Synuclein A90C) with shaking at 250 rpm. Znf706 A2C or A76C were labeled with MTSL (for Znf706 A2C or A76C) for NMR studies. The excess AF488 or MTSL unincorporated label was removed by passing the samples over a PD-10 desalting column (Cytiva, 17085101). The sample buffer was next changed to 20 mM NaPi, 100 mM KCl, 0. 05% NaN_3_, pH 7. 4 by either running the protein on a size-exclusion chromatography column (Superdex S75 10/300 GL, Cytiva) or by concentrating the sample 20 times using a 3 kDa cut-off filter (Amicon^®^, UFC500324). The AF488 labeling efficiency was determined by UV absorbance following the manufacturer’s protocol. An analog of MTSL which has no PRE effect i. e., N-ethylmaleimide (NEM) labeled A2C Znf706 was also prepared following the above-described procedure for NMR studies.

### Oligonucleotide synthesis and G-quadruplex formation

Desalted G-rich DNA oligonucleotides, DNA polynucleotides (A/T/C/G/N), and G-quadruplex forming DNA and RNA oligonucleotides were purchased either from Integrative DNA Technology (IDT) or synthesized in-house as described below. 5’ 6-FAM labeled oligonucleotide sequences were purchased from IDT as HPLC purified, desalted DNA or RNA oligonucleotides. These oligonucleotides were dissolved in nuclease-free water and further desalted by 15 cycles of centrifugation using nuclease-free water in a 3 kDa cut-off filter (Amicon^®^, UFC500324). Oligonucleotide concentration was determined using the extinction coefficients obtained using IDT Oligoanalyzer tool. Oligonucleotides were also synthesized in-house using a K&A Laborgeraete GbR H-8 DNA/RNA Synthesizer. Standard phosphoramidite chemistry was used. Deprotection was done using aqueous ammonia at 55 °C for 24 h. Samples were purified and desalted by 15 cycles of concentration in water using a 3 kDa cut-off filter (Amicon^®^, UFC500324). The concentration of the oligonucleotides was determined from their predicted extinction coefficients, obtained from IDT Oligoanalyzer tools, using a UV-1900i (Shimadzu) spectrometer. Oligonucleotides were next folded in either 20 mM NaPi, 100 mM KCl, 0. 05% NaN_3_, pH 7. 4 or 20-mM Tris-HCl, 100 mM LiCl, 0. 05% NaN_3_, pH 7. 4 and subjected to a heating and cooling cycle using a thermal cycler heating to 98 °C for 5 minutes, then cooling from 75 to 4 °C at a rate of 1 °C/minute.

### Fluorescence polarization and anisotropy

The binding of oligonucleotides to Znf706 was determined at 25 °C by measuring the change in fluorescence polarization or anisotropy using a Tecan Infinite M1000 Pro plate reader or a Cary Eclipse fluorophotometer, respectively. Fluorescence anisotropy measurements were done using 200 nM of AF488 labeled Znf706 (A2C) and increasing concentrations of unlabeled DNA, RNA, and G-quadruplex forming oligonucleotides. For fluorescence polarization measurements, 20 µL of the sample containing 100 nM of 5’ 6-FAM labeled G-quadruplexes and increasing concentrations of unlabeled Znf706 were incubated for 1 hour before fluorescence polarization measurements using a 384 well plate (Corning #4514). The binding dissociation constant of G-quadruplexes to Znf706 was determined by a non-linear regression curve fit, using one site binding-saturation model in GraphPad Prism. All the fluorescence assays were carried out in triplicate in 20 mM NaPi, 100 mM KCl, 0. 05% NaN_3_, pH 7. 4 except for those with unfolded G-quadruplexes, which were done in 20 mM Tris-HCl, 100 mM LiCl, 0. 05% NaN_3_, pH 7. 4 or 20 mM NaPi, 100 mM NaCl, pH 7. 4 for G4C2 4-repeat sequence. The binding of α-Synuclein and Znf706 was studied by fluorescence polarization using either 100 nM of AF488 labeled α-Synuclein A90C and increasing concentrations of unlabeled Znf706 or 100 nM of AF488 labeled Znf706 A2C and increasing concentrations of unlabeled α-Synuclein.

### Microscale thermophoresis

20 nM of 5’ 6-FAM labeled Kit*, the human telomeric G-quadruplexes hTel and TERRA were preincubated with increasing concentrations of Znf706 in 20 mM NaPi, 100 mM KCl, 0. 05% NaN_3_, pH 7. 4 for ∼1 hour. This was followed by microscale thermophoresis measurements performed at 25 °C using a Monolith NT. 115 (NanoTemper) instrument using the blue filter, 10-20% LED, and 80% microscale thermophoresis power.

### Circular Dichroism spectroscopy

Znf706 secondary structure was studied using a JASCO J-1500 or a JASCO J-820 spectropolarimeter. Far-UV Circular Dichroism (CD) spectra were collected using a 200 µL quartz cuvette with a path length of 1 mm. CD spectrum of Znf706 (50 µM) dissolved in 20 mM NaPi, 4 mM KCl, and pH 7. 4 was collected as an average of 8 scans. The effect of zinc coordination on Znf706 secondary structure was examined by measuring the CD spectrum of Znf706 (50 µM) preincubated with a 20×molar excess of EDTA for 15 minutes at 25 °C. Znf706 thermal stability was monitored by recording its far-CD absorptions at 208 and 222 nm while subsequently heating the sample from 5 °C to 95 °C. The CD data were collected by averaging 2 scans/temperature for each 5 °C time shift with a 3-minute delay following each temperature shift.

The binding effect of Znf706 and G-quadruplex was studied by CD spectroscopy at 25 °C. We averaged data collected from 8 scans. 20 µM and 40 µM of Znf706 were titrated with 20 µM cMyc or Bcl2WT G-quadruplexes dissolved in 20 mM NaPi, 100 mM KCl, pH 7. 4. The buffer baseline signal was subtracted from the mixture sample while monitoring G quadruplex structural changes. To monitor the secondary structural changes in Znf706, the G-quadruplex signals were subtracted from the mixture sample. To study the effect of Znf706’s binding on G-quadruplexes melting temperature, 20 µM G-quadruplexes (pre-folded in 20 mM NaPi, 100 mM KCl, pH 7. 4) were resuspended in 20 mM NaPi, 4 mM KCl, pH 7. 4 in the absence or presence of an equimolar or 5×molar excess of Znf706. Melting experiments were done using the same settings we used for the protein only samples, as described above. To monitor the influence of Znf706 on the refolding process of G-quadruplexes, G-quadruplexes mixed with or without Znf706 samples dissolved in 20 mM NaPi, 4 mM KCl, pH 7. 4, were first melted at 95 °C and cooled at 10 °C/min followed by CD measurements at 25 °C. CD spectra of 20 µM TERRA dissolved in 20 mM NaPi, 100 mM KCl, pH 7. 4 or 20 mM Tris, 100 mM LiCl, pH 7. 4, and 20 µM G4C2 4-repeat DNA sequence dissolved in 20 mM NaPi, 100 mM KCl, pH 7. 4 or 20 mM NaPi, 100 mM NaCl, pH 7. 4 were measured at 25 °C averaging data collected from 8 scans.

### Analytical Ultracentrifugation

The sedimentation velocity of various Znf706 and G-quadruplex complexes was determined using analytical ultracentrifugation (AUC). The AUC measurements were done by comparing two different samples; type-1, which contained unlabeled Znf706, and type-2, which contained Znf706 labeled with eGFP (A206K). The type-1 samples were monitored by measuring the absorption of the cMyc G-quadruplex at 260 nm. In some cases, eGFP(A206K) labeled Znf706 was used. This is because the single tyrosine in Znf706 absorbs weakly at 260 nm interfering with the distinct observation of the G-quadruplex in AUC analysis. To bypass this issue, we also used eGFP-Znf706 and monitored the protein absorption at 488 nm in the type-2 samples. This wavelength filters out the G-quadruplex absorbance, making it a desirable method for studying complex size in AUC analysis. 420 µL of cMyc G-quadruplex samples with an OD_260_ of 0. 35 or 0. 7, were prepared in the presence or absence of either an equimolar or a 2×molar excess of Znf706. All components were dissolved in 20 mM NaPi, 100 mM KCl, 0. 05% NaN_3_, pH 7. 4, and samples were loaded onto epon-charcoal two-channel centerpieces in an An60Ti rotor in a Beckman Optima Xl-I analytical ultracentrifuge using a 1. 2-cm path length. AUC measurements were carried out in the intensity mode at 58, 000 rpm. The type-2 sample measurements were done at 32, 000 rpm at 22 °C. The sedimentation velocity data were analyzed using Ultrascan III software (version 4) and exported to the LIMS (Laboratory Information Management System) server. The data fitting was done on the LIMS server using the computing clusters available at the University of Texas Health Science Center and XSEDE (Extreme Science and Engineering Discovery Environment) sites at the Texas Advanced Computing Center. The partial specific volume of Znf706 and different G-quadruplex sequences were estimated within UltraScan III. Raw AUC intensity data were converted into pseudo-absorbance. To subtract time-invariant noise, a two-dimensional sedimentation spectrum analysis was performed. This was followed by fitting the radially invariant noise within an S range of 1 to 10, using 64 S-value grid points and 64 f/f0 grid points. The two-dimensional sedimentation spectrum analysis was iterated 10 times, after fitting the meniscus. The two-dimensional sedimentation spectrum analysis data resolution was further refined by employing genetic algorithm parameters to remove any false positive data, using 50 Monte Carlo iterations.

### NMR spectroscopy and Znf706 structure modeling

NMR measurements were done for Znf706 dissolved in either 20 mM NaPi, pH 7. 4, or 20 mM Tris-HCl, pH 7. 4, as specified. 1D proton, 2D ^15^N-^1^H TROSY experiments (at 4, 25 and 37 °C) using 16 scans, 100 t1 increments, and a 1. 5 s delay, or ^1^H/^13^C HSQC experiments using 16 scans, 128 t1 increments, and a 1. 5 s delay, and 3D (HNCA, HNCACB, CBCA(CO)NH, ^15^N-NOESY-HSQC, ^13^C-NOESY-HSQC, HNCO, and HNCOCA) heteronu-clear NMR experiments were carried out on a Bruker 800 MHz spectrometer equipped with a triple-resonance cryoprobe and operated by Topspin 4. 1. 4. A sample volume of 275 µL containing 0. 75 mM of ^15^N-^13^C Znf706 dissolved in 20 mM NaPi, 20 mM NaCl pH 7. 4, and 7. 5% D_2_O filled into a Shigemi tube was used for the 2D and 3D NMR data collections. All spectra were recorded at 4 °C, processed in Topspin 4. 1. 4, and analyzed using NMR-FAM-Sparky. The Znf706 backbone assignments were done in NMRFAM Sparky. The thermostability of Znf706 was tested by collecting 1D proton NMR spectra of 100 µM Znf706 at variable temperatures (4 °C to 65. 5°C) in the absence or presence of EDTA. NMR titration experiments were done to collect the 1D proton spectrum (1 s delay, 512 ns) of cMyc, Bcl2WT, and Bcl2SH G-quadruplexes, and G-rich oligonucleotides (Table 1) in different conditions (as specified in results), using a fixed G-quadruplex and variable Znf706 concentrations as specified in the results. 2D ^15^N-^1^H TROSY NMR experiments were conducted using 100 µM ^15^N labeled Znf706 titrated with increasing (5, 20, and 50 µM) concentrations of G-quadruplexes. 1D proton and 2D ^15^N-^1^H TROSY NMR experiments were done at 37 °C and 50 °C for a sample mixture containing 200 µM ^15^N labeled Znf706 and 200 µM of cMyc G-quadruplexes. ^15^N-^1^H TROSY spectrum of 50 µM ^15^N Znf706 mixed with 100 µM 4-repeat G4C2 oligonucleotides prepared in 20 mM NaPi, 0. 05% NaN_3_, 7. 5% D_2_O, pH 7. 4, 100 mM KCl or in 20 mM NaPi, 7. 5% D_2_O, 0. 05% NaN_3_, pH 7. 4, 100 mM NaCl was acquired at 4 °C.

**Table 1.**
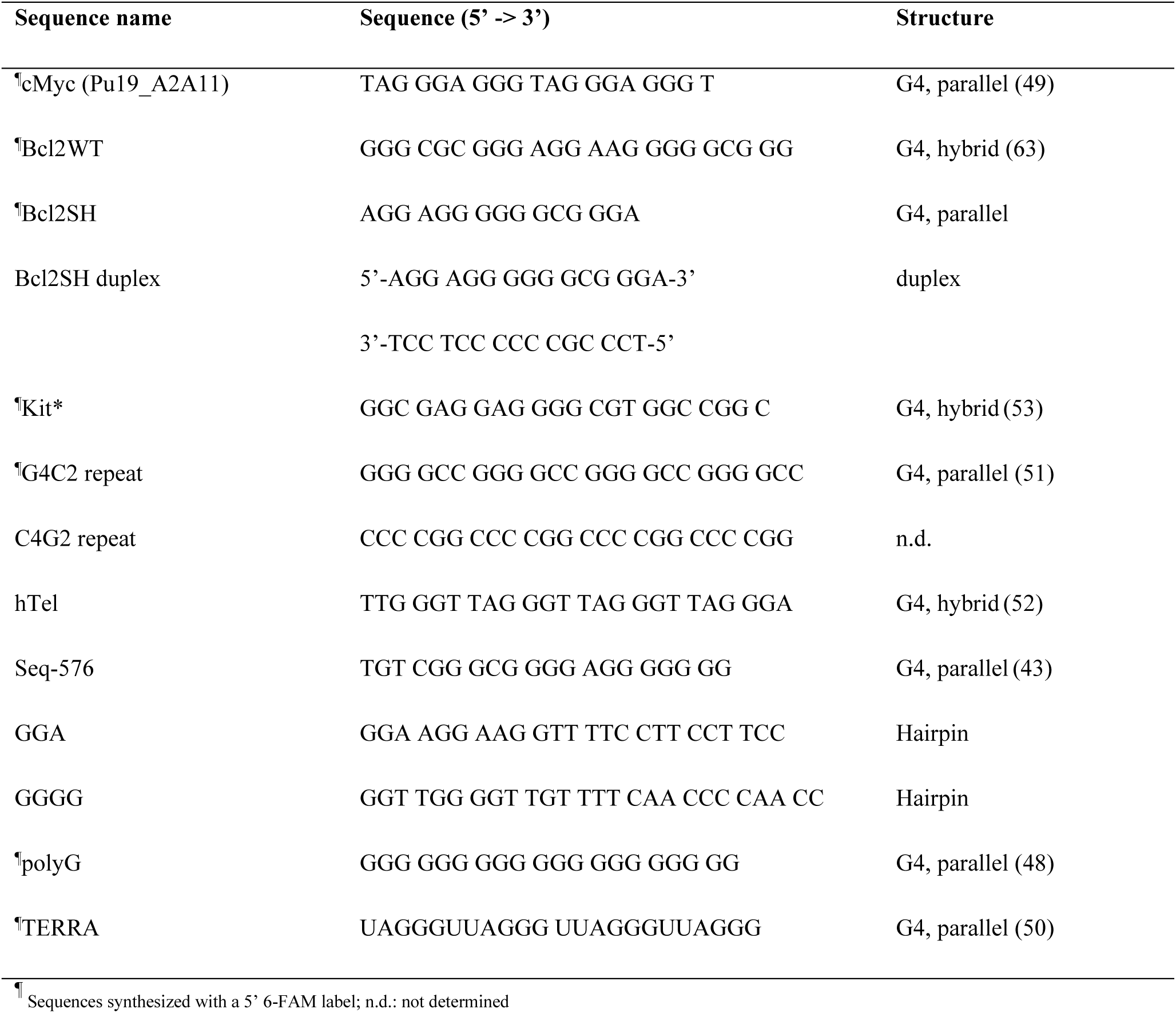
List of G-quadruplex and hairpin oligonucleotides used in this study.

2D ^15^N/^1^H TROSY and 3D HNCA, HNCACB, CBCA(CO)NH, and HNCO spectra were collected for 200 µM uniformly labeled ^13^C-^15^N Znf706 mixed with 100 µM of cMyc. The secondary structure of 200 µM Znf706 mixed without or with 100 µM cMyc was predicted from the backbone chemical shifts (C_α_, C_β_, N, NH, and CO) using the TALOS-N program (35). 200 µM ^15^N Znf706, in the absence or presence of 100 µM cMyc G-quadruplex was used for R1, R2, and hetero-NOE type relaxation studies. ^15^N relaxation experiments were carried out at 4° C on a Bruker 600 MHz spectrophotometer. The NMR spectra used for R1 analysis were collected by using the following relaxation delay times: 20, 50, 90, 130, 200, 320, 450, 580, 750, 900, 1200, and 2005 ms. R2 relaxation data were collected using the following relaxation delays: 16. 96, 50. 88, 84. 80, 135. 68, 169. 60, 220. 48, 305. 28, 373. 12, 457. 92, 542. 72, 678. 40, and 1356. 80 ms. All relaxation data were processed using TopSpin 4. 1. 1. The R1 (spin-lattice) and R2 (spin-spin) relaxation rates were determined by separating the resonance heights and fitted as a function of the relaxation delay time using NMRFAM Sparky. The hetero NOE data were collected with a relaxation delay of 2. 5 s and the spectra were analyzed using Sparky.

Paramagnetic relaxation enhancement (PRE) NMR measurements were done using 100 µM of ^15^N labeled A2C-MTSL Znf706 dissolved in 20 mM NaPi, 100 mM KCl, 7. 5% D_2_O, 0. 05% NaN_3_ (w/v) pH 7. 4 at 4° C, mixed with and without 50 µM of cMyc G-quadruplex DNA. For a comparative study, uniformly ^15^N labeled Znf706 A2C mutants were tagged with N-ethylmaleimide (NEM), and NMR data were recorded on samples containing 100 µM of A2C-NEM Znf706 mixed with 50 µM of cMyc G-quadruplex. NMR spectra of 100 µM Znf706-A2C-MTSL were recorded in paramagnetic conditions following incubation with 5-molar excess sodium ascorbate for 3 hours to obtain the corresponding diamagnetic spectrum. To compare the effect of cMyc binding on Znf706-A2C-MTSL’s paramagnetic effects, ^15^N/^1^H heteronuclear spectra were recorded first for 50 µM Znf706-A2C-MTSL titrated with 5, 10 and 20 µM of cMyc G-quadruplexes. Next, the last titrated sample containing 50 µM Znf706-A2C-MTSL and 20 µM cMyc G-quadruplexes was reduced using 20 mM sodium ascorbate for ∼ 3 hours and the NMR spectrum was acquired in the diamagnetic condition. NMR data were collected using a triple resonance cryoprobe equipped in 600 MHz or 800 MHz Bruker spectrometers, as specified in results wherever needed. All NMR spectra were processed in Topspin 4. 1. 1 and data analysis was done using NMRFAM-SPARKY 1. 47.

The 3D model structure of Znf706 was built using the backbone chemical shifts (C_α_, C_β_, N, NH, and CO) derived from 3D NMR assignments by CS-ROSETTA (36). The CS-ROSETTA model building was successful in generating a converged 10 low-energy structure ensemble of the folded non-flexible zinc-finger (ZNF) domain but failed to converge for a full-length structure because of the flexible N-terminal SERF (N-SERF) characterized with high dynamics. To successfully build a full-length structure, the lowest energy CS-ROSETTA structure of the zinc-finger domain (residues 39-76) was appended to the lowest energy CS-ROSETTA structure of the dynamic N-terminus (residues 1-38) using the advanced modeling tutorial in MODELLER 10. 1 (37) The appended structure was superimposed with the CS-ROSETTA derived zinc-finger domain, to ensure no structure change occurred. This was followed by comparing it with the NMR structure of the C2H2 type zinc-finger 12 Miz-1. The zinc ion coordination to residues Cys41, Cys44, His57, and His62 in both the Znf706 and the zinc-finger domain (residues 39-76) were docked using the High Ambiguity Driven Protein Docking (HADDOCK2. 2) program (38). A distance restraint table was parsed using a HADDOCK2. 2 server for zinc docking in Znf706. This was generated using the solution NMR structures of several human C2H2 type zinc-fingers as a reference. These included PDB IDs: 7MC3, 2LVR, 2EM3, 2EMA, 2YTB, 2EFZ and 2EQW. This coordination showed an average distance of ∼2. 2 Å and 2. 0 Å for SG (Cys) and NE2 (His) atoms, respectively, and the average distance between the Cys-Cys, Cys-His, and His-His as ∼3. 6, 3. 6 and 3. 3 Å, respectively. These average distance restraints were referred for zinc docking to Znf706 with a lower and upper distance limit set to ±0. 5 Å. 200 water-refined HADDOCK model structures were clustered and the cluster with the lowest root mean square deviation from the overall lowest-energy structure was used to represent the zinc coordinated structure of human Znf706.

### Immunofluorescence and Microscopy

HEK 293T/17 cells (ATCC® CRL-11268™) derived from human embryonic kidney, were grown in DMEM media supplemented with 2 mM glutamine, 1x Penicillin-Streptomycin solution (Gibco™ 10378016), and 10% fetal bovine serum (FBS). The cells (∼30, 000) were seeded onto coverslips, and grown overnight in 24 well plates. Cells were then fixed with 4% paraformaldehyde (Electron Microscopy Science) in PBS for 10 min, washed three times with PBS, permeabilized with 0. 1% Triton X-100 in PBS for 30 mins, and blocked with 1% BSA/PBS solution for 1h. All steps were carried out at room temperature. Thereafter, the cells were incubated with anti-Znf706 rabbit polyclonal primary antibody (1: 200 dilutions; catalog no. AV32987, Sigma-Aldrich) and mouse anti-DNA G-quadruplex structure antibody – clone BG4 (Sigma Aldrich, MABE917), diluted 500-fold in blocking buffer, incubated overnight at 4°C, and then washed 3 times with PBST (0. 1% Tween). Finally, the cells were incubated with Znf706 specific secondary antibody (Goat anti-Rabbit Alexa Fluor™ 488, Thermo Fisher, Cat# A-32731) and anti-DNA G4 specific antibody (Goat anti-Mouse Alexa Fluor™ 647, Cat#A-21235) for visualization. For co-localization studies, slides were mounted with ProLong Gold Antifade reagent with DAPI (Invitrogen). Images were captured using a Leica TCS SP8 STED 3X confocal microscope with a 100x oil objective.

The formation of liquid-liquid phase transition (LLPT) droplets possibly occurring from the combination of Znf706, and G-quadruplexes, and non-G-quadruplex forming oligonucleotides were studied by microscopy. For imaging purposes, 16-well CultureWell™ ChamberSLIP cover glasses were coated with 5% Pluronic® F-127 co-polymer surfactant overnight followed by 10 washes with 0. 3 ml sample buffer (20 mM NaPi, 100 mM KCl, 7. 5% D_2_O, pH 7. 4). Variable concentrations of Znf706 (1, 10, 20, 50, and 100 µM) prepared in 20 mM NaPi, 100 mM KCl, 7. 5% D_2_O, pH 7. 4 were mixed with equimolar concentrations of G-quadruplexes to monitor the effect of protein and G-quadruplex concentrations on droplet formation. The effect of salt on Znf706 and G-quadruplex droplet formation was studied at fixed Znf706 and G-quadruplex (100 µM) concentrations and variable salt concentrations of 0, 20, 50, 100, and 300 mM KCl. G-quadruplexes prepared in 20 mM Tris-HCl, 100 mM LiCl, pH 7. 4 were also tested. The formation of droplets was tested for 100 µM Znf706 mixed with A, T, G, and C polynucleotides. These various protein and G-quadruplex mixed samples were either incubated overnight in 5 mm thin-walled glass NMR tubes or freshly prepared in low-binding centrifuge tubes, transferred to 16-well trays, and followed by Differential Interference Contrast (DIC) microscopy or fluorescence imaging. Droplet formation of 100 µM Znf706 mixed with 50 µM of cMyc was also monitored at varied time points (1, 5, and ∼18 hours) using DIC imaging. Before setting up NMR experiments, 100 µM of Znf706 A2C samples labeled with A2C-MTSL or A2C-NEM were incubated for 5 hours with 50 µM of cMyc to test LLPT formation.

To monitor the effect of α-Synuclein on liquid droplet formation of Znf706 and G-quadruplexes, 100 µM Znf706, 100 µM α-Synuclein and 100 µM cMyc/Bcl2WT G-quadruplexes were incubated for ∼1 hour in either a non-crowding buffer of 20 mM NaPi, 100 mM KCl, pH 7. 4, or a crowding buffer containing 20 mM NaPi, 100 mM KCl, pH 7. 4, 10% PEG8000. For fluorescence imaging and FRAP analysis, 1/100^th^ of AF488 Znf706 was spiked into the sample mixture, and droplet formation was confirmed using DIC and green signals from the AF488-tagged protein. To visualize the G-quadruplexes with the droplets, 5 µM of N-methyl mesoporphyrin IX (NMM) was added to each sample well, mixed gently, incubated for ∼10 minutes, and then imaged. The effect of the cMyc G-quadruplex on 100 µM α-Synuclein LLPT was tested at variable concentrations (50, 100, and 200 µM) at room temperature in 20 mM NaPi, 100 mM KCl, pH 7. 4 containing 20 % PEG8000. The samples were imaged within ∼1 hour of sample preparation using a Nikon Ti2-E motorized inverted microscope controlled by NIS Elements software with a SOLA 365 LED light source at 100X oil immersion objective. Fluorescence images and FRAP data analysis were done using the Fiji ImageJ program.

### Western blot analysis

HEK293T cells cultured in 60 mm dishes were washed with 1x PBS and harvested using trypsin-EDTA (0. 25% trypsin, 1 mM EDTA, Gibco). The pellets were then lysed with 2×SDS sample buffer and separated by 4-15% gradient SDS-PAGE. The following antibodies were used: rabbit anti-Znf706 (catalog no. AV3298, Sigma-Aldrich) at 1:1000 dilution and rabbit anti-GAPDH (catalog no. G9545, Sigma Aldrich) at 1:2500 dilution. The secondary antibody used was LICOR IRDye 800 CW goat anti-rabbit IgG (H+L) Biosciences, 926-32211; 1:5000 dilution). Blotted membranes were scanned with a LI-COR scanner and western blot quantification analysis was performed with Image Studio Lite (LI-COR).

### RNA interference and RNA-seq

For transient siRNA knockdowns, HEK293T and HeLa cells were grown in DMEM media supplemented with 2 mM glutamine, 1x Penicillin-Streptomycin solution (Gibco™ 10378016), and 10% FBS. The cell passage numbers were kept between 5-15. 30, 000 cells were seeded onto a 6-well plate and reversely transfected with 25 nM of either non-targeting control siRNA (Horizon Discovery, D-001206-13-20) or SMARTpool siRNA targeting Znf706 (Horizon Discovery, M-021025-01-0005) using Lipofectamine RNAiMAX (Invitrogen, 13778150). The transfection media was exchanged after 24 hours, and cells were harvested after ∼48 hours of knockdown. Total RNAs were extracted using Trizol reagent (Invitrogen, 15596026) according to the manufacturer manual.

Poly(A) RNA-seq sequencing was conducted on total RNAs using services from LC Science LLC (Houston, Texas, U. S. A. ). Paired-ended 150 bp sequencing was performed on Illumina’s NovaSeq 6000 sequencing system. Three biological replicates were sequenced for each condition in different cell lines. Sequencing reads were trimmed by Cutadapt (39) and quality checked using FastQC (http://www.bioinformatics.babraham.ac.uk/projects/fastqc/). The trimmed reads were then mapped to the human genome hg38 using HISAT2 (40). StringTie (41) was used to assemble a transcriptome for each sample and to calculate fragments per kilobase of exon per million (FPKM) of mRNAs. Differential expression analysis of mRNAs was performed by DESeq2 (42). The mRNAs with a parameter of false discovery rate/q-value ≤ 0. 05 and an absolute fold change ≥ 2 were considered differentially expressed mRNAs. The observed quadruplex density of the human genome was derived from Supplementary Table 6 in Chambers et al. 2015 (21). For comparative analysis of the quadruplex density of the human genome obtained in Znf706 knockdown HEK 293T cells, we retrieved the data published by Chambers et al. 2015 (21) for DHX36 helicase knockout in HEK293 cells.

### α-Synuclein fibrillation and protein aggregation assay

α-Synuclein aggregation assays were carried out using our previously published protocol (12). Briefly, the aggregation kinetics of α-Synuclein were monitored using a Thioflavin-T (ThT) fluorescence assay (Excitation=440 nm; Emission=485 nm) using a half-area 96-well plate (Corning #3881). 100 µM (high) or 50 µM (physiological) α-Synuclein protein was dissolved in 20 mM NaPi, 100 mM KCl, pH 7. 4, and mixed with either 25 µM or 12. 5 µM ThT, respectively, and varied concentrations of Znf706 and cMyc G-quadruplexes (25, 50 and 100 µM). ThT was added to all sample wells, except those containing G-quadruplexes. Two 3-mm glass beads were added to each sample and aggregation kinetic measurements were recorded every 600 seconds under continuous agitation (orbital shaking, 3mm amplitude, 600 s) at 37 °C using a Tecan Infinite M200 plate reader.

Porcine citrate synthase was purchased Sigma #C3260-5KU) and prepared for thermal aggregation using a method described elsewhere (43). Briefly, 1 mL of citrate synthase was dialyzed in 50 mM Tris-HCl, 2 mM EDTA, pH 8. 0 at 4 °C overnight, and then concentrated using a 10-kDa cut-off filter. The protein concentration was then measured using an extinction coefficient (ɛ280=78060 M^-1^. cm^-1^) and thermal aggregation was performed using 300 nM citrate synthase dissolved in 40 mM HEPES-KOH, pH 7. 5 buffer at 48 °C under constant stirring in the absence or presence of a variable amount of Seq576, Bcl2WT and cMyc G-quadruplexes. The effect of a variable concentration of Znf706 (0. 06 - 1. 2 µM) on citrate synthase (300 nM) thermal aggregation was monitored, and final aggregate product images were taken using DIC. Light scattering was measured using a Hitachi F-4500 fluorometer with both excitation and emission wavelengths set to 360 nm and a slit width of 1 nm.

## RESULTS

### Structure of Znf706

Though the human Znf706 protein is very small, only 76 amino acids in length, it is predicted using SMART (44) and IUPred2A (45) to possess two distinct domains: a disordered, low complexity, N-terminal domain homologous to the 4F5 (SERF) family of proteins (15) (Figure 1A and Supplementary Figure S1A-C), and an ordered C-terminal domain homologous to C2H2 zinc fingers. The SERF homologous domain spans the first 36 residues and the zinc finger domain spans residues 39-62. The SERF family of proteins was initially characterized by its ability to accelerate amyloid aggregation and was later shown to bind to nucleic acids (11, 12, 46). Whether these two functions are connected remains unclear. We decided to attempt to get experimental structural information for Znf706 since the solubility and very small size of this protein make it a viable candidate for NMR-based approaches. We found the ^15^N/^1^H heteronuclear correlation spectra of Znf706 were well resolved. This allowed us to assign 57/70 of the non-proline amide peaks (Supplementary Figure S1D) observed for zinc-bound Znf706. The zinc binding in these samples was verified using a 4-(2-pyridylazo) resorcinol assay (Supplementary Figure S1E). We were also able to assign the N, Cα, Cβ, NH, and CO backbone chemical shifts (BMRB: 52308) to determine the structure of Znf706. The de novo structure obtained using CS-ROSETTA (36) from the backbone NMR chemical shifts, is shown in Figure 1B. This structure confirmed our predictions that the N-terminal domain (residue 1-36) of Znf706 is dynamic and disordered, and the C-terminus (residues 39-74) is a folded zinc-finger domain. The HADDOCK program generated the lowest energy model structure of Znf706 with zinc coordinated to the zinc-finger residues Cys41, Cys44, His57, and His62. This zinc finger is characterized by two short anti-parallel β-strands (residues 39-41 and 45-48), and two α-helices (residues 51-61 and 69-74) which closely resemble the structure of the C2H2 zinc-finger domain-12 present in Miz-1 (PDB ID: 7MC3).

**Figure 1.**
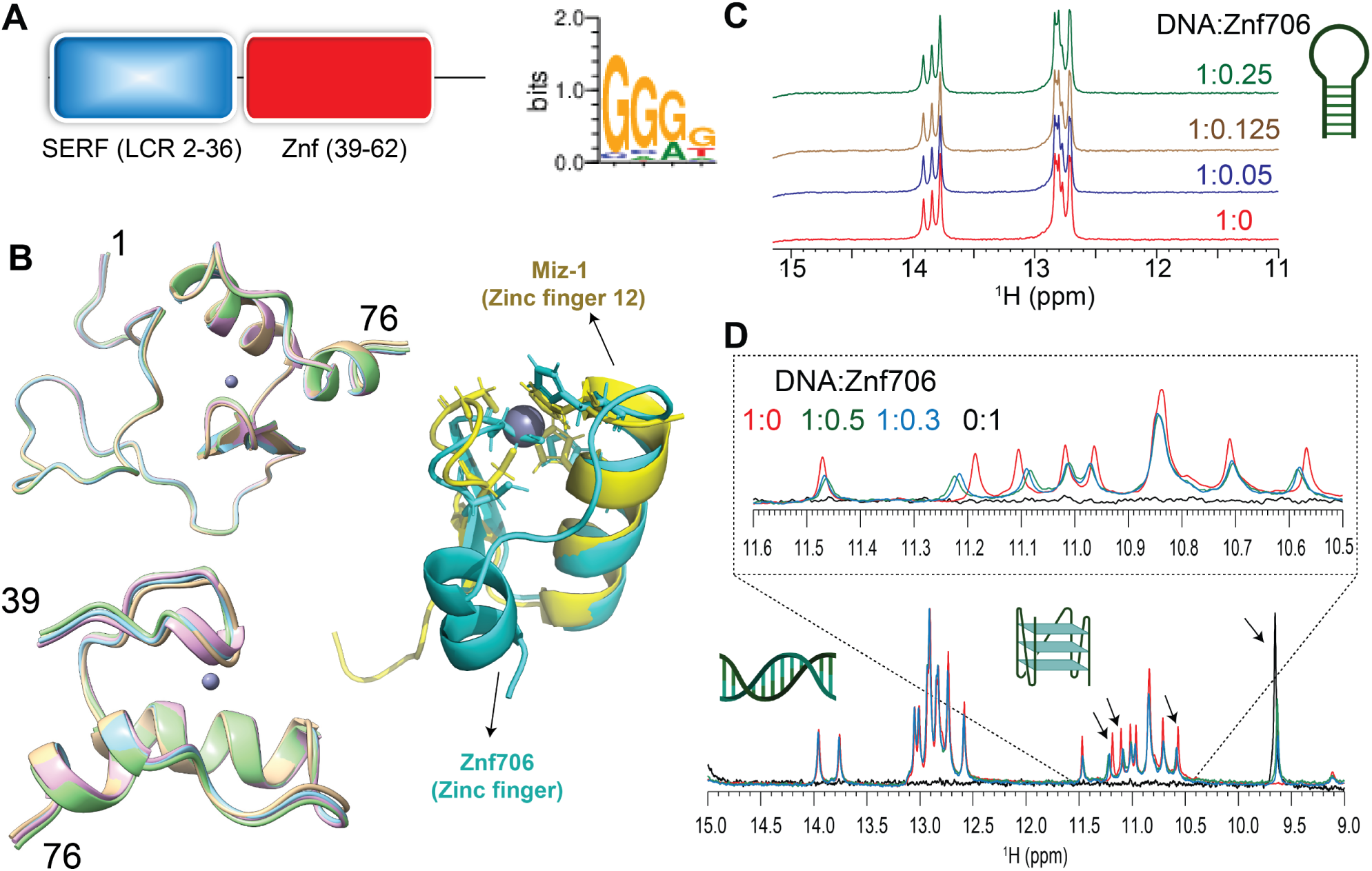
Znf706 is partially disordered and binds G-quadruplexes. **(A)** Schematic diagram showing that Znf706’s domain organization includes a conserved N-terminal low-complexity domain, colored in blue (residues 2-36) that is homologous to the SERF family and a single C2H2 type zinc-finger domain that is shown in red (residues 39-62). The DNA binding specificity of this zinc finger was predicted using the interactive PWM predictor to be GGGG, with the first residues of the motif more favored to be G than the latter ones. **(B)** De novo structures of full-length (1–76) and Znf706 (39–76) generated by CS-ROSETTA using Cα, Cβ, CO, N, and Hα NMR chemical shifts. Superimposed zinc-coordinated structures of Znf706 models were generated using HADDOCK. Cartoon structures of the CS-ROSETTA model of the human Znf706 zinc-finger domain in cyan superimposed with the solution NMR model structure of C2H2 type Miz-1 (yellow) zinc finger domain-12 (PDB ID: 7MC3). **(C)** 1D ^1^H NMR spectra showing signals of imino protons involved in Watson-Crick base pairs, with no significant change in the chemical shift observed in the presence of Znf706. This indicates no interactions between a G-rich (GGA) hairpin DNA (200 µM) and increasing concentrations of Znf706, at the indicated molar excesses of DNA relative to Znf706. **(D)** Competitive NMR titration measurements probing the binding specificity of Znf706 to G-quadruplexes and duplexes containing the same sequence of Bcl2SH and its complementary strand as listed in Table 1. The arrows indicate peaks showing substantial chemical shift changes upon Znf706 binding to the Bcl2SH G-quadruplex and the duplex mixture. The inset shows an enlarged image of the region of the Bcl2SH G-quadruplex imino protons (∼10. 4-11. 5 ppm) showing chemical shift changes. No substantial chemical shifts were observed in the Watson-Crick base pair regions (∼12. 5-14 ppm) indicating that Znf706 binds exclusively to the G-quadruplex structures of the Bcl2SH sequence. All 1D NMR samples were prepared in 20 mM phosphate buffer containing 100 mM KCl and 7. 5% D2O (pH 7. 4).

### Znf706 preferentially binds to G-quadruplexes

To help identify Znf706’s binding preference, we pursued the possibility that Zn706, like the majority of C2H2 zinc fingers, interacts with nucleic acids. Extensive knowledge exists about the nucleic acid binding specificity of zinc finger proteins and their binding specificity can be predicted from the amino acid sequence alone, using the PWM predictor interactive position weight matrix tool (47). We began by validating the program writer’s claim that this program can successfully classify proteins containing known zinc fingers according to their DNA binding motifs, independent of whether they are G-rich or A and T-rich (Supplementary Figure S2). This was followed by inputting the amino acid sequence of Znf706 into the PWM predictor, and it gave a binding specificity for Znf706 of GGGG (Figure 1A). Fluorescence anisotropy showed that only polyG DNA binds to Znf706 in an experimental setting, while polyA, polyT, polyC, and polyN showed no detectable binding (Supplementary Figure S3A). Since polyG is known to readily form the non-canonical structures known as G-quadruplexes (48) we investigated if Znf706 binds to other known G-quadruplexes. We also evaluated sequences that were G-rich but not predicted to form G-quadruplexes (Supplementary Figure S3B) nor did so when tested experimentally by fluorescent dyes N-methylmesoporphyrin IX and thioflavin T (ThT) dyes known to bind G-quadruplexes (Table 1 and Supplementary Figures S3B and S4). We found that all the tested oligonucleotides known to form G-quadruplexes showed interaction with Znf706, and all the tested G-rich sequences known not to form G-quadruplexes, failed to bind Znf706.

To distinguish between G-quadruplex and duplex binding, we utilized the NMR spectral properties of G-quadruplexes. G-quadruplexes have a characteristic chemical shift region around ∼10. 5 and 11. 5 ppm where signals resonate from the G-tetrad forming guanine imino protons that are hydrogen bonded in the Hoogsteen geometry. These signals serve as a diagnostic G-quadruplex fingerprint since they are well separated from the imino proton’s signals, stabilized by Watson-Crick bonds, that resonate between ∼12. 5 and 14 ppm. When added to G-rich (GGA or GGGG) hairpin duplex structures, Znf706 was found to be unable to induce changes in proton chemical shifts. This indicates that Znf706 does not interact with these hairpins, even when both the G-rich hairpin and Znf706 are present at very high concentrations (Figure 1C and Supplementary Figure S5A-C). Next, to directly probe if Znf706 prefers to bind a G-quadruplex structure over a duplex formed from the same sequence and its complementary strand, a competitive NMR titration experiment was performed using a modified 15-nucleotide long sequence, as shown in Table 1, derived from a Bcl2 promotor referred to here as Bcl2SH that showed well-dispersed imino proton signals. This Bcl2SH sequence is capable of forming a parallel G-quadruplex either as a single-strand oligo, or a stable duplex, when in the presence of its complementary strand. Imino proton chemical shift analysis showed that Znf706 binds exclusively to the G-quadruplexes and fails to interact with duplex DNA, even at high micromolar concentrations. This indicates that Znf706 recognizes G-quadruplex structures and not solely the G-rich nature of the sequence (Figure 1D and Supplementary Figure S5D, E).

### G-quadruplex topology and folding influence Znf706 binding affinity

To further explore the substrate binding specificity of Znf706, we examined its binding capability to previously characterized G-quadruplexes. We tested oligonucleotides capable of forming G-quadruplexes derived from the promotor regions of the c-MYC (Pu19_A2A11) (49), BCL2, and c-KIT oncogenes, as well as the human telomeric repeat sequence (hTel) and TERRA (50) and a G4C2 repeat expansion sequence (51) associated with amyotrophic lateral sclerosis (ALS). All these G-quadruplexes have been extensively characterized (Table 1 and references therein). Using fluorescence anisotropy and microscale thermophoresis, we found that Znf706 binds to these G-quadruplexes with dissociation constants (K_d_) ranging from ∼ 1 to 20 µM. Znf706 bound strongly to Bcl2SH, TERRA, and cMyc, all of which form parallel type G-quadruplexes. Znf706 binds less strongly to hTel, dG4C2, and Bcl2WT sequences which form hybrid structures (52), and weaker still to Kit*, which forms antiparallel structures (53) (Figure 2A and Supplementary Figures S6 and S7A). These observations suggest that Znf706 prefers binding parallel over hybrid G-quadruplex structures. However, since parallel G-quadruplexes have previously been shown to be more tightly folded than hybrid structures (54), we cannot rule out the possibility that Znf706 prefers binding to tightly folded G-quadruplexes.

**Figure 2.**
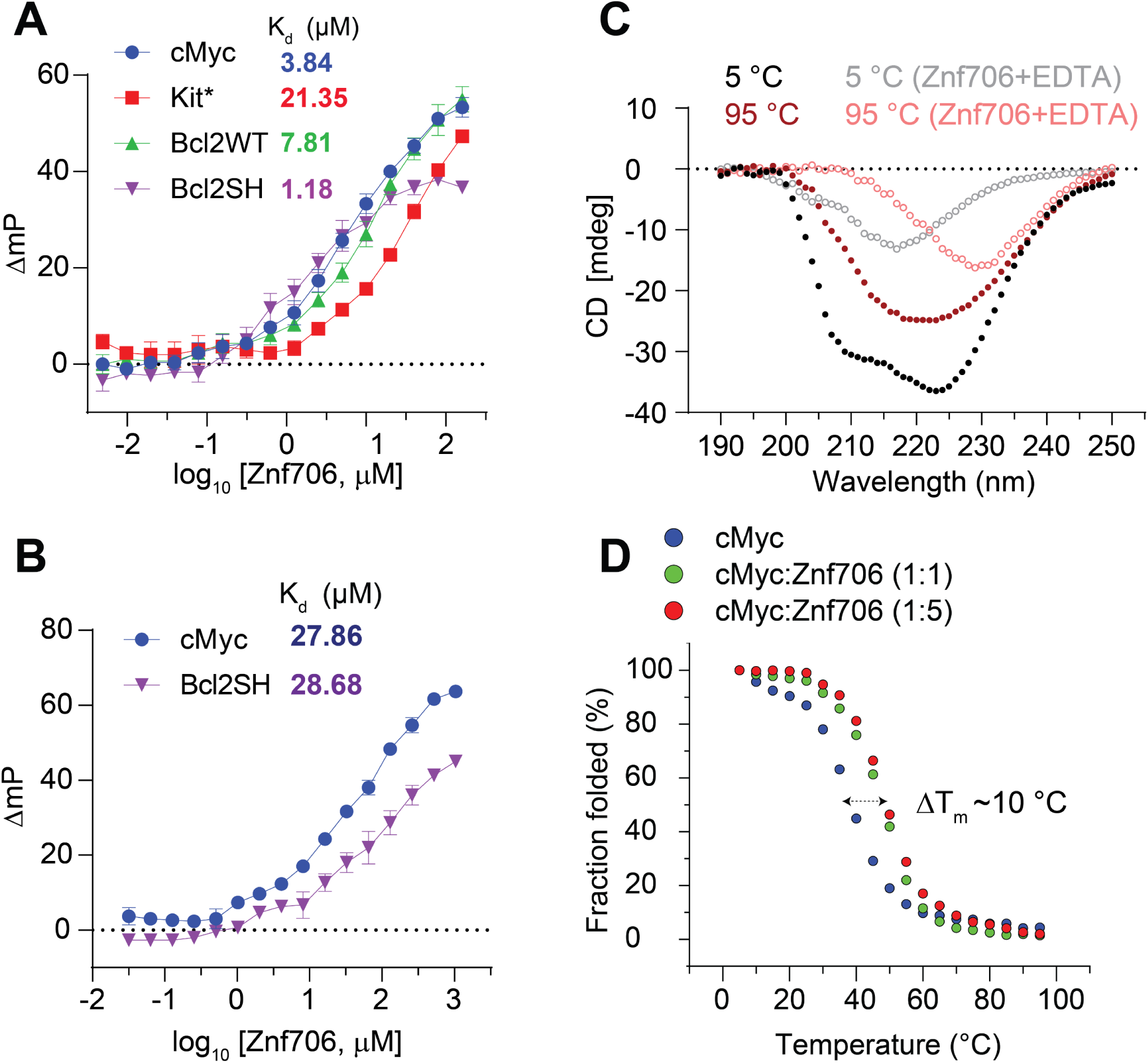
Znf706 displays thermal stability and binds more tightly to well folded G-quadruplexes. **(A-B)** FP binding assay with Znf706 and 5’ 6-FAM labeled G-quadruplexes prepared in 20 mM NaPi, 100 mM KCl, pH 7. 4 **(A)** or 20 mM Tris-HCl, 100 mM LiCl, pH 7. 4 **(B).** The indicated Kd values are calculated by non-linear regression analysis and one site binding saturation model in GraphPad Prism at an increasing concentration of Znf706 (4 nM to 130 µM for (**A**) and 32 nM to 1. 04 mM (**B**)). Error bars represent standard deviations derived from three replicates. **(C)** Secondary structure analysis of Znf706 (50 µM), in the absence (solid circles) or presence of 20×molar excess of EDTA (open circles), studied using Circular dichroism (CD) spectroscopy recorded at different temperatures as indicated. **(D)** CD melting curves of 20 µM cMyc G-quadruplex, in the absence (blue, Tm= 38. 67±0. 51 °C) or presence of equimolar (green, Tm= 47. 70±0. 52 °C) or 5×molar excess (red, Tm = 48. 88±0. 33 °C) of Znf706, dissolved in 20 mM NaPi, 4 mM KCl, pH 7. 4. The CD molar ellipticity at 264 nm, as a function of temperature, was normalized to generate the melting curves, and the CD melting temperatures (Tm) were calculated by fitting the fraction folding curves in the Origin program and the arrow indicates the change in Tm.

Consistent with the hypothesis that Znf706 interacts with G-quadruplexes in a structure-specific fashion, we observed a substantial increase in the K_d_ for parallel G-quadruplex forming oligonucleotides, such as Bcl2SH and cMyc (Table 1), when they are dissolved in buffer containing LiCl, which disfavors G-quadruplex formation as compared to when they are dissolved in a KCl solution, which favors G-quadruplex formation (Figure 2A, B). Similar observations were noticed in a four-repeat G4C2 G-quadruplex forming DNA sequence that folds better and binds stronger (K_d_ = 15. 05±2. 14) in KCl solution, as compared to in a NaCl (> 200 µM) solution (Supplementary Figure S7A, B). NMR experiments further showed that the G4C2 G-quadruplex forming oligonucleotide has a negligible binding effect on Znf706 ^1^H/^15^N chemical shifts even at a 2-molar excess concentration in the presence of NaCl as compared to KCl (Supplementary Figure S8). This is consistent with the knowledge that these oligos fold into more defined G-quadruplexes in KCl as compared to in NaCl (51). Conversely, the RNA G-quadruplex TERRA showing small structural differences by circular dichroism (CD) spectroscopy in KCl and LiCl solution presented a similar K_d_ for Znf706 (Supplementary Figure S7C, D).

### Znf706 binding stabilizes and can affect the topology of G-quadruplexes

It is known that protein binding can regulate the biological activity of G-quadruplexes by stabilizing or destabilizing them or affecting their topology (55). However, before investigating if Znf706 binding affects G-quadruplex folding, we investigated how Znf706 and the G-quadruplexes fold in isolation. Thermal denaturation of Znf706 by CD spectroscopy showed that it is highly thermostable. It slowly unfolds and folds in a highly reversible and non-cooperative manner sometimes seen in proteins containing multiple folding intermediates (56) (Supplementary Figure S9). This unusual heat resistance and folding behavior has previously been observed for the Znf706 homolog SERF2, which led to it being classified as a HERO, a “Heat Resistant protein of Obscure function” (57). Though Znf706’s temperature-dependent CD spectrum indicates loss of helicity upon shift to 95 °C, it does retain a substantial amount of its β-structural characteristics at this strongly denaturing temperature (Figure 2C and Supplementary Figure S9). For such a small disulfide-free protein, Znf706 is thus unexpectedly thermostable. The thermal stability of Znf706 was further confirmed by using 1D NMR, with spectra taken at temperatures ranging from 4 °C to 65. 5 °C (Supplementary Figure S10). These spectra continued to show well-dispersed, unchanged amide (N-H) peaks, in the range of ∼7 to 9. 2 ppm which correspond to a folded protein. Loss of structure, however, was observed when we treated Znf706 with the metal ion chelator EDTA, measured by both CD and NMR spectroscopy. Upon EDTA addition, the amide peaks narrowed into the 7 to 8 ppm range, indicating that metal chelation leads to the structural unfolding of its thermally stable, but metal ion-dependent, zinc-finger domain (Supplementary Figures S1D and S10). These findings support that the Znf706’s zinc finger plays a role and is important for Znf706 folding.

In isolation, all tested G-quadruplexes were found to unfold and refold reversibly with melting midpoints (T_m_) ranging from ∼37 to 53 °C, and ∼75 °C for Bcl2WT (Figure 2D and Supplementary Figure S11). These temperatures minimally impact Znf706 stability except for Bcl2WT, enabling us to determine if Znf706 impacts G4 folding, without being concerned that Znf706 is unfolding during the measurements. Through CD spectroscopy, we determined if Znf706 impacts G-quadruplex stability. CD results showed that for three of the four quadruplexes their T_m_ increased upon Znf706 binding (Figure 2D and Supplementary Figure S11). The high thermal stability of Znf706 also encouraged us to test the influence of Znf706 on the refolding pathway of G-quadruplexes, considering that G4 quadruplexes are thought to be metastable in cells (58, 59). We first tested if Znf706 binds G-quadruplexes at high temperatures and chose cMyc G-quadruplex to study because it shows well-dispersed imino signals. We observed cMyc binding induces ^15^N/^1^H chemical shifts in Znf706 at 50 °C and vice-versa. The binding at 50 °C is very similar to that observed at 37 °C (Figure 3A and Supplementary Figure S12). To understand if these binding events have any effect on G-quadruplex refolding pathways, the G-quadruplexes were denatured by heating to 95 °C, followed by rapid cooling, both in the presence and absence of Znf706. cMyc and Bcl2SH refolded very rapidly into parallel structures and Znf706 binding did not affect their refolding. However, the presence of Znf706 resulted in Bcl2WT and Kit* refolding into hybrid-like structures with an increase in antiparallel-like structures. This was evidenced by a decrease in the 260 nm CD peak intensities and an increase in the 295 nm CD peak intensities when compared to the spectra that were measured in the absence of Znf706 (Supplementary Figure S13). Therefore, we proposed that Znf706 is capable of not only stabilizing pre-folded G-quadruplexes but also capable of altering their folding behavior likely by intervening in their folding pathway.

**Figure 3.**
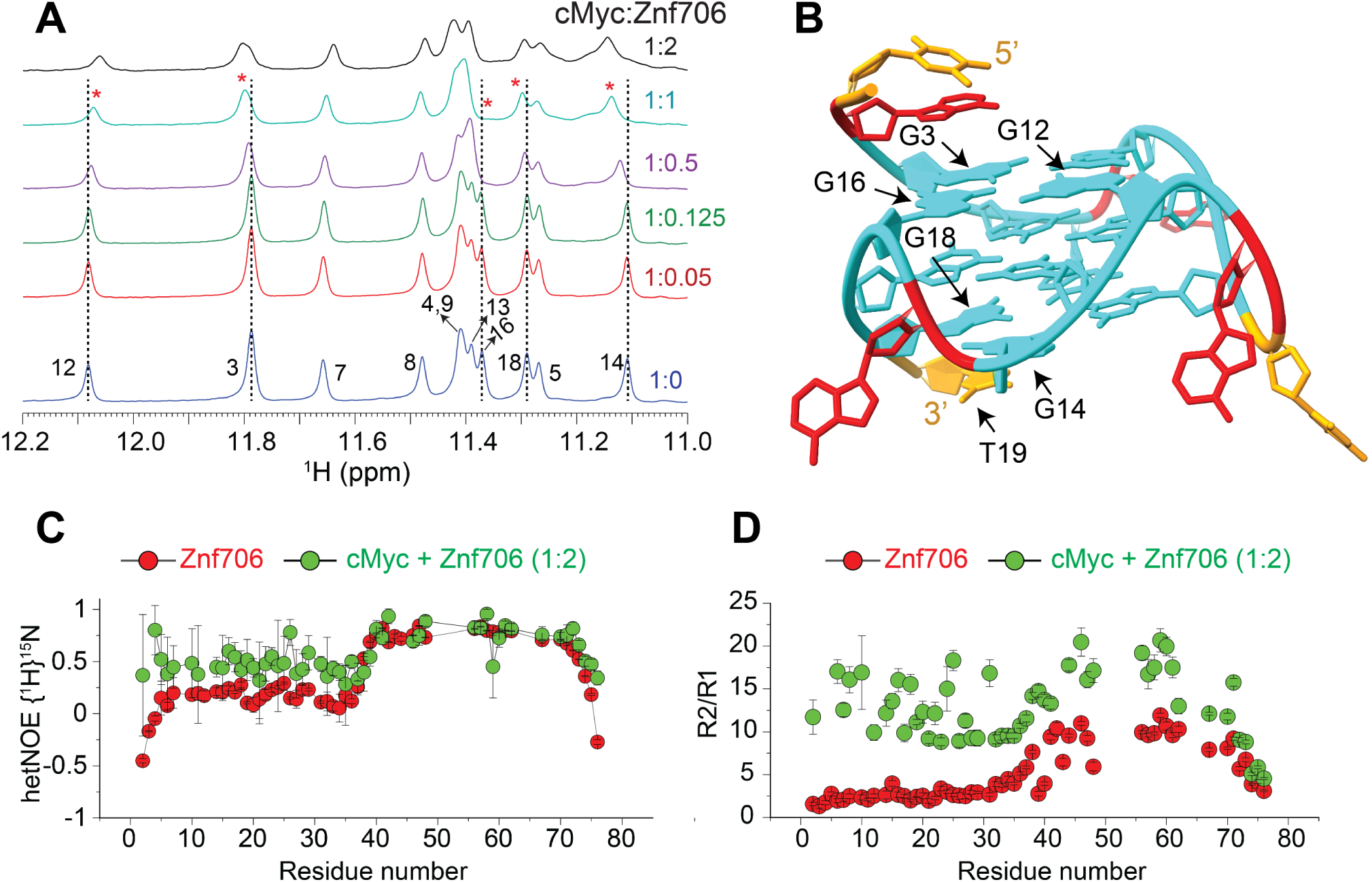
NMR-guided structural and dynamic studies showing cMyc G-quadruplex binding induces conformational rigidity in the N-terminal SERF domain of Znf706. **(A)** 1D proton NMR spectra recorded at 37 °C showing the imino proton chemical shifts in cMyc G-quadruplex (200 µM), in the absence (blue) and presence of an increasing concentration of Znf706 (10 – 400 µM) dissolved in 20 mM NaPi (pH 7. 4), 100 mM KCl. The G-imino peaks showing significant chemical shift changes at sub-stoichiometric protein concentration are marked with dashed lines and asterisks. **(B)** A high-resolution structure of cMyc (PDB ID: 2LBY) with the nucleotides likely involved in the binding with Znf706 and/or undergo conformational alteration upon Znf706 binding obtained from Figures 3A and S14 are indicated by arrows (G3, G12, G14, G16, G18, and T19). A, T, and G nucleotides are shown in red, orange, and cyan, respectively. **(C)** Heteronuclear NOE measurement of 200 µM ^15^N Znf706, in the absence (red) and presence (green) of 100 µM cMyc, showing the induction of structural rigidity in Znf706’s N-terminal upon cMyc binding. Standard errors are estimated from the signal-to-noise ratios. **(D)** R2/R1 relaxation ratios of 200 µM Znf706 in the absence (red) and presence (green) of 100 µM cMyc G-quadruplex. The heteronuclear NOE and relaxation NMR experiments were done on a Bruker 600 MHz at 4 °C in 20 mM NaPi, 100 mM KCl, pH 7. 4 buffer containing 7. 5% D2O.

### cMyc G-quadruplex binding induces conformational rigidity and rearrangement of Znf706

Despite the role that G-quadruplexes play in affecting gene regulation and protein aggregation, and the importance of protein binding in the formation and stabilization of G-quadruplexes in vivo, relatively little is known about how proteins interact with G-quadruplexes (26, 31) or the role that protein dynamics may play in controlling protein and G-quadruplex interactions (51, 60, 61). NMR titration measurements showed that cMyc interacts with Znf706 at 37 °C through nucleotides G3, G12, G14, G16, G18, and T19 (Figure 3A, B). Based on these results, the binding interface spans all three stacked G-quartets in cMyc (Figure 3B). The G-quartet serves as the fundamental structural unit in all G-quadruplexes (62). The Znf706 protein most likely interacts through the grooves of the G3 and G16 residues and forms a structural arrangement with the G5-G9-G14-G18 tetrad and the T19 residue on the 3’ end (Figure 3B and Supplementary Figure S14). NMR titration measurements also showed that like cMyc, when Znf706 binds to Bcl2WT G-quadruplexes (49, 63), substantial chemical shift changes occur in the signals of guanine imino protons that include G1, G2, G3, G17, G18, and G23 nucleotides distributed across all three stacked G-quartets (Supplementary Figure S15). This large discontinuous nature of this binding surface implies that either multiple Znf706 molecules are involved in G-quadruplex binding or that Znf706 binding induces conformational changes in cMyc and Bcl2WT. Since the CD analysis indicates that the addition of Znf706 induces a small structural change in folded cMyc and Bcl2WT G-quadruplexes, we judged the second possibility to be less likely. Conversely, the decrease in Znf706’s CD ellipticity near ∼218-222 nm, where cMyc and Bcl2WT have no CD absorption, may suggest a change in Znf706’s secondary structure (Supplementary Figure S16). However, such changes could also arise due to a change in rotational symmetry, optical activity, and 3D chirality of Znf706 in a G-quadruplex bound state. To further understand these structural properties, we predicted the secondary structure of the Znf706 complex with cMyc from its backbone chemical shifts Cα, Cβ, and CO using TALOS-N (35). The structure comparison of Znf706 complexed with cMyc G-quadruplex showed a minimal secondary structure change when compared with Znf706 alone as shown in Supplementary Figure S17A suggesting Znf706 remains unstructured at N-terminus during G-quadruplex interaction.

Our Znf706 NMR structural data favorably positioned us to investigate the effect that G-quadruplex binding has on Znf706’s structure and dynamics. To determine if G-quadruplex binding influences the dynamics of Znf706, we exploited the ^1^H/^15^N heteronuclear Overhauser Effect. This approach is often used to measure local backbone dynamics. Heteronuclear NOE values close to one indicate a high degree of order, while values less than or closer to zero indicate a high degree of disorder (64). Using this criterion, the N-terminal residues of Znf706 were identified to be disordered, and the zinc-finger region was identified to be ordered (Table S1). We decided to study the effect that parallel cMyc G-quadruplexes have on Znf706’s structure and dynamics at 4 °C where major of the ^15^N/^1^H cross peaks are detectable (Supplementary Figure S18). The heteronuclear NOE experiment of cMyc G-quadruplex binding was shown to substantially restrict the backbone motion of the N-terminal domain of Znf706 while leaving the zinc finger’s protein dynamics minimally perturbed (Figure 3C and Table S1). This change in disorder in Znf706 was further investigated by measuring the R2/R1 ratio of the spin-lattice (R1) and spin-spin (R2) relaxation rates (Figure 3D). This ratio measures the motion of individual residues, where lower values indicate fast motion and local flexibility, and higher values indicate slow motion and local rigidity. The N-terminal SERF homologous domain (residues 1-38) and C-terminal zinc finger domain (residues 39-76) of Znf706, in the absence of cMyc G-quadruplex, showed an average R2/R1 ratio of ∼2. 9 and ∼7. 8 respectively.

This is confirmation of their respective fast and slow dynamics (Table S1). Upon cMyc G-quadruplex binding, the motion in the N-terminal domain is greatly constrained as the average R2/R1 value increases from ∼2. 9 to ∼12. 3, and the zinc-finger domain is somewhat constrained as an increase in the R2/R1 ratio is seen from ∼7. 8 to ∼13. 9. This data indicates that both the N- and C-terminal domains of Znf706 may coordinate with one another in binding cMyc G-quadruplexes (Figure 3D). Heteronuclear NOEs that report faster time-scale motions, compared to R2/R1 which report slow time-scale motions, were able to capture the N-terminal domain motion but failed to capture the zinc-finger domain motion likely due to their slow motion in the cMyc G-quadruplex bound state.

This suggests in the Znf706-cMyc complex, the N-terminal disorder domain possesses a relatively fast dynamic motion required for G-quadruplex recognition as observed in other disorder proteins (65).

### Mapping G-quadruplex binding sites in Znf706

We continued our studies to identify the G-quadruplex binding sites on Znf706. ^1^H /^15^N 2D correlational NMR titration experiments allowed us to determine the specific residues of Znf706 that are involved in G-quadruplex binding. The addition of sub-stoichiometric quantities (5 µM) of G-quadruplexes showed that chemical shift perturbations predominantly occurred in many of the N-terminal, SERF-homologous regions of Znf706 in cMyc and Bcl2WT, but not in Kit* (Figures 4A, B and Supplementary Figure S17B). The residues A2, R3, K17, L37, and V43 were the most sensitive to G-quadruplex addition. Since these residues, containing varied N-terminal residues, were uniformly perturbed when mixed with 50 µM G-quadruplexes (DNA: Protein=1:2) independent of the G-quadruplexes added, it strongly suggests that Znf706 engages its N-terminal charged interface to bind different G-quadruplex structures, independent of their sequence or topology. Increasing the amount of G-quadruplex added, resulted in all the N-terminal residues having significant chemical shift perturbations, which additionally emphasizes the importance of this SERF homologous domain in G-quadruplex interaction. Furthermore, these increases in the G-quadruplex concentration to 50 µM also resulted in weak but noticeable chemical shift perturbations in the very C-terminal residues (70–75) of Znf706 (Figures 4C, D). These findings indicate either the N- and C-termini of Znf706 contribute to G-quadruplex binding or that a conformational change in Znf706 occurs upon G-quadruplex binding. Fluorescence polarization binding analysis of individual domains showed that a peptide consisting of just 35 N-terminal domain residues (residues 1-35) bound with K_d_ values ranging from ∼7 - 42 µM to the three tested G-quadruplexes (cMyc, Bcl2SH, and Bcl2WT). This is substantially weaker than their binding to full-length Znf706. The C-terminal zinc-finger domain showed very weak binding to all three G-quadruplexes, with K_d_ values too high to be accurately determined (> 200 µM) (Supplementary Figure S19A, B). Although the PWM predictor program indicates that Znf706’s C-terminal zinc finger interacts with polyG sequences, on its own the C-terminal zinc finger does not appear to be sufficient for high affinity interactions with G-quadruplexes.

**Figure 4.**
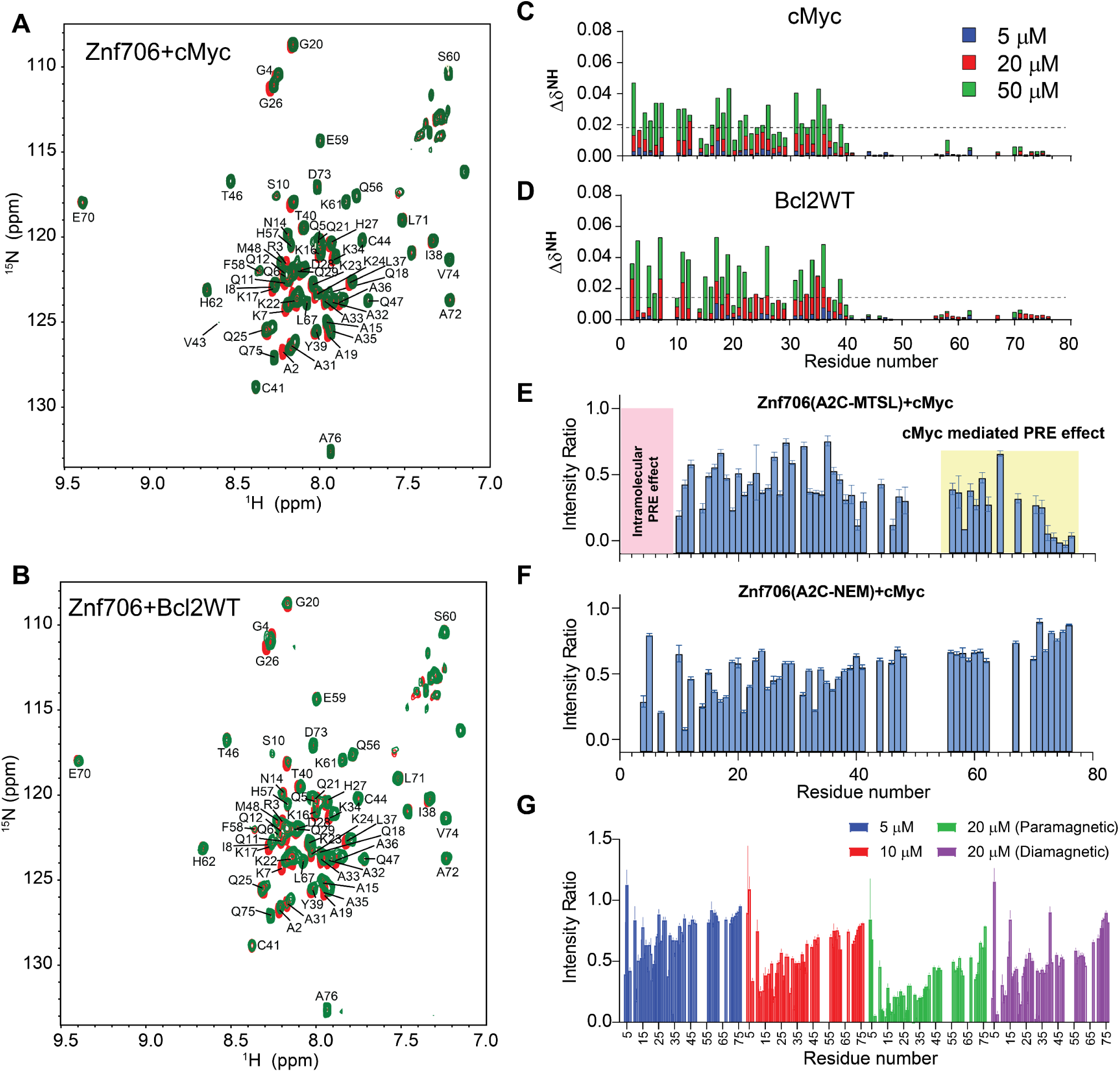
^15^N/^1^H heteronuclear and PRE NMR studies show that Znf706’s N-terminal SERF predominantly binds to G-quadruplex and its C-terminal zinc-finger domain facilitates complex formation. **(A-B)** ^15^N/^1^H 2D correlation spectrum of 100 µM uniformly ^15^N labeled Znf706 dissolved in 20 mM NaPi, 100 mM KCl, pH 7. 4 in the absence (red) and presence of 50 µM (green) cMyc **(A)** and Bcl2WT G-quadruplexes **(B**). The non-proline backbone amide resonances were assigned using a series of 3D NMR experiments that include HNCA, HNCACB, CBCA(CO)NH, HNCO, and HNCOCA and 2D ^15^N- and ^13^C-HSQC. **(C-D)** Plots showing chemical shift perturbations (CSPs) derived from the ^15^N/^1^H 2D spectrum of 100 µM Znf706 titrated with variable concentrations of cMyc **(C)** and Bcl2WT **(D)** G-quadruplexes as indicated in colors. The CSPs were calculated using the equation 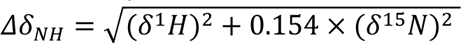 and the dashed lines indicate the average CSPs in each group. **(E-G)** Signal intensity ratio of amide protons observed for 100 µM ^15^N Znf706 A2C-MTSL **(E),** Znf706 A2C-NEM **(F)** mixed with 50 µM cMyc G-quadruplex. The intramolecular PRE effects at N-terminus are shaded in pink, whereas cMyc binding-induced PRE effects are highlighted in yellow. **(G)** Monitoring the paramagnetic effect upon cMyc binding to 50 µM ^15^N Znf706 A2C-MTSL at the indicated concentration. The final titrated product containing 50 µM Znf706 and 20 µM cMyc was reduced for ∼3 hours following the acquisition of the diamagnetic spectrum. NMR spectra were collected either on a Bruker 800 or 600 MHz spectrometer at 4 °C for samples dissolved in 20 mM NaPi, 100 mM KCl, pH 7. 4 buffer containing 7. 5% D2O. Standard errors were estimated from the signal-to-noise ratios.

To better understand the role of the C-terminal zinc-finger domain on substrate recognition, we performed NMR paramagnetic relaxation enhancement (PRE) experiments using two different Znf706 cysteine mutants (A2C and A76C) (Supplementary Figure S20). The high sensitivity of PRE NMR has often been used to measure intramolecular and intermolecular interactions (66). We hypothesize that if G-quadruplex binding is solely driven by the N-terminal SERF-homologous domain, the incorporation of a PRE tag, MTSL, into the N- terminus via an introduced cysteine mutant, A2C, should have no PRE effect on the C-terminus zinc-finger domain and vice versa. However, the addition of cMyc G-quadruplex to the PRE tagged A2C-MTSL Znf706 at a 2:1 Znf706^A2C-MTSL^:cMyc ratio did induce a strong PRE effect, locally as well as in several regions of the Znf706’s N- and C-terminus (Figure 4E). To rule out the possibility that A2C mutation and PRE tagging may be impacting the binding mode of Znf706 to the cMyc G-quadruplex, we tested the binding effect of the cMyc G-quadruplex on the same protein construct (Znf706 A2C) tagged with a non-PRE tag N-ethylmaleimide (NEM). This protein, measured under the same experimental conditions and at the same molar ratios i. e. 2:1 Znf706^A2C-NEM^:cMyc, showed very little change in the signal intensities of the C-terminal residues 70-75, in contrast to the PRE tagged version, which did have large signal intensity changes in these residues (Figure 4F). This result argues against the possibility that cMyc binding directly influences the C-terminal residues signal intensities. To further investigate this possibility in the context of Znf706^A2C-NEM^ binding to cMyc, we compared the relative change in signal intensities of corresponding residues in diamagnetic (reduced) and paramagnetic (oxidized) environments which correlate to the difference in the PRE effect in Znf706 and Znf706-cMyc complex systems (Figures 4G and Supplementary Figure S21A). First, a slow titration of cMyc to Znf706 A2C-MTSL was carried out to monitor the PRE effect on C-terminal residues 70-75. At 1:10 cMyc: Znf706^A2C-MTSL^ molar ratio, a substantial loss of signal intensity was observed only for the N-terminal residues. Upon increasing cMyc concentration (1:5 cMyc:Znf706), signal loss in the C-terminal residues 70-75 appeared which subsequently increased at a 1:2. 5 cMyc:Znf706 ratio (Figure 4G). Introduction of a diamagnetic environment recovered an average ∼35 % loss of signal intensities for residues 70-75 implying that C-terminal residues of Znf706^A2C-MTSL^ may be involved in cMyc complex formation (Figure 4G and Supplementary Figure S21A). However, since cMyc binding to Znf706 ^A2C-NEM^ did not show a significant decrease in intensity for C-terminus residues and the diamagnetic condition did restore signal loss of the C-terminal residues (Figure 4E-G), we concluded that the PRE induced change in signal intensities is not likely due to direct cMyc G-quadruplex binding. A more plausible explanation is that it is due to Znf706’s structural rearrangements or oligomerization that occurs upon G-quadruplex binding when the N- and C-termini are brought close together. Consistent with this interpretation, a reciprocally similar PRE effect on the N-terminus residues was observed when a PRE tag was placed at the C-terminus (A76C) as in Znf706^A76C-MTSL^ (Supplementary Figure S21B).

### Znf706 and G-quadruplex interactions mediate liquid-liquid phase transitions

Many nucleic acid binding proteins, in the presence of RNA or DNA, readily undergo LLPT in vitro and nucleic acids are important components of cellular LLPT (67). Proteins, like Znf706, containing low complexity regions are overrepresented in LLPT compartments and G-quadruplexes have recently been reported to accumulate in one of these compartments known as stress granules (24, 68, 69). Upon mixing Znf706 with G-quadruplexes, LLPT droplets formed and were found to contain both the protein and the DNA (Figures 5A-D, Supplementary Figures S22-S25, and Supplementary Videos SV1 and SV2). Droplet formation ceased at high salt concentrations (300 mM) which demonstrates the importance of electrostatic interactions in driving LLPT (Figure 5B). Droplets were formed in the presence of cMyc, Bcl2WT, and polyG G-quadruplex forming oligonucleotides. No droplets were observed in the presence of the Kit* G-quadruplex, nor polyA, polyT, polyC, and polyN (Supplementary Figure S26), which do not form G-quadruplexes. Interestingly, all these non-droplets forming oligonucleotides bind weakly or not at all to ZnF706. Fluorescence recovery after photobleaching (FRAP) experiments were carried out to investigate the liquid-like properties of these droplets. Results showed that Znf706 can diffuse within the Znf706 and G-quadruplex droplets after complexing with G-quadruplex DNA at half-lives varying from 80-180 seconds (Figure 5E-H). Similar FRAP recovery times are observed for proteins binding to anionic biomolecules such as nucleic acids and polyphosphates (70, 71). Bcl2WT and cMyc G4 sequences prepared in LiCl solution showed no droplets, but aggregate-like structures, when mixed with Znf706, suggesting G-quadruplex structures mediate LLPT (Supplementary Figure S27).

**Figure 5.**
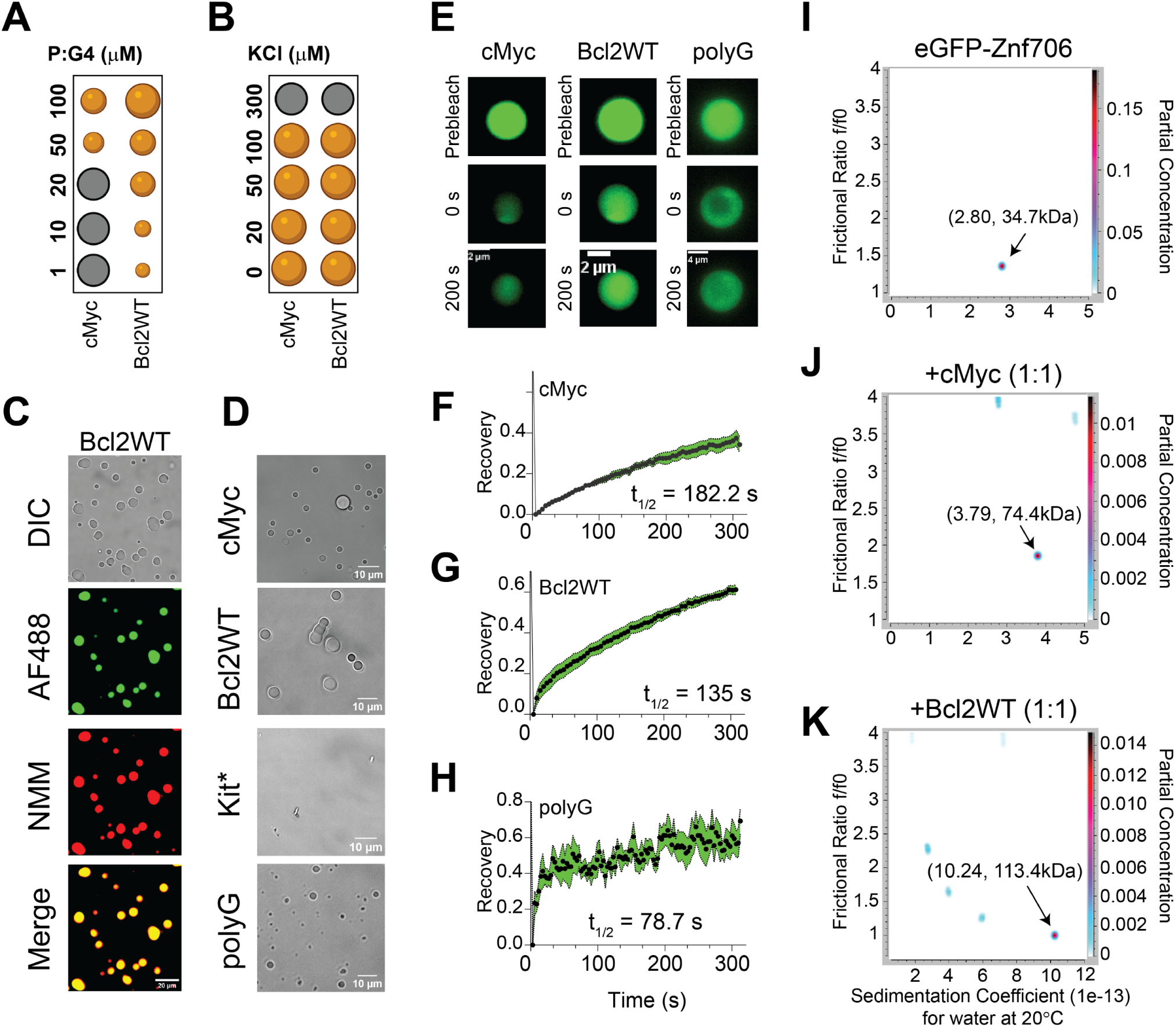
Znf706 binding to G-quadruplex forming DNA oligonucleotides induces liquid-liquid phase transition under physiological salt conditions. **(A-B)** Regime diagrams illustrating the effect of Znf706 and G-quadruplex (1:1) on droplet formation (black: no droplets, orange: droplets) at the indicated concentration **(A)** and variable salt concentrations with equimolar 100 µM Znf706 and G-quadruplex **(B)** on droplet formation. Sample solutions were incubated overnight before imaging. **(C)** Fluorescence images of Znf706-Bcl2WT droplets prepared in 20 mM NaPi, 20 mM KCl, pH 7. 4. The sample mixtures were prepared at room temperature by incubating 100 µM Znf706, 1 µM AF488-Znf706 (green signals), and 100 µM Bcl2WT. Droplet formation was confirmed by differential interference contrast (DIC) imaging, followed by the addition of 5 µM NMM to visualize Bcl2WT G-quadruplexes (red signals). **(D)** Monitoring droplet formation of 100 µM Znf706 mixed with an equimolar amount of topologically distinct G-quadruplexes as indicated. **(E)** FRAP analysis of Znf706 droplets formed in the presence of equimolar Znf706 and G-quadruplex reveals a long recovery suggesting slow Znf706 dynamics. Fluorescence images retrieved before FRAP, right after FRAP (0 sec), and at increasing diffusion times are shown. **(F-H)** Znf706 and G-quadruplex droplets were half-bleached and the normalized FRAP recovery plots are shown for Znf706 droplets formed in the presence of cMyc **(F),** Bcl2WT **(G),** and polyG **(H)**. G-quadruplexes were fitted to a one-phase association curve in GraphPad Prism to obtain the recovery half times (t1/2) which are indicated in the figures. Errors represent standard deviations of measurements derived from 4 droplets in each sample. **(I-K)** Determination of Znf706 and G-quadruplex complex size using analytical ultracentrifugation (AUC). eGFP-Znf706 (12. 2 µM, predicted molecular weight 35. 5 kDa) dissolved in 20 mM NaPi, 100 mM KCl, pH 7. 4 **(I)** was incubated with equimolar cMyc **(J)** or Bcl2WT **(K)** G-quadruplexes for ∼1 hour before AUC measurement (absorbance 488 nm). AUC data were analyzed in Ultrascan and a two-dimensional plot of frictional ratio f/f0 versus sedimentation coefficient ‘S’ are shown with an indicated estimated molecular weight as shown by arrows. The partial concentration represented by color intensity in the z-dimension represents the abundance of each species.

### Znf706 and G-quadruplex complexes are polymorphic

Multivalent interactions are recognized as being important for biomolecular condensate formation (72–74). Our 2D correlational NMR observations suggest that Znf706’s SERF homologous and zinc-finger domains may be directly and indirectly involved in G-quadruplex recognition as evidenced by strong and weak chemical shift perturbations, respectively (Figure 4C). The N-terminal residues 2-37 are predominantly involved in G-quadruplex binding, whereas the C-terminal residues 70-75 are spatially rearranged, likely stimulating complex formation as evidenced by the PRE analysis (Figure 4E). Since both domains are accompanied by a substantial PRE effect experienced by residues located in one terminus while in the presence of a PRE tag on the other terminus, it suggests that G-quadruplex binding may mediate structural rearrangement of Znf706 or its oligomerization (Figure 4F, G). In isolation, G-quadruplexes showed multiple signals when tested using AUC and size-exclusion chromatography, indicating the presence of multiple oligomer structures (Supplementary Figures S28 and S29). However, in the presence of Znf706, these oligomers resolved into a single higher molecular weight Znf706 containing species (Figures 5I-K and Supplementary Figure S28). Size calculations show that Znf706 in the presence of an equimolar amount of cMyc G-quadruplex, results in the formation of a 2:1 protein:cMyc complex. In the presence of the Bcl2WT G-quadruplex, larger more polymorphic complexes were detected (Figures 5I-K) indicating that G-quadruplex binding induces oligomerization. Our results obtained from NMR chemical shift, PRE NMR, domain-specific binding affinity and AUC experiments have enabled us to propose a model of Znf706 binding to cMyc G-quadruplex where two Znf706 molecules bind to opposite G-tetrad surfaces of the G-quadruplex mainly via Znf706’s charged N-terminus (Supplementary Figure S19C).

### Znf706 colocalizes with G-quadruplexes in vivo

To investigate if Znf706 and G-quadruplex interactions occur in vivo, we studied the colocalization of endogenously expressed Znf706 and DNA G-quadruplexes in human embryonic kidney (HEK) 293T cells. DNA G-quadruplexes are abundantly distributed in the nucleus of cells such as HeLa, U2OS, K562, HaCaT, hESCs, MEFs, and H1299 as visualized using different approaches (23, 68, 75–78). Znf706, stained in green using an anti-Znf706 antibody (Supplementary Figure S30A), and DNA G-quadruplexes, stained in red using the anti-DNA G-quadruplex antibody, were almost exclusively localized to the nucleus (Figure 6A). Colocalization was further confirmed using the intensity profiles obtained from the 2D single-cell image analysis (Figure 6B). This showed a significant signal intensity overlap between Znf706 in green and the anti-DNA G-quadruplex antibody clone 1H6 in red. The Pearson’s correlation coefficient between DNA G-quadruplexes and Znf706 staining was 0. 43± 0. 13, very similar to the correlation coefficients previously observed between anti-DNA G-quadruplex staining antibodies and known DNA G-quadruplex binding proteins such as TRF1, acetylated H3K9, demethylated H3K9, PML bodies, and RNA polymerase II. All of these have in vivo Pearson’s correlation coefficients with G-quadruplex antibodies ranging from 0. 15 to 0. 68 (Figure 6B and Supplementary Figure S30B) (75). Thus, we conclude that Znf706 likely colocalizes with G-quadruplexes in vivo.

**Figure 6.**
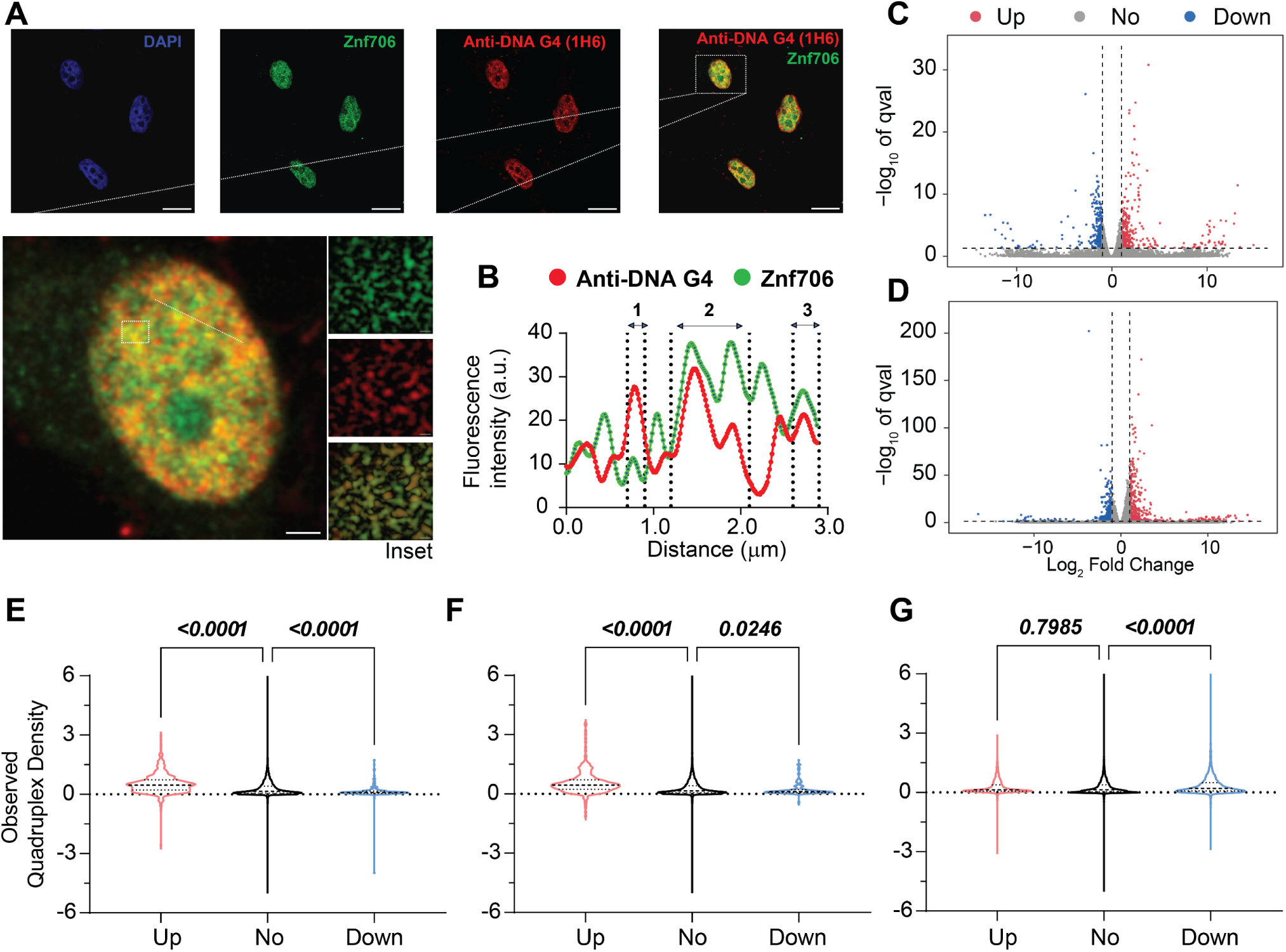
Znf706 co-localizes with DNA G-quadruplexes and its depletion alters gene expression in vivo. **(A)** Formaldehyde fixed HEK 293T cells were immunolabeled and counterstained with DAPI, anti-Znf706, and anti-DNA G-quadruplex binding specific antibodies, clone 1H6, at the indicated colors. A zoomed single cell, with an inset highlighting the Znf706 and DNA G-quadruplex colocalization (yellow signals), is shown. **(B)** The degree of Znf706 and G-quadruplex colocalization was calculated by quantifying the red and green fluorescence signal intensities across a straight line drawn along a selected cell, as shown in zoom using ImageJ. The scale bar represents 10 μm. (**C-D**) Volcano plots of RNA-seq differential expression analysis in Znf706 knockdown HeLa **(C)** and HEK 293T **(D)** cells, compared to a control knock-down. The genes with a minimal fold change in gene expression levels are represented within the dashed vertical lines. Znf706 depletion down- and up-regulates 458 and 696 genes in HeLa, and 294 and 405 genes in HEK 293T cells, respectively, and are indicated on the left and right side in the volcano plots. **(E-F)** Violin plots of observed G-quadruplex density of annotated genes from Znf706 knockdown RNA-seq in HeLa **(E)** and in HEK 293T **(F)** cells. A similar analysis was done to generate a violin plot for the relationship between G-quadruplex density and genes affected by a DHX36 knockout in HEK 293 cells **(G)**, using the data obtained from work done by Chambers et al (21).

### G-quadruplexes suppress the effects of Znf706 on protein aggregation

SERF proteins were originally isolated by their ability to accelerate the aggregation of amyloid-prone proteins such as α-Synuclein, which has been neuropathologically linked to Parkinson’s disease (79). Interestingly, G-quadruplexes have recently been reported to possess very potent anti-aggregation activity (43) and remodeling of G-quadruplex structures by α-Synuclein has also been demonstrated in vitro (80). These findings led us to investigate whether the previously reported pro-amyloid formation of SERF might be related to our demonstration that the SERF homolog Znf706 can interact with G-quadruplexes. We started by demonstrating that Znf706 can accelerate α-Synuclein aggregation effectively, even at sub-stoichiometric concentrations (Figure 7A and Supplementary Figure S31A, B). This was monitored using ThT fluorescence, a dye whose fluorescence changes upon binding to amyloid fibers (81). Determining the effect of G-quadruplexes on α-Synuclein amyloid formation using this assay, however, is complicated by the observation that ThT also fluoresces upon binding to G-quadruplexes (Supplementary Figure S4). To get around this issue, we used light scattering and electron microscopy to monitor α-Synuclein amyloid formation. The addition of G-quadruplexes suppressed the ability of Znf706 to accelerate α-Synuclein aggregation (Figure 7B), presumably by competing for binding. Transmission electron microscopy (TEM) imaging revealed that 100 µM α-Synuclein, when mixed with equimolar Znf706 or cMyc G-quadruplexes and incubated for ∼6 hours, forms elongated fibrils, as compared to those fibrils that just contain α-Synuclein (Figure 7C). Abundant amorphous aggregates (and a few visible fibers) were generated when α-Synuclein was mixed with an equimolar concentration of both Znf706 and cMyc G-quadruplexes (Figure 7C). TEM images taken at a longer time-point (∼ 48 hours) using a physiological α-Synuclein (50 µM) concentration at varied Znf706 and G-quadruplex ratios gave similar results (Figure 7D).

**Figure 7.**
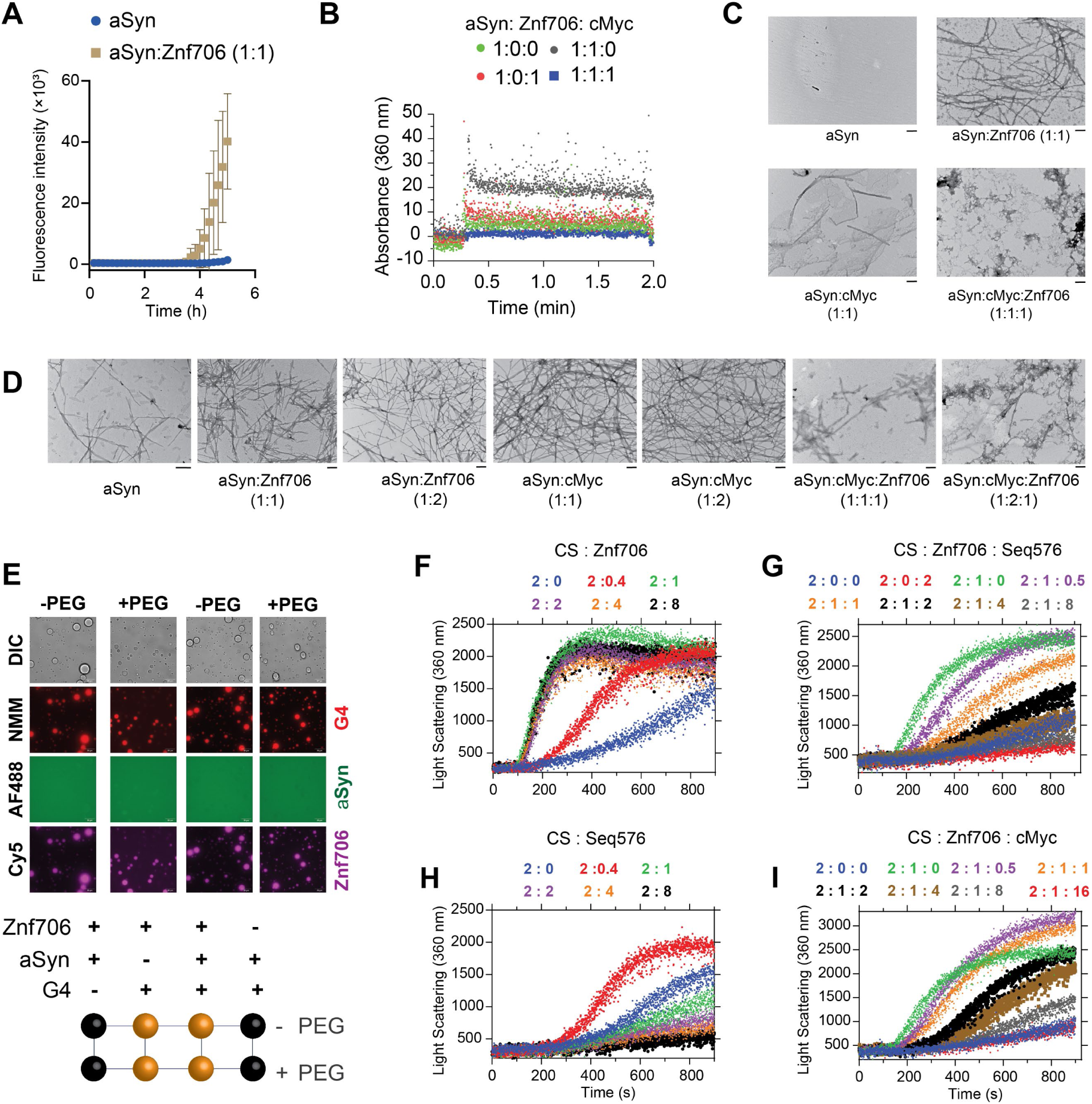
G-quadruplex binding suppresses the aggregation activity of Znf706. **(A)** Monitoring the aggregation kinetics of 100 µM α-Synuclein (aSyn) dissolved in 20 mM NaPi, 100 mM KCl, pH 7. 4 mixed, without (blue) or with 100 µM Znf706 (gold), by ThT fluorescence. Aggregation of aSyn in a sample containing equimolar Znf706 and cMyc G-quadruplexes is not shown and carried out in the absence of ThT. **(B)** Light scattering of aSyn aggregates (10 µL of aSyn aggregates dissolved in 1590 µL buffer) taken at time ∼6 hours at the indicated Znf706 and cMyc concentrations. **(C)** TEM images of aSyn aggregates (10 µL) taken at time ∼6 hours mixed with or without Znf706 and cMyc G-quadruplex as indicated. **(D)** TEM images of 50 µM aSyn aggregates taken at time ∼48 hours (see methods) mixed with the indicated concentration of Znf706 and cMyc G-quadruplexes. The scale bar is 200 nm. **(E)** Evaluation of the effect of aSyn (100 µM) on Znf706 and G-quadruplex LLPT in 20 mM NaPi, 100 mM KCl, pH 7. 4 without crowding agents (-PEG) and 20 mM NaPi, 150 mM KCl, 10% PEG8000, pH 7. 4 (+PEG). For fluorescent imaging, 0. 5 µM of AF488 labeled aSyn A90C and Cy5 labeled Znf706 A2C are mixed with 100 µM of unlabeled proteins. Droplet formation was confirmed by DIC imaging followed by the addition of 5 µM NMM, to visualize G-quadruplexes (red signals). A phase diagram for aSyn, Znf706, and G-quadruplex mixtures in ±PEG8000 is shown in the lower panel (see also DIC images in Supporting Figure 32). **(F-I)** Thermal aggregation of 300 nM citrate synthase (CS) was monitored using light-scattering mixed without (blue) or with an increasing concentration of Znf706 **(F**) or Seq576 G-quadruplexes **(H)** as indicated. A competitive thermal aggregation assay presenting the effect of an increasing concentration of Seq576 **(G)** and cMyc **(I)** G-quadruplexes on 300 nM CS mixed with 150 nM of Znf706. The thermal aggregation experiments were performed at 48 °C in 40 mM HEPES-KOH, pH 7. 5.

Further, we found that Znf706 mixed with Bcl2WT or cMyc G-quadruplexes readily forms droplets under physiological salt concentrations, in both crowded and non-crowded conditions (Figure 7E). However, α-Synuclein mixed with Znf706, or G-quadruplexes, does not undergo droplet formation under the tested conditions and showed no effect on Znf706 and G-quadruplex phase separation (Supplementary Figure S32). Three-component fluorescence analysis revealed the liquid droplets contain Znf706 and G-quadruplexes, but not α-Synuclein (Figure 7E). Using fluorescence polarization, we determined the relative affinity of Znf706 for G-quadruplexes and α-Synuclein in the same buffer conditions. Results showed that Znf706 has a very weak binding affinity for α-Synuclein as compared to G-quadruplexes (Figure 2A and Supplementary Figure S31C). These results support the idea that Znf706’s physiological function is more likely to be centered on G-quadruplex binding rather than on direct interactions with amyloid-prone proteins like α-Synuclein. In a condition where α-Synuclein undergoes phase separation (100 µM α-Synuclein in 20% PEG8000), we observed a liquid droplet to gel-like structure transition for α-Synuclein mixed with cMyc G-quadruplexes. Our results suggest that cMyc in isolation appears to promote α-Synuclein fibrillation (Figure 7D), and Znf706 binding suppresses this action (Supplementary Figure S33).

Our observations suggest that both Znf706 and G-quadruplexes affect not just α-Synuclein amyloid formation but also amorphous aggregate formation. We postulate that Znf706’s stronger affinity for G-quadruplexes over α-Synuclein suppresses Znf706’s ability to facilitate amyloid formation. Understanding that α-Synuclein can form both amyloids and amorphous aggregates and that Znf706 and G-quadruplexes can affect both processes, we decided to further investigate the effects of these macromolecules on amorphous aggregation using the classic chaperone substrate citrate synthase (82). This was inspired by the recent observation that G-quadruplexes are extremely efficient anti-aggregation agents in vivo and in vitro (43, 83). Unexpectedly, Znf706 effectively promoted citrate synthase amorphous aggregation, hinting that its molecular function in driving protein aggregation may not be limited to amyloid-prone proteins (Figure 7F and Supplementary Figure S34). Conversely, the G-quadruplexes that included Seq576, cMyc, and Bcl2WT (Table 1), all suppressed citrate synthase aggregation in the effective order of Seq576>cMyc≥Bcl2WT (Figure 7G-I and Supplementary Figure S35). Due to Seq576 having demonstrated the largest anti-aggregation effect (Table 1) from a large screening of DNA sequences (43), its efficacy as an anti-aggregation agent is expected. We next wondered if the chaperone function of G-quadruplexes is altered upon Znf706 binding. We found that the Seq576 G-quadruplexes reduced the citrate synthase aggregation that is promoted by Znf706, when present at super-stoichiometric concentrations (Figure 7H). In comparison to Seq576, the cMyc, and Bcl2WT G-quadruplexes showed relatively weaker activity in suppressing Znf706’s ability to facilitate citrate synthase aggregation. Therefore, we conclude that Znf706 can promote both amyloid formation and amorphous aggregation, while G-quadruplexes work in opposition to suppress both types of aggregation processes.

Given Znf706’s ability to influence G-quadruplex folding and the proposed role of these non-canonical structures in regulating gene expression, we wondered if Znf706 knockdowns impose any effect on gene expression in vivo and if this is in any way related to the presence or absence of G-quadruplexes within genes. Using RNA-seq, we found that a knockdown of Znf706 in two different cell lines results in the significant upregulation and downregulation of hundreds of genes. In HeLa cells, 696 annotated genes were > log2 (fold change) upregulated and 458 genes were downregulated (Figure 6C), while in HEK293T cells, an overlapping set of 405 genes were upregulated and 294 genes were downregulated (Figure 6D). Interestingly, these differentially expressed annotated genes correlate positively with previously observed quadruplex density (21). Those genes upregulated in Znf706 knockdown contain a significantly higher quadruplex density in their mRNAs, as compared to genes whose expression was not significantly changed. In contrast, genes downregulated in Znf706 knockdowns contain significantly fewer G-quadruplexes in their mRNAs or promoter regions than either the upregulated genes or the unchanged genes (Figure 6E, F). The depletion of DHX36, a known G-quadruplex unwinding helicase, shows a very different correlation between gene expression and quadruplex density (Figure 6G). In DHX36 knockout HEK293 cells, downregulated genes possess a significantly higher quadruplex density (21). Also known is that DHX36 can unwind G-quadruplexes in vitro (26), and therefore may affect gene expression effects through the regulation of G-quadruplex formation in 3′ UTRs and mRNA stabilization. We have shown that Znf706, by binding G-quadruplexes, can mediate G-quadruplex stability or folding in vitro. Znf706 may be mediating the observed changes in gene expression in vivo by G-quadruplex binding events that result in changes at the transcriptional level or changes in mRNA stability.

## Discussion

G-quadruplexes appear to play important roles in regulating biological processes (84) but how proteins bind these noncanonical structures and the impact this binding has on G-quadruplex structure and function remains under investigation (85). Uncovering the roles of G-quadruplexes is difficult because of the transient and dynamic nature of their formation in vivo (58). They can switch between quadruplex structures, non-quadruplex structures, and different G-quadruplex topologies (e. g., parallel, antiparallel, and hybrid). Protein binding significantly influences G-quadruplex folding and structure and thus impacts biological function (80, 85).

We have shown here that Znf706 specifically recognizes G-quadruplex structures, and that binding affects the structure and dynamics of both binding partners. The human Znf706 protein recognizes different structural motifs of G-quadruplexes including groove regions and planar surfaces, known as G-quartets (62), which are exposed on the top and bottom of a typical G-quadruplex structure. Znf706 preferentially binds G-quadruplexes with parallel or hybrid topologies, though during G-quadruplex refolding, Znf706 binding can alter the G-quadruplex folding resulting in different topologies. Recently, several other proteins have been identified that bind to G-quadruplexes and once bound, assist in regulating their folding, stability, and biological function. For example, nucleolin (60), FUS (51), helicases (27, 86), and G3BP1 (87) are known to bind to G-quadruplexes and regulate their functions by stabilizing or destabilizing the G-quadruplex structures, driving phase transition (88, 89), altering their cellular localization, or through inducing stress granule formation.

It is assumed that G-quadruplex binding proteins have the ability to recognize and modulate G-quadruplex structures in vivo and thereby regulate their biological processes. Thus, studying protein-G-quadruplex interaction is an important but challenging task (90). Our high-resolution NMR results show the involvement of the Znf706’s dynamic N-terminal domain in G-quadruplex binding (Figure 4) in a topology-dependent manner. This highlights the importance of this evolutionarily conserved domain in regulating the biological function of G-quadruplexes. Znf706 residues involved in G-quadruplex interaction map mainly to the low-complexity, highly disordered, N-terminal domain of Znf706, and G-quadruplex binding enhances dynamic stability in this disordered N-terminal region. This adds to the growing evidence that G-quadruplexes can bind with disordered protein domains (91, 92). Znf706 undergoes major structural rearrangements upon binding and multiple molecules can bind to a single G-quadruplex (Figures 4E, G and 5J, K). By displaying a limited and distinct number of binding sites, possessing only a moderate degree of disorder, and being biophysically well-behaved, Znf706 and the model G-quadruplexes that it interacts with appear to present a good opportunity to understand the liquid-liquid phase transition in biophysical detail. This has advantages over the study of larger, more disordered proteins which interact with each other, and their nucleic acid partners, in a less ordered and more multivalent fashion.

Znf706 binding to G-quadruplexes was shown to drive LLPT (Figure 5), a biological process increasingly being recognized as important for membraneless intracellular organization and gene expression (93, 94). Other recent studies have also linked protein binding to G-quadruplexes with phase transition (51, 95), for example, G-quadruplexes can trigger phase transition and stress granule formation in the cytoplasm (88, 96–98) and nuclear condensates such as histone protein condensation (99–101). Recent studies have established a relationship between G-quadruplexes and stress granule formation (102), implying that it may well be worthwhile to further study the relationship between G-quadruplexes’ cellular stress response and their possible association with human diseases.

Though DNA and RNA G-quadruplexes are structurally similar, both consisting of a series of G-tetrad stacks, RNA G-quadruplexes are more often observed with parallel topology than DNA G-quadruplexes. Znf706 specifically binds to parallel DNA G-quadruplexes and also to the RNA G-quadruplex TERRA indicating it binds to both RNA and DNA G4-quadruplexes. In ALS, the DNA form of the (GGGGCC)n repeat expansion forms droplets similar to the RNA droplets identified in cellular ALS aggregates that have been linked to disease phenotypes (103). DNA G-quadruplexes have been shown to mimic the macroscopic phenotype of RNA G-quadruplexes, and the FUS protein binding to G-quadruplexes has been linked to the suppression of neurodegeneration (104). Though Znf706 has been characterized as a transcription repressor, its ability to bind to both DNA and RNA G-quadruplexes with similar affinities encourages broader investigation of its in vivo activities.

Liquid-liquid phase transitions are known to nucleate and modulate the rate kinetics of α-Synuclein aggregation (79, 105), a process that initially involves liquid droplet formation of misfolded proteins followed by their maturation into more amyloid-like gels (105, 106). We found that Znf706 and G-quadruplexes form liquid-liquid phase transitions that are dynamic, undergo fusion, possess fluid-like reversible behavior, and are formed using multivalent interactions (Figure 5). Biophysical approaches including analytical ultracentrifugation, and NMR techniques such as homo- and hetero-nuclear multidimensional experiments, paramagnetic relaxation enhancement (PRE), and relaxation NMR studies proved amenable to studying the structures and dynamics of the Znf706 and G-quadruplex complexes. This allows for the rare possibility of obtaining detailed insights into how protein and nucleic acid interactions affect the structure and dynamics of the binding partners and how their interaction modulates LLPT. A plethora of evidence highlights the role of LLPT in exacerbating neuropathophysiology (107) and G-quadruplexes have been linked to many folding diseases (33). G-quadruplex forming sequences are enriched in Alzheimer’s aggregates (108) and several aggregation-prone proteins underlying ALS including hnRNP family proteins (109), FUS (110), TDP-43 (111), Ewing’s sarcoma protein (112), and TIA1, are shown to bind G-quadruplexes (33). Additionally, the FMRP protein, whose loss or mutation is linked to another folding disease, Fragile X syndrome, also binds G-quadruplex structures (113).

Znf706’s N-terminus is homologous to SERF, a protein that was originally discovered due to its ability to accelerate amyloidogenesis in disease models (15). Given its homology to SERF, it is not surprising that Znf706 also accelerates α-Synuclein fibrillation. Somewhat more surprising is our finding that Znf706 also accelerates citrate synthase amorphous aggregation (Figure 7). This observation suggests that the SERF family of proteins may have a broader role in regulating protein folding than previously appreciated (12, 16). Our observation that Znf706 binds specifically to G-quadruplex forming sequences in combination with the recent observation that G-quadruplexes serve as potent molecular chaperones (43), led us to investigate the relationship between Znf706’s ability to accelerate protein aggregation and G-quadruplexes ability to inhibit that aggregation. We find that G-quadruplexes suppress Znf706’s ability to promote both α-Synuclein fibril maturation and citrate synthase amorphous aggregate formation (Figure 7), and Znf706 suppresses G-quadruplex chaperone activity. We thus can propose a model where Znf706 and G-quadruplexes affect protein folding but in opposite ways. Znf706 knockdowns affect mRNA levels in a way that is related to the presence or absence of G-quadruplexes in the affected genes (Figure 6). That Znf706 can affect G-quadruplex folding and stability in vitro raises the possibility that this could be the mechanism by which it regulates mRNA levels in vivo. Whether this is due to G-quadruplexes acting on transcription or mRNA stability remains to be shown. Our results suggest that Znf706 may play a role in the control of gene expression through its ability to bind G-quadruplexes and to competing roles of the SERF family proteins and G-quadruplexes in controlling protein aggregation.

### Author Contributions

B. R. S. and J. C. A. B. conceived the original idea; J. C. A. B. and J. P. supervised the research; B. R. S., N. W. C., X. D., and E. W. performed the experiments in this study. B. R. S., E. W., A. K., B. B. G., N. M., H. Y., and D. D., performed data analysis. B. R. S., V. K., and J. C. A. B. wrote the manuscript. The manuscript was written through the contributions of all authors. All authors have approved the final version of the manuscript.

## Acknowledgments

We thank Dr. Anthony Vecchiarelli and Joseph Basalla for providing an instrumentation facility and help with fluorescence imaging.

## Funding Sources

J. C. A. B. is funded by the Howard Hughes Medical Institute. D. D. is funded by a UNC start-up and a grant from the National Institutes of Health, R35GM142864. J. P. and V. K. acknowledge financial support from the Slovenian Research Agency [grants P1-0242, Z1-3192, and J1-1704].

## Notes

### Competing Interest Statement

The authors have declared no competing interest.

### Summary of Updates

This version of the manuscript has been revised with several additional experiments and major of these data are included in the supporting information. This version contains an updated methods section and also includes some of the experimental method details that were missing in the previous version.

