## Supplemental File for "Protein G-quadruplex interactions and their effects on phase transitions and protein aggregation"

**Table of Contents**

**Supplementary Table S1** ………………………………………………………. 2

**Supplementary Figures S1-35**…………………..………………………..……..…3-37

**Supplementary videos SV1-SV2**…………… ……………………………….…......37

**Supplementary Table S1.** Summary of relaxation rates and heteronuclear NOE for Znf706 N- and C-terminal domains.

| **Znf706 (residues)** | **$R_1$ ($S^{-1}$)** | **$R_2$ ($S^{-1}$)** | **hetNOE** |
|---|---|---|---|
| Znf706 (1-38) | 1.61±0.10 | 4.71±0.07 | 0.17±0.06 |
| Znf706 (39-76) | 1.51±0.10 | 11.83±3.91 | 0.67±0.17 |
| Znf706 (1-38) + cMyc | 1.35±0.07 | 16.70±4.25 | 0.46±0.11 |
| Znf706 (39-76) + cMyc | 1.28±0.10 | 17.82±5.49 | 0.68±0.19 |

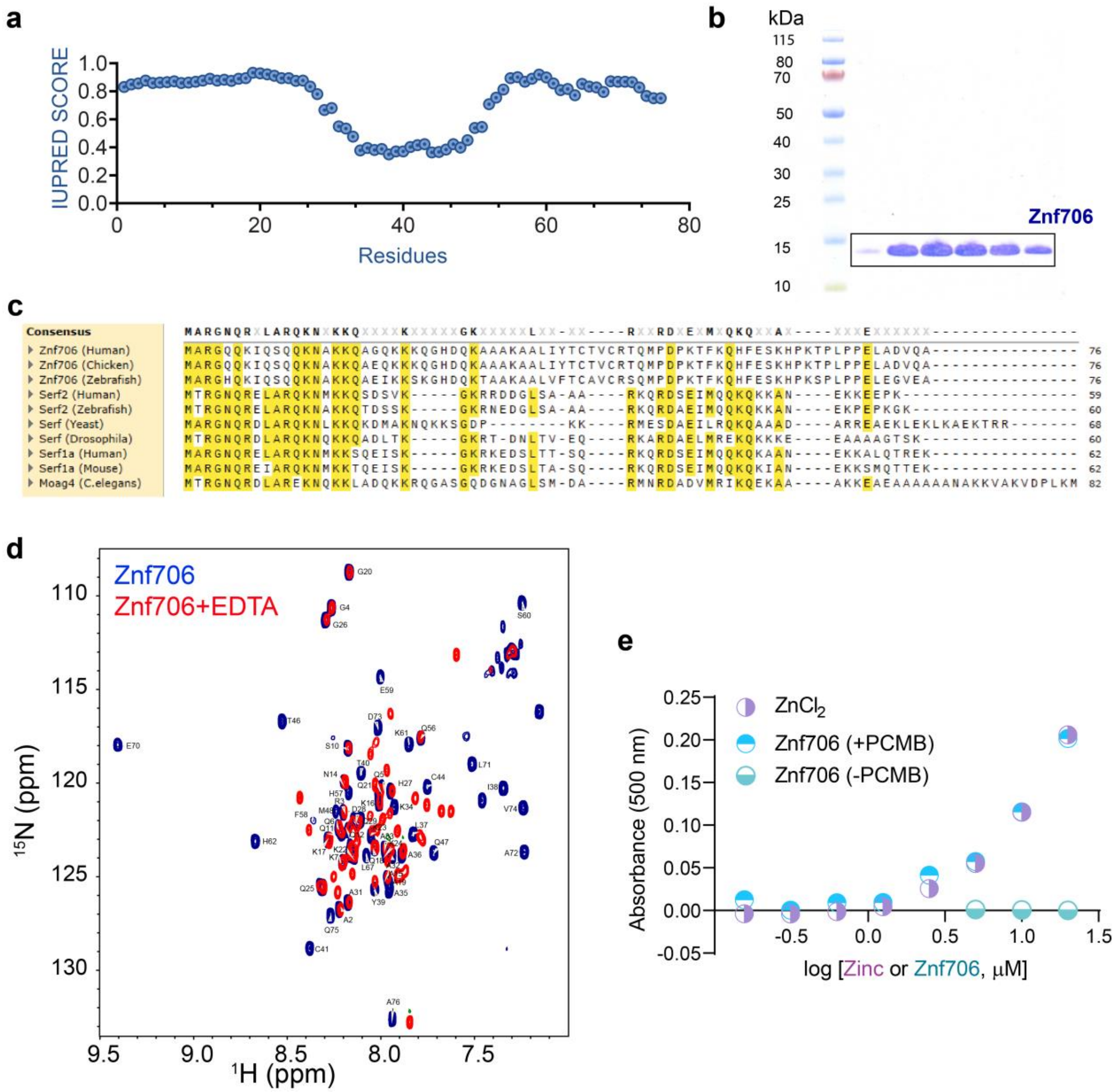


**Figure S1**. (**a**) Prediction of the disordered regions in Znf706 using IUPred2A, with disordered regions indicated by a high IUPred score. (**b)** Recombinantly expressed and purified human Znf706 as analyzed by SDS-PAGE and Coomassie Blue staining, box indicates the pooled fractions. **(c)** Sequence alignment of human Znf706 with different SERF proteins generated using Clustal Omega. Identities are highlighted in yellow, and the conserved residues are shown on the top. **(d)** $^{15}N/^{1}H$ TROSY experiments of 50 µM Znf706 dissolved in 20 mM NaPi, 100 mM KCl, 0.05% $NaN_3$, pH 7.4 in the absence (blue) or presence of 100x molar excess of EDTA (red) recorded at 4 °C. **(e)** Estimating the concentration of zinc in Znf706 by 4-(2-pyridylazo) resorcinol (PAR) assay. Zinc coordinated cysteines in Znf706 were reduced by para-chloromercuribenzoic acid (PCMB) and compared with Znf706 under a non-reducing condition, without PCMB (-PCMB).

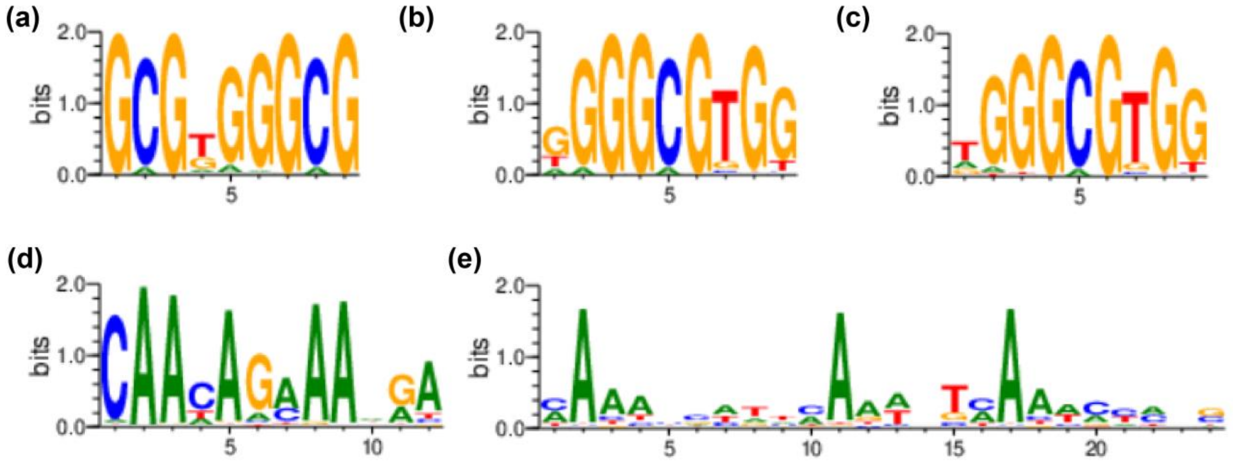


**Figure S2**. Prediction of five C2H2 type zinc-finger proteins that are known to bind to G-rich sequence motifs that include (a) Zif268 (UniProt: Q8MJM9), (b) Sp-1 (UniProt: P08047), and (c) Krueppel-like factor 3 or BKLF (UniProt: P57682), (d) yeast AZF1 (UniProt: P41696) which is known to bind AAAAGAAA (d), and (e) human SALL4 (UniProt: Q9UJQ4) that binds to AT-rich motifs (e) using the Interactive PWM predictor (http://zf.princeton.edu/logoMain.php).

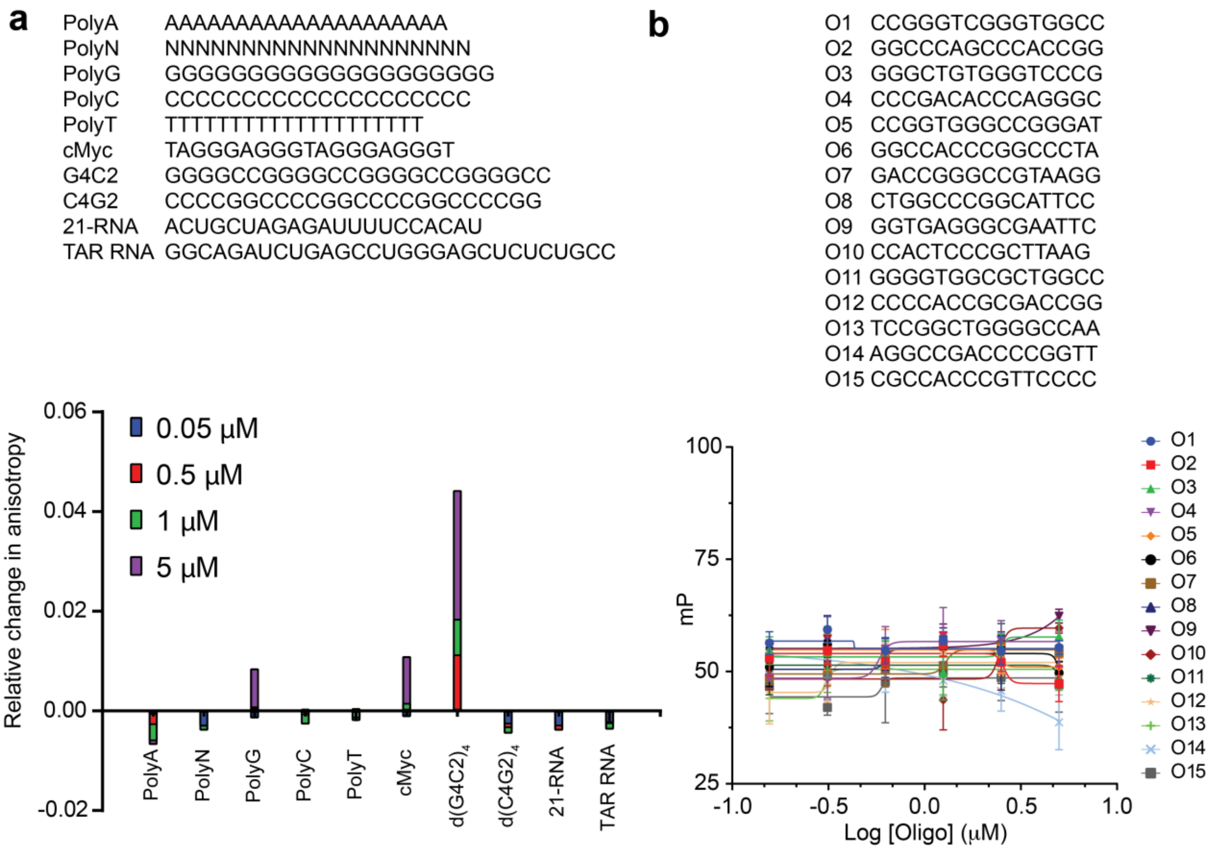


**Figure S3**. **(a)** Fluorescence anisotropy analysis, shown in lower panel was used to screen the binding specificity of AF488-Znf706 to a number of nucleotide sequences shown in the top panel that include DNA polynucleotides, DNA quadruplexes (cMyc and G4C2), a DNA C4G2 repeat, and single-stranded RNAs (21-mer and TAR) at the oligo concentration that is indicated in a color-coded fashion shown in the lower panel. The only sequences showing significant binding in this assay are PolyG, cMyc, and 4-repeat DNA G4C2, all of which are known to form G-quadruplexes. **(b)** Single-stranded DNA oligonucleotides named O1-O15 were designed with varying G-repeat sequences for a fluorescence polarization (FP) analysis that is shown in the lower panel. This assay shows changes in polarization value (mP) for 20nM AF-488 Znf706 titrated with a variable concentration of DNA oligonucleotides O1-O15 at the indicated colors. Though G-rich, none of the sequences are predicted to form G-quadruplexes and none appear to bind tightly to Znf706 using this assay.

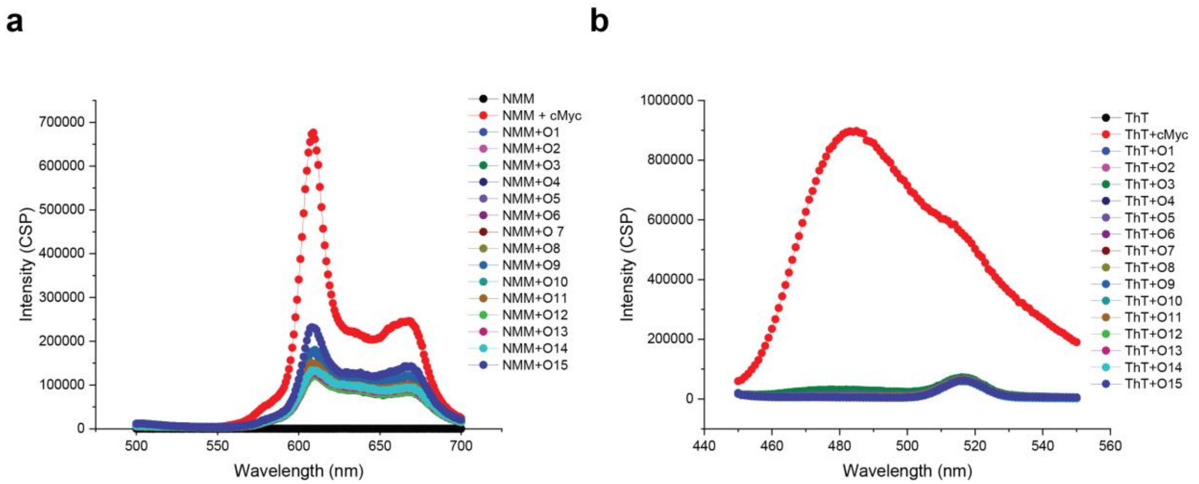


**Figure S4**. Monitoring the change in 10nM N-methylmesoporphyrin IX (NMM, **a**) and thioflavin-T (ThT, **b**) fluorescence that occurs upon mixing a 10x molar excess of G-rich oligonucleotides O1-O15 or the cMyc G-quadruplex shown as red filled circles (refer to sequence information in **Figs. S3a and b**).

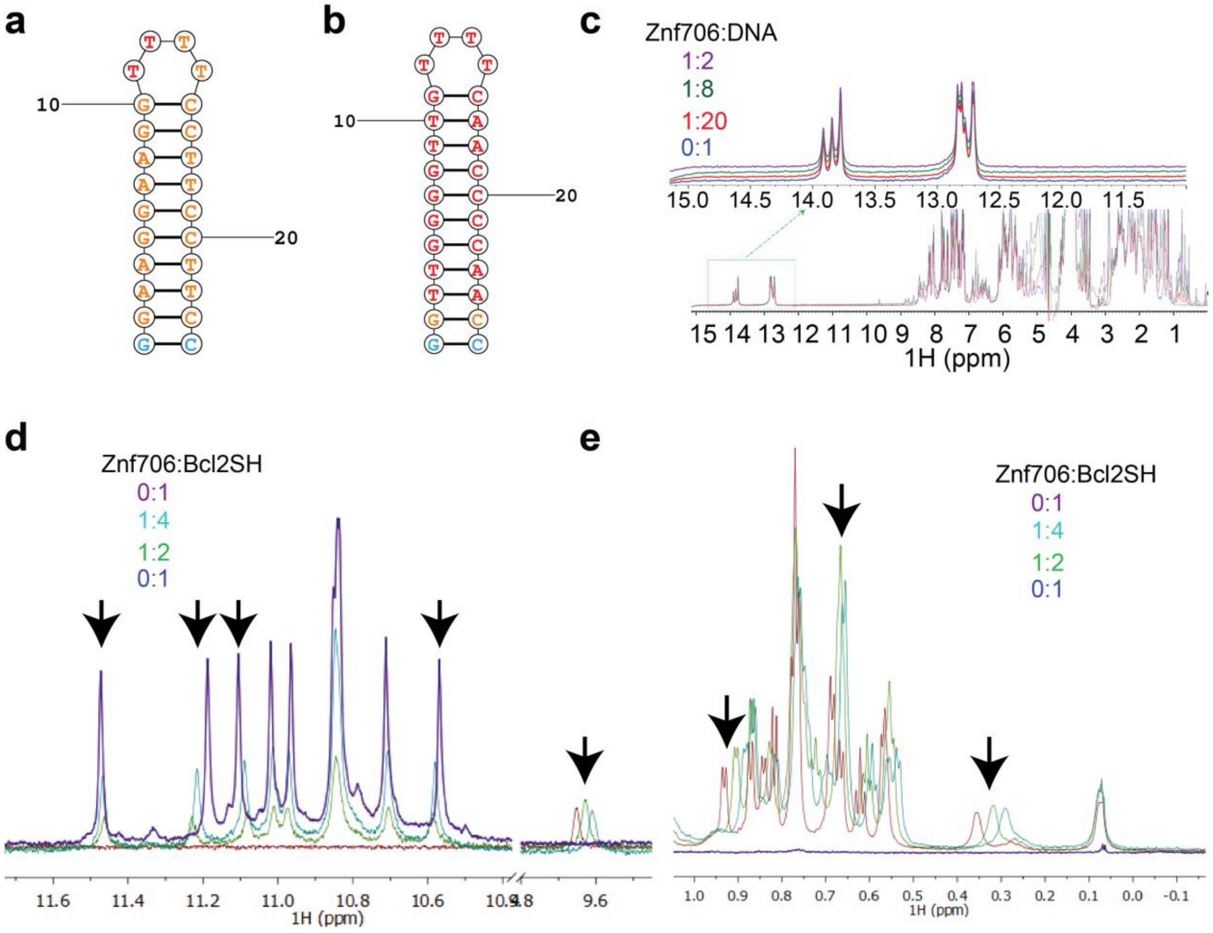


**Figure S5**. **(a-b)** Secondary structure prediction of GGA **(a)** and GGGG **(b)** hairpin DNA sequences. **(c)** 1D proton NMR spectra of 200 µM GGGG-hairpin oligonucleotide, recorded at 37 °C, titrated with variable concentrations of Znf706 as indicated in the designated colors. Zoomed spectra are shown to highlight that no chemical shift change occurs for the imino protons. A corresponding 1D spectrum of GGA showing no Znf706 induced chemical shifts in imino protons is shown in main text Fig. 1d. **(d-e)** 1D NMR spectrum of Bcl2SH G-quadruplex (200 µM) in the absence or presence of increasing concentrations of Znf706 as shown in the indicated colors. The imino **(d)** and methyl **(e)** protons showing chemical shift perturbations upon Znf706 addition are indicated with arrows. The 1D proton NMR spectrum of Znf706 in the absence of DNA is presented in maroon color.

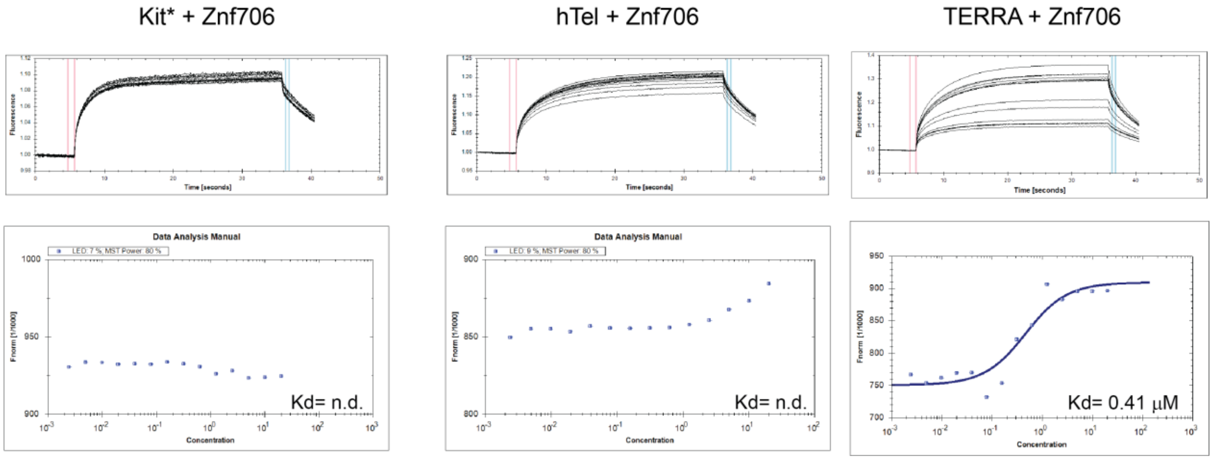


**Figure S6**. Microscale thermophoresis (MST) of the interaction between Znf706 and G-quadruplexes. MST traces of a titration of Znf706 (2.4 nM to 20 µM) against 100 nM DNA (hTel and Kit*) and RNA (TERRA: UA GGG UUA GGG UUA GGG UUA GGG) 5' 6-FAM G-quadruplexes. Traces on the top and bottom represent bound and unbound fractions, respectively. The MST curves are analyzed using the Nanotemper analysis program and the dissociation constant ($K_D$) as shown in the figure for each system was calculated using the equation

$$f(c) = unbound + \frac{(bound - unbound)}{2} \times (FluoConc + c + Kd - \sqrt{((FluoConc + c + Kd)^2 - 4 \times FluoConc \times c}).$$

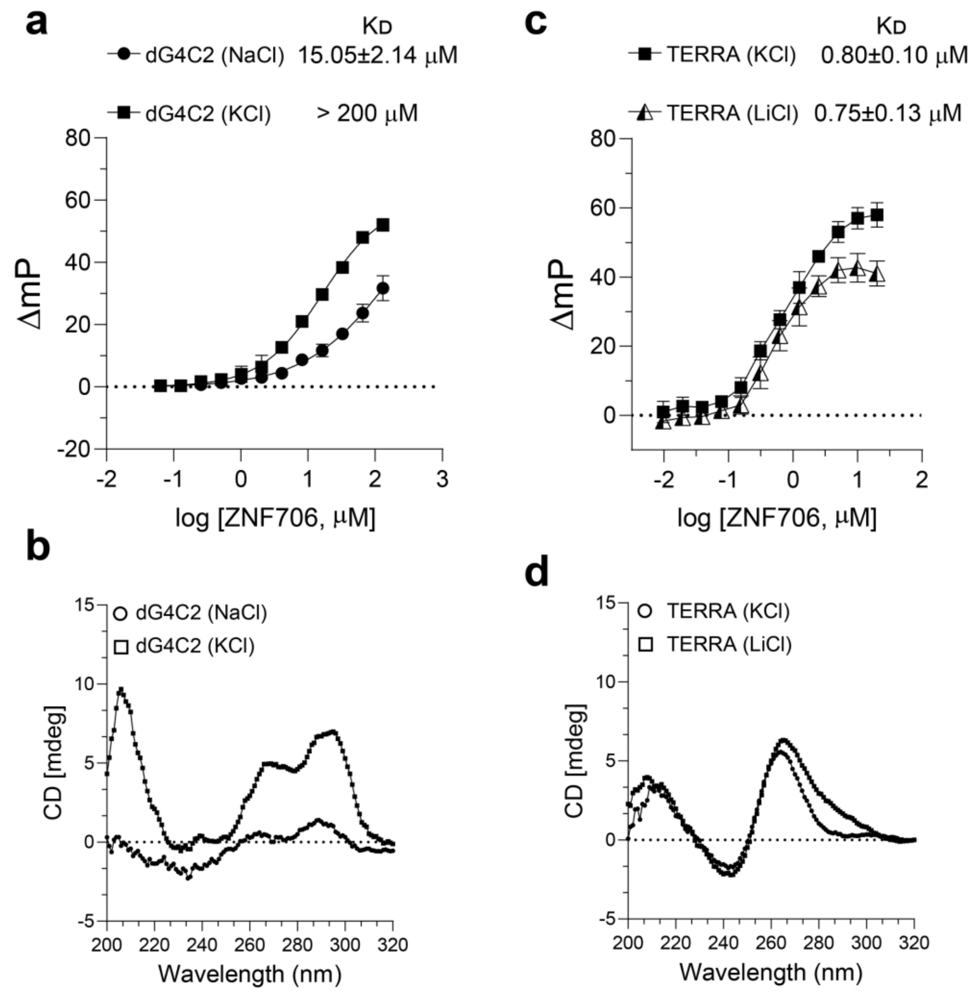


**Figure S7**. Fluorescence polarization (FP) assay measuring the binding affinities between Znf706 (variable concentration) and 100 nM 5’ 6-FAM labeled G4C2 4-repeat G-quadruplexes **(a)** or 100 nM 5’ 6-FAM labeled TERRA **(c)** dissolved in a buffer containing different salts as indicated. G4C2 oligos were prepared either in 20 mM NaPi (pH 7.4), 100 mM KCl or 20 mM NaPi (pH 7.4), 100 mM NaCl; and TERRA oligos were prepared in 20 mM NaPi (pH 7.4), 100 mM KCl or 20 mM Tris-HCl (pH 7.4), 100 mM KCl for FP assay. The CD spectra of G4C2 and TERRA oligos in respective buffers as mentioned above are shown in **b** and **d**, respectively. CD spectra were collected using 20 μM oligonucleotides at 25 °C using a JASCO J-1500 spectropolarimeter.

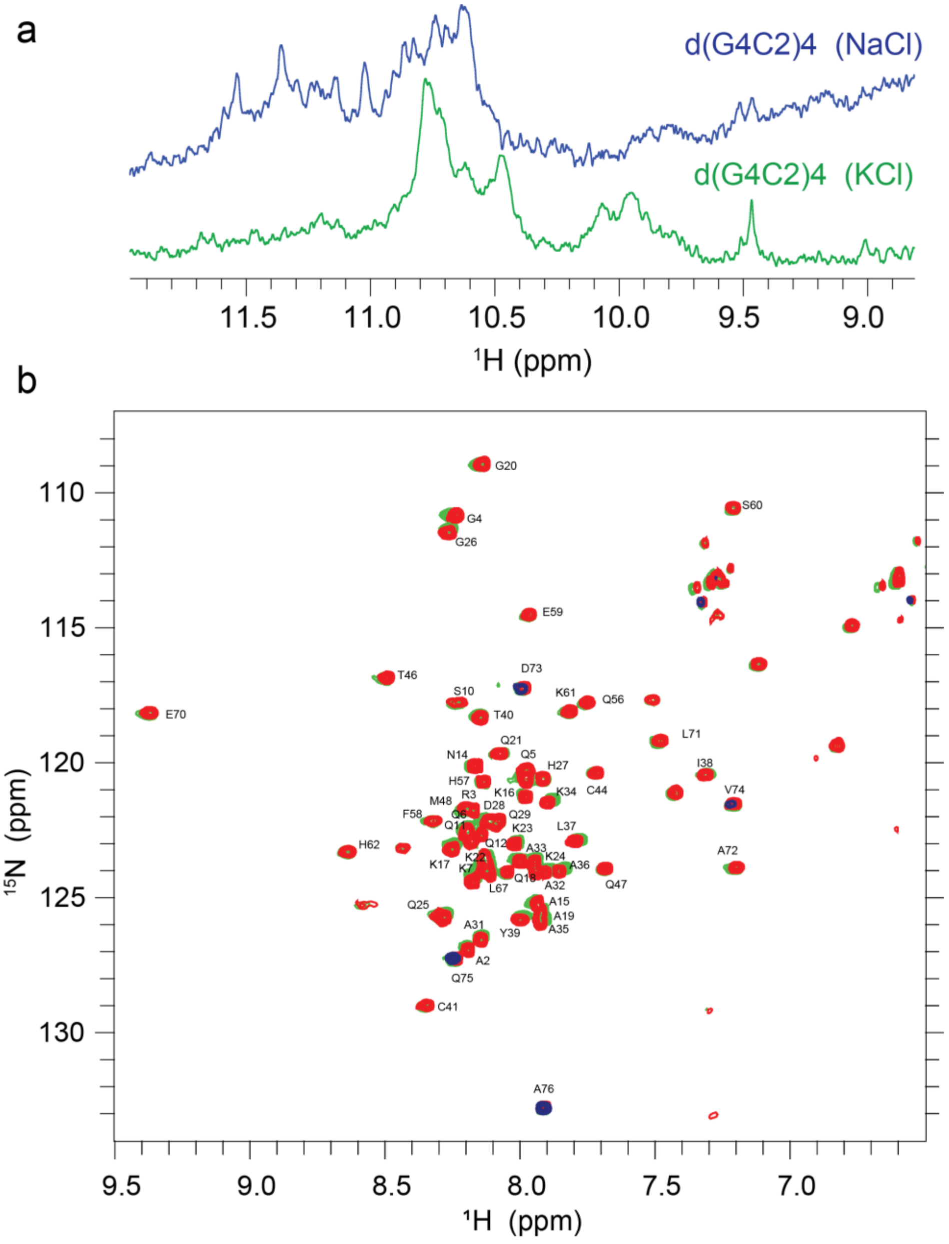


**Figure S8**. **(a)** 1D NMR spectra of 100 µM G4C2 4-repeat DNA sequence dissolved in 20 mM NaPi, 100 mM KCl, pH 7.4 (blue) and 20 mM NaPi, 100 mM NaCl, pH 7.4 (green). **(b)** TROSY spectrum of 50 µM $^{15}N$ Znf706 (red spectrum) mixed with 100 µM $(G4C2)_4$ prepared in 20 mM NaPi, 0.05% $NaN_3$, 7.5% $D_2O$, pH 7.4, 100 mM KCl (blue spectrum) or in 20 mM NaPi, 7.5% $D_2O$, 0.05% $NaN_3$, pH 7.4, 100 mM NaCl (green spectrum). The TROSY spectra were acquired at 4 °C on a Bruker 600 MHz spectrometer.

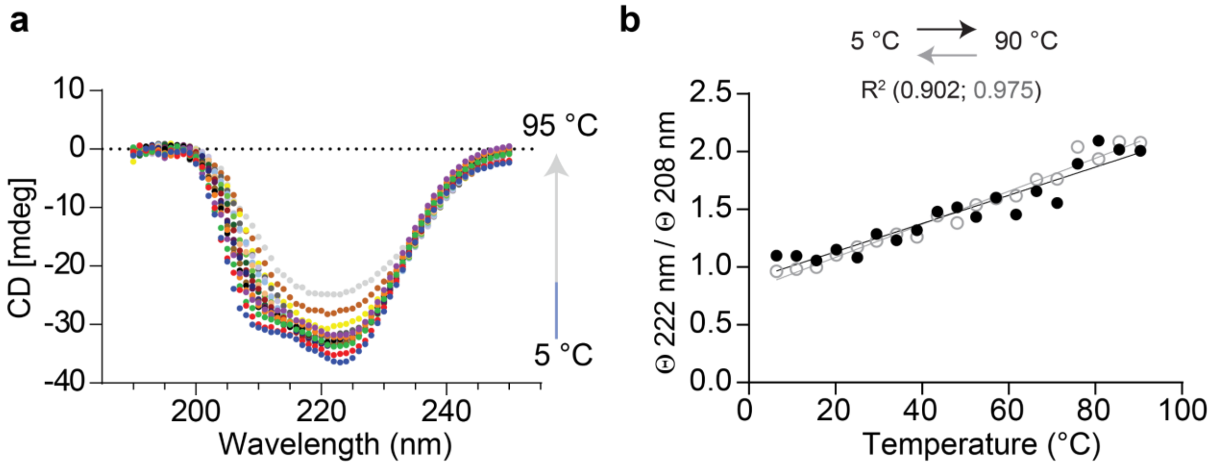


**Figure S9**. **(a)** Monitoring Znf706 (50 µM) conformational changes as a function of temperature by CD spectroscopy. CD spectra were recorded in every 5 °C interval in temperatures ranging from 5 to 95 °C. **(b)** Monitoring the conformational reversibility in 50 µM Znf706 by CD spectroscopy as a function of the change in CD signal intensity ratio at 222 and 208 nm. The folding and unfolding are reversible. However, the direct correlation between temperature and Znf706 secondary structure suggests a highly noncooperative folding transition. Solid and open circles represent ellipticity recorded during the melting (5 to 90 °C) and cooling cycle (90 to 5 °C), respectively.

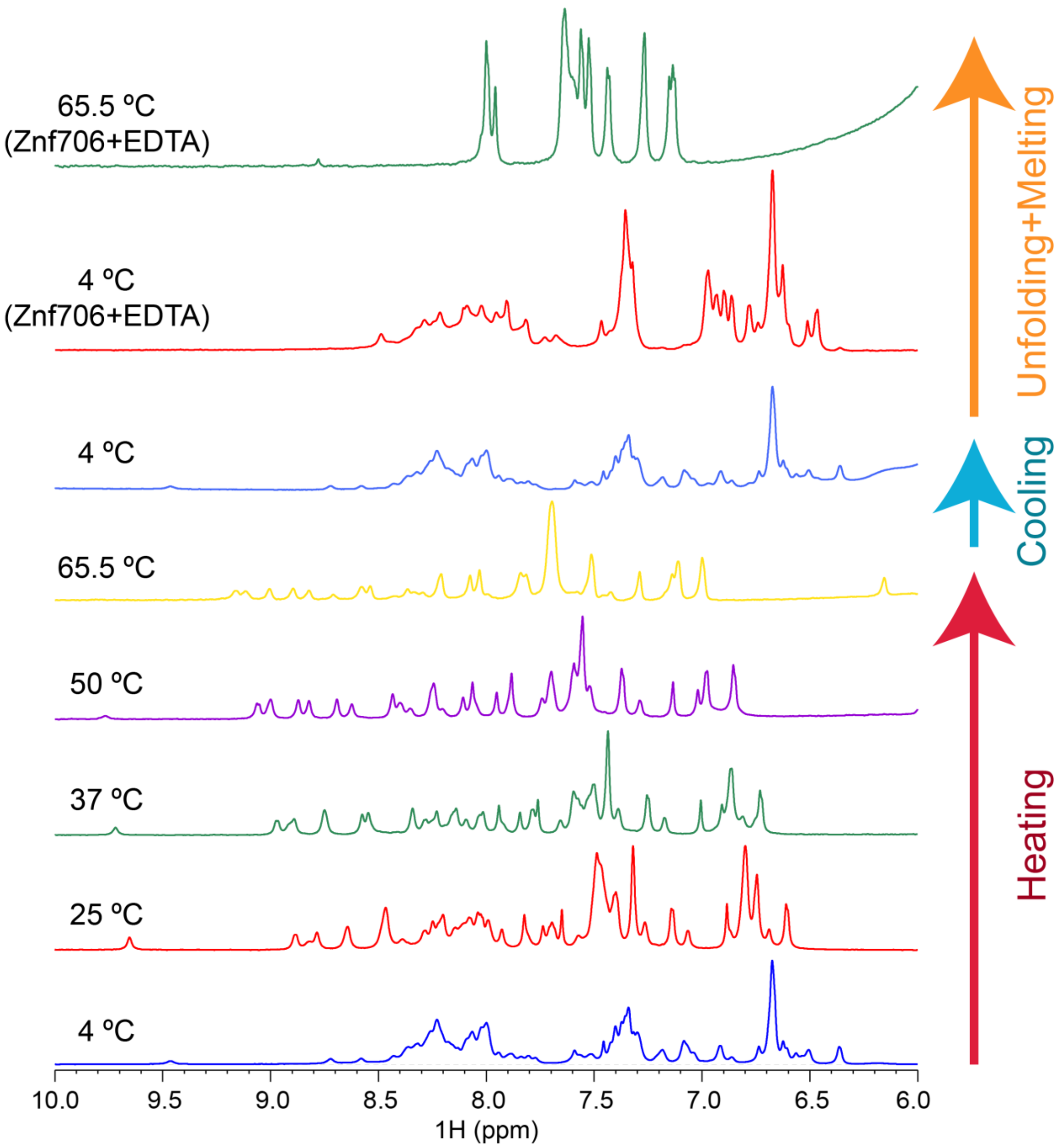


**Figure S10**. Monitoring thermostability of Znf706 using 1D proton NMR. Proton NMR spectra of 100 µM Znf706 dissolved in 20 mM NaPi, 100 mM KCl, 0.05% $NaN_3$, pH 7.4 were recorded at the indicated temperatures. Znf706 was slowly heated from 4 °C to 65.5 °C (red arrow) and then cooled back to 4 °C (blue arrow) followed by 10-molar excess EDTA treatment to test its effect on Znf706 structure (orange arrow). 1D NMR spectra were recorded on a Bruker 800 MHz spectrometer.

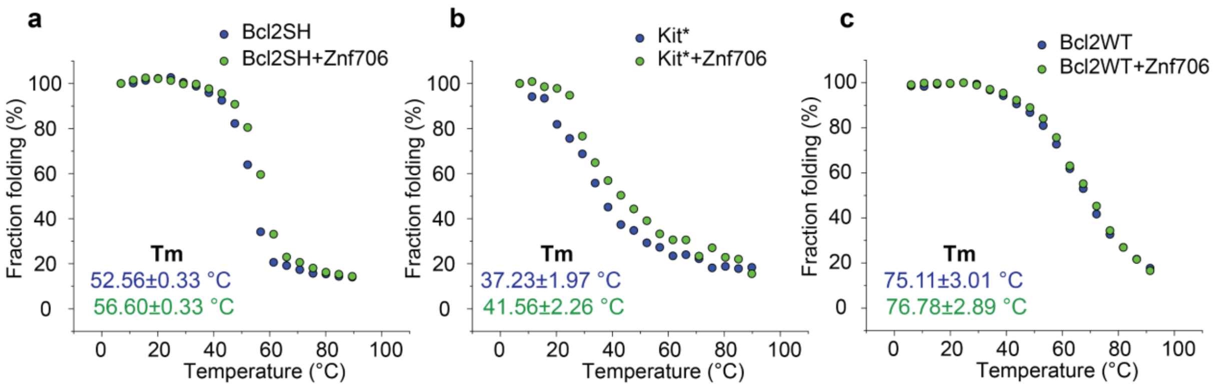


**Figure S11**. CD melting temperature (Tm) analysis of 20 µM Bcl2SH **(a),** Kit* **(b),** and Bcl2WT **(c)** in the absence (blue) or presence of equimolar Znf706 (green).

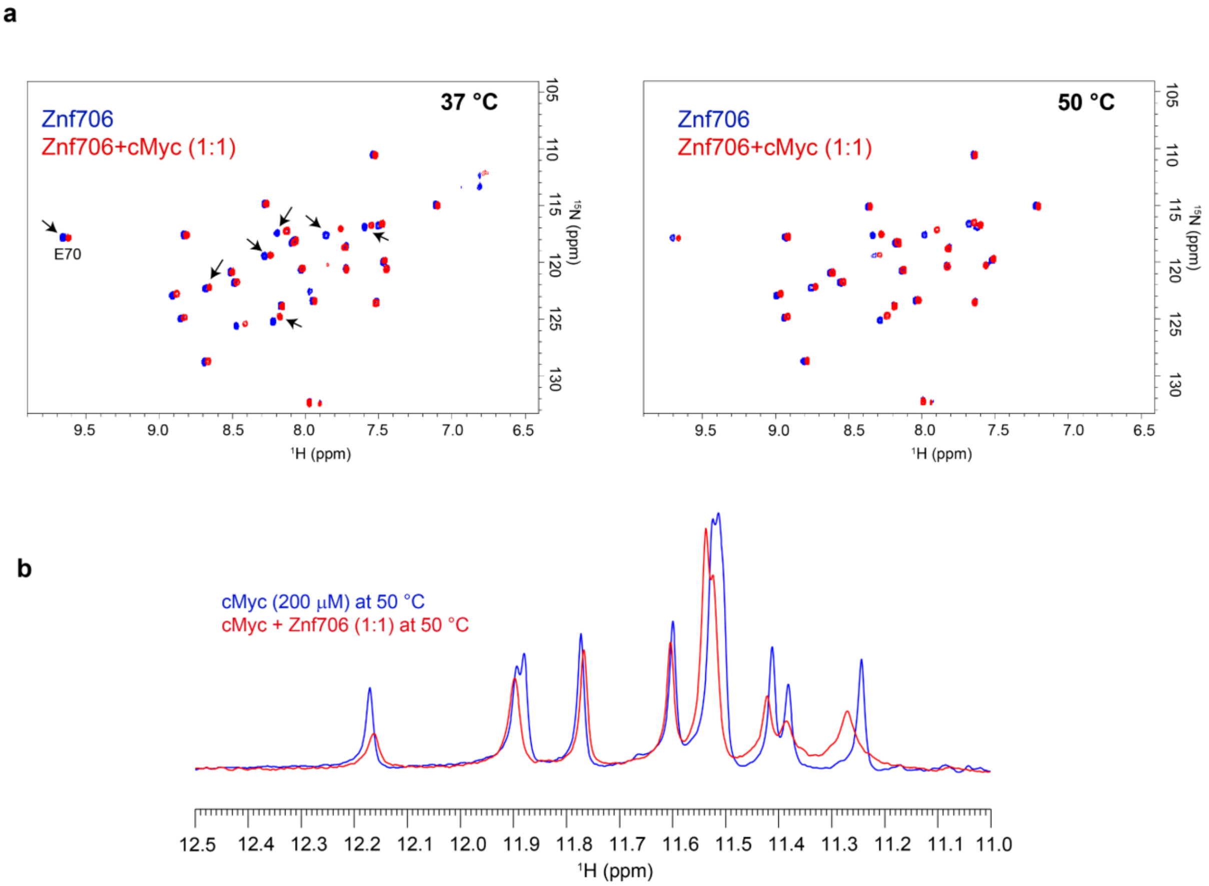


**Figure S12**. **(a)** Overlaid 2D TROSY $^{15}N/^{1}H$ NMR spectrum of 200 µM Znf706 (blue) mixed with equimolar cMyc G-quadruplexes (red) recorded on an 800 MHz Bruker spectrometer at 37 or 50 °C. Arrows indicate Znf706 residues showing chemical shift perturbation upon cMyc binding that are unassigned in this study at these indicated temperatures. **(b)** Overlaid 1D proton NMR spectrum of 200 µM cMyc imino protons (blue) mixed with equimolar Znf706 (red). The corresponding chemical shift changes in cMyc imino protons recorded at 37 °C are shown in the main text Figure 3a.

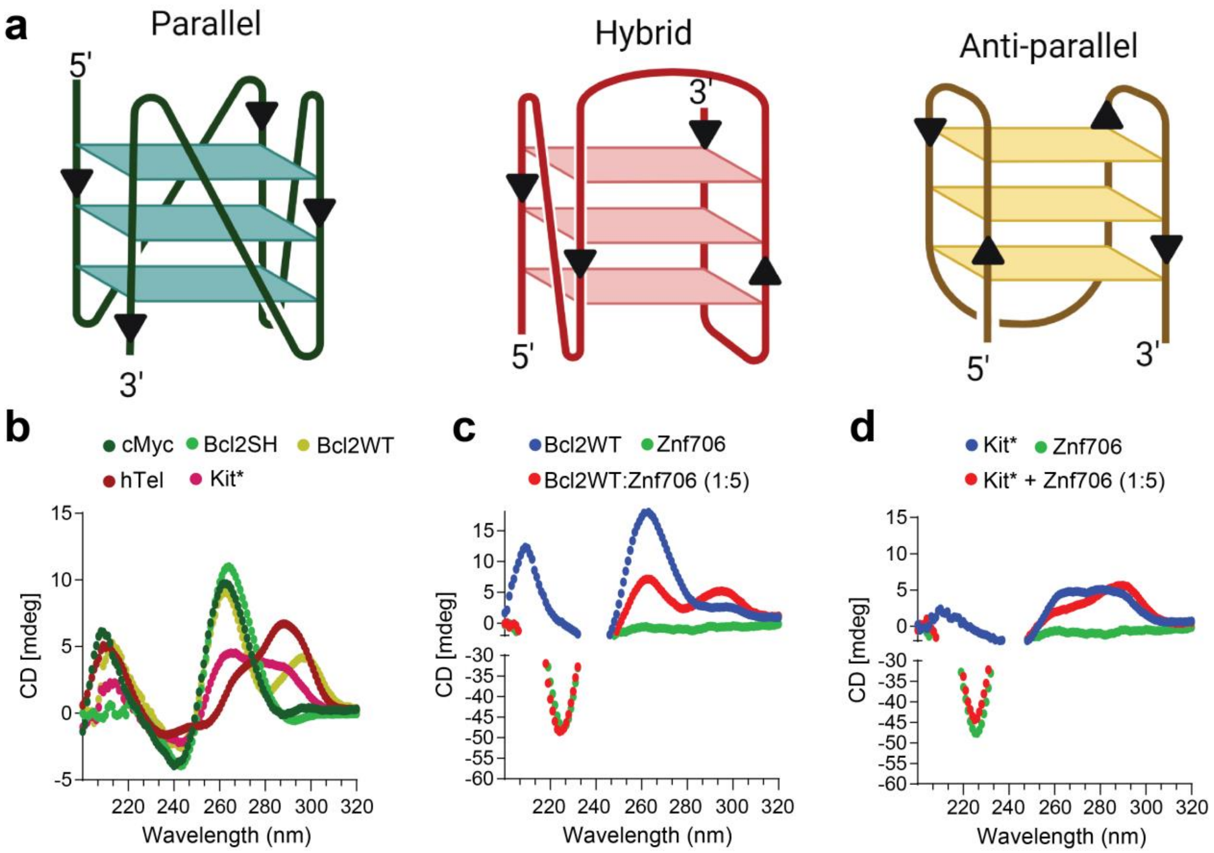


**Figure S13**. **(a)** Schematic representation of three different G-quadruplex topologies. **(b)** CD spectrum of 20 µM G-quadruplexes is characterized by three different topologies. CD spectrum for cMyc and Bcl2SH is typical for quadruplexes with parallel topology; hybrid topology is indicated for hTel, Kit*, and BCl2WT. **(c-d)** Effect of Znf706 on the refolding of Bcl2WT **(c)** and Kit* (**d**) as studied using CD spectroscopy. Addition of a 5x molar excess Znf706 induces antiparallel conformation of 20 µM Bcl2WT and Kit* prepared in 20 mM NaPi, 4 mM KCl, pH 7.4.

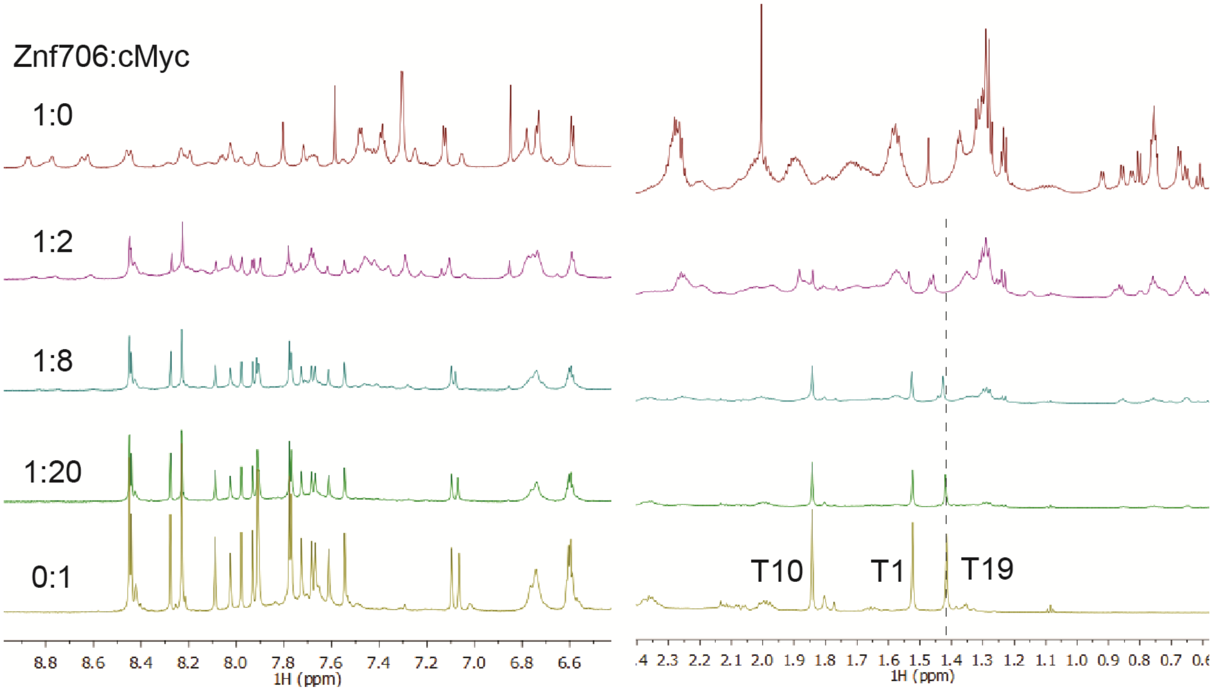


**Figure S14**. 1D NMR spectra (aromatic regions: left, methyl regions: right) of 200 µM cMyc mixed with increasing concentrations of Znf706. The methyl peaks are labeled, and the dashed line indicates a chemical shift change in T19 that occurs upon Znf706 binding.

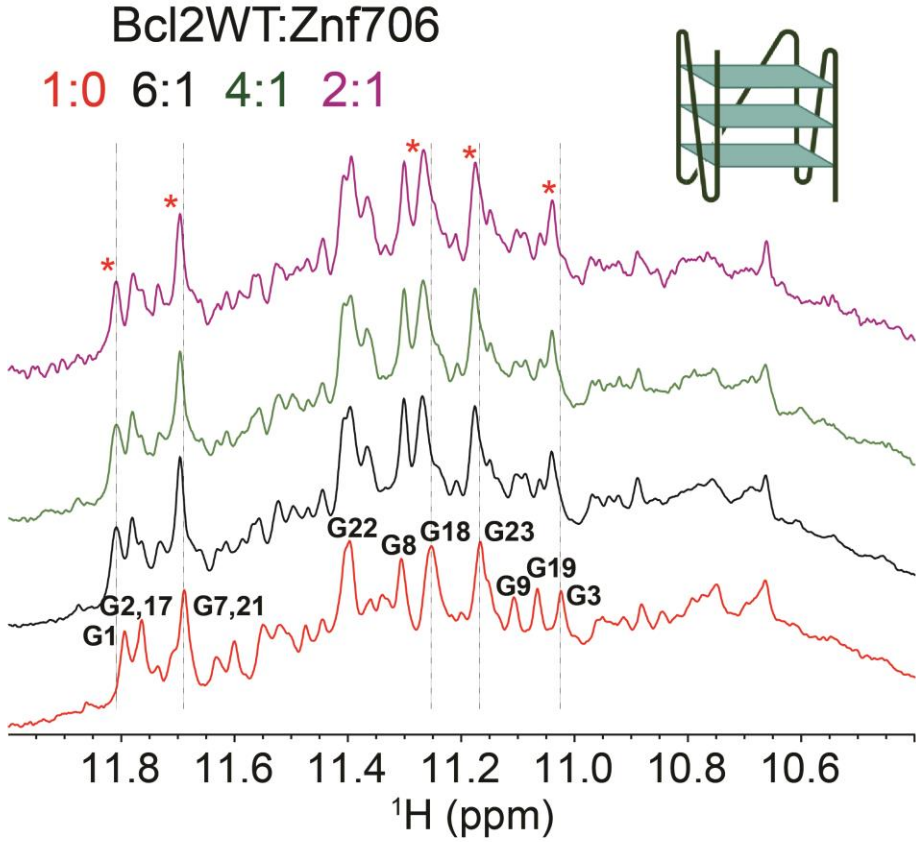


**Figure S15**. Mapping of the specific interaction sites within the Bcl2WT structure with which Znf706 binds using 1D proton NMR. The guanines in Bcl2WT that have assigned chemical shifts[14] are labeled on the red spectrum. When increasing concentrations of Znf706 are added, several Bcl2WT G-quadruplex imino protons show chemical shift changes as highlighted by the dashed lines. These represent guanines directly involved in Znf706 interaction and are denoted with a red asterisk "*".

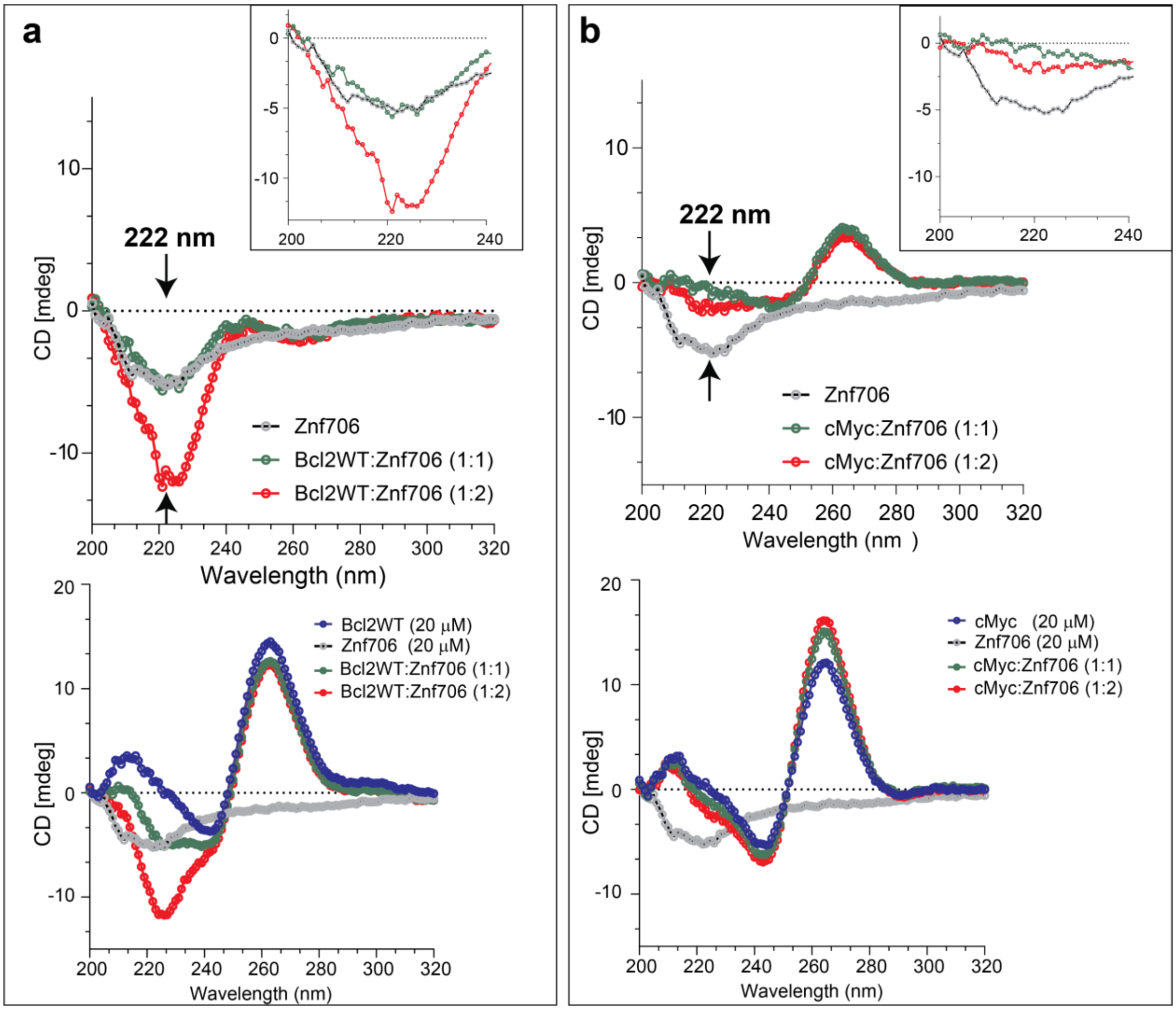


**Figure S16**. CD spectroscopy analysis to probe the binding effect of Znf706 binding on cMyc **(a)** and Blc2WT **(b)** G-quadruplex secondary structures. 20 µM G-quadruplexes were folded in 20 mM NaPi, 100 mM KCl, pH 7.4 and the effect of Znf706 addition dissolved in an identical buffer was monitored at 25 °C at the indicated molar ratios by subtracting the CD signals of the buffer (left panel, **a** and **b**), and CD signals of G-quadruplexes for the Znf706 alone spectrum (right panel, **a** and **b**). The Znf706 signal shown in the inset shows the CD region that corresponds to the secondary structure change in Znf706 in the G-quadruplex mixture sample. The arrows indicate regions ~218-222 nm where G-quadruplexes have a minimum (~ 0) and Znf706 has a maximum CD absorbance.

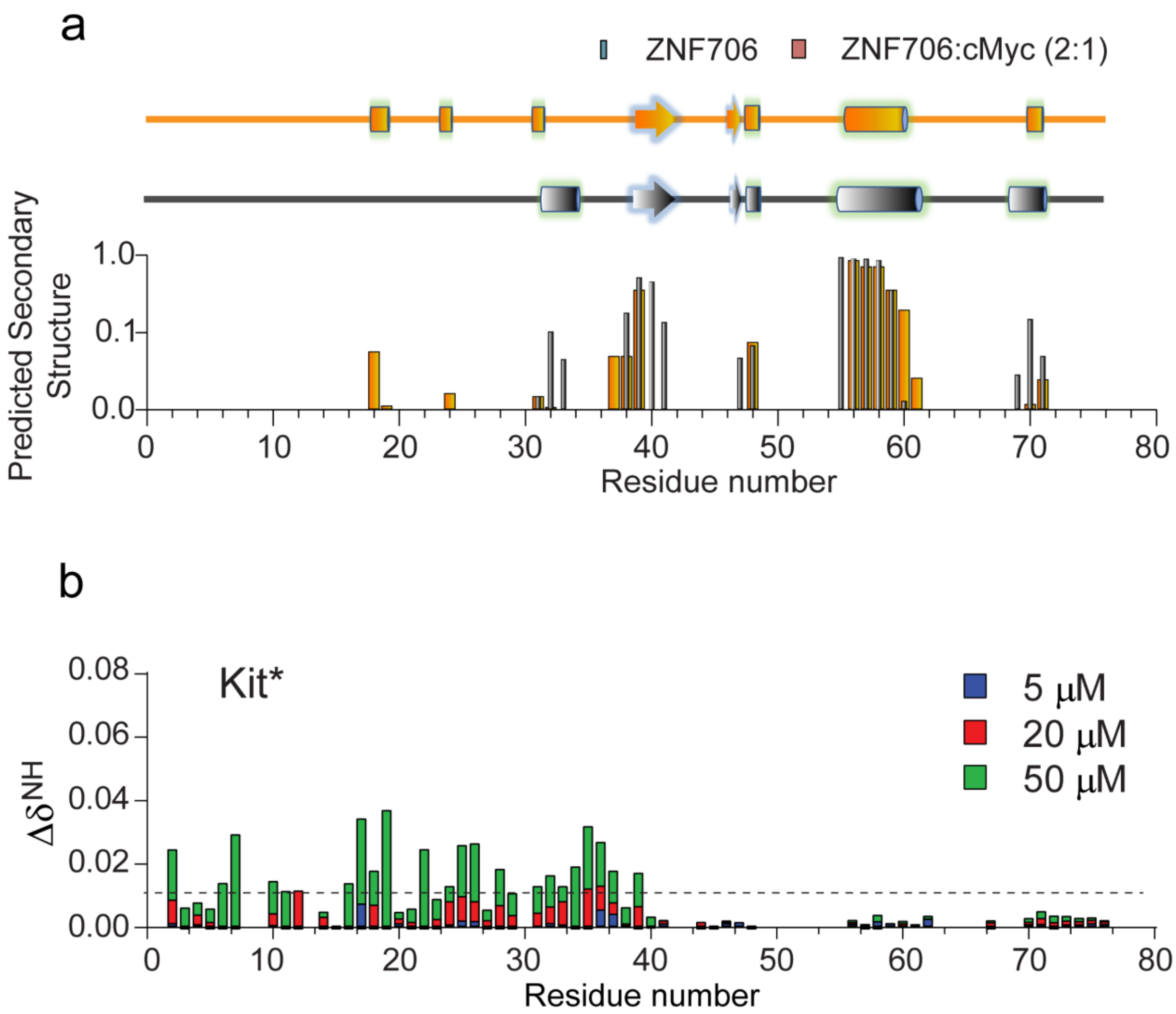


**Figure S17**. **(a)** Predicted secondary structure of Znf706 (black) or Znf706-cMyc (orange) obtained from backbone NMR chemical shifts (Cα, Cβ and CO) using the TALOS-N program. **(b)** The chemical shift perturbations (CSPs) plots are derived from the $^{15}N/^{1}H$ 2D NMR spectrum of 100 µM Znf706 titrated with variable concentrations of Kit* as indicated in colors. The CSPs are calculated using equation $\Delta\delta_{NH} = \sqrt{(\delta^{1}H)^{2} + 0.154 \times (\delta^{15}N)^{2}}$ and the dashed lines indicate the average CSPs**.**

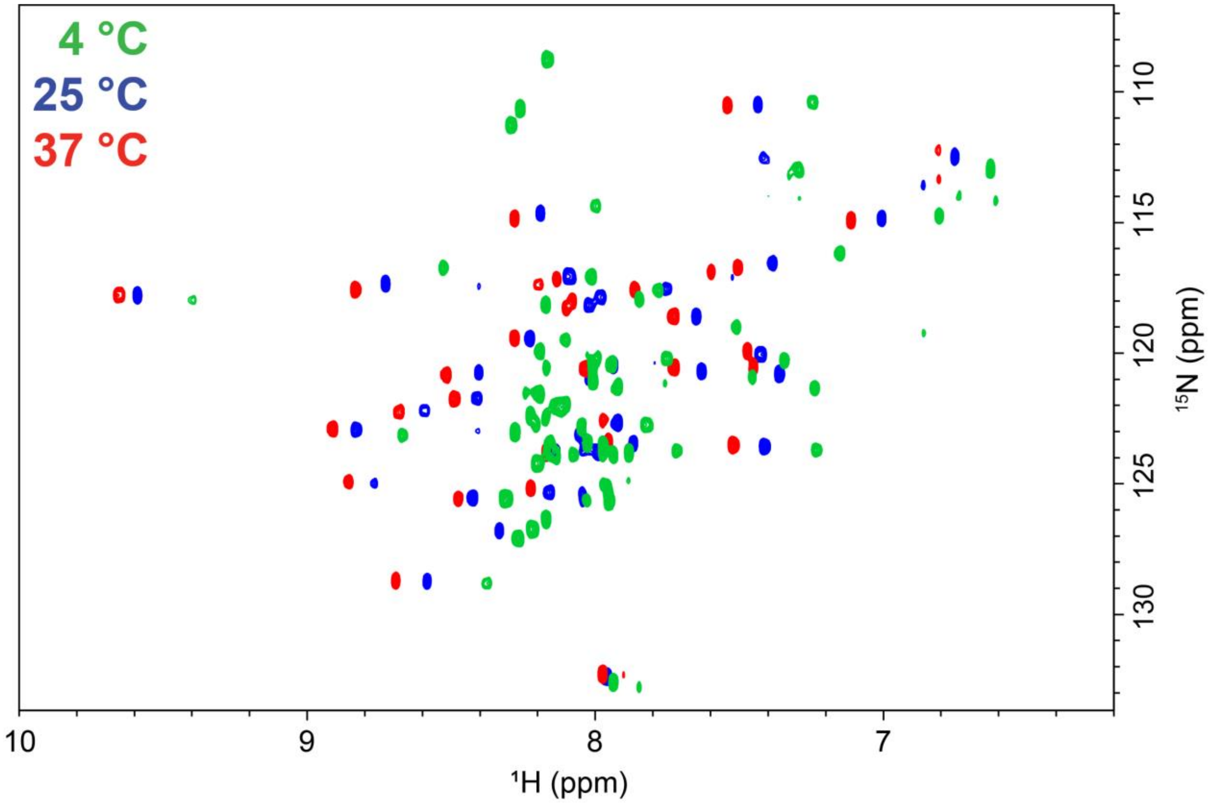


**Figure S18**. Overlaid $^{15}N/^{1}H$ TROSY spectrum of 100 μM Znf706 dissolved in 20 mM NaPi, 100 mM KCl, pH 7.4, and 8% $D_2O$ recorded on an 800 MHz spectrometer at the indicated temperatures.

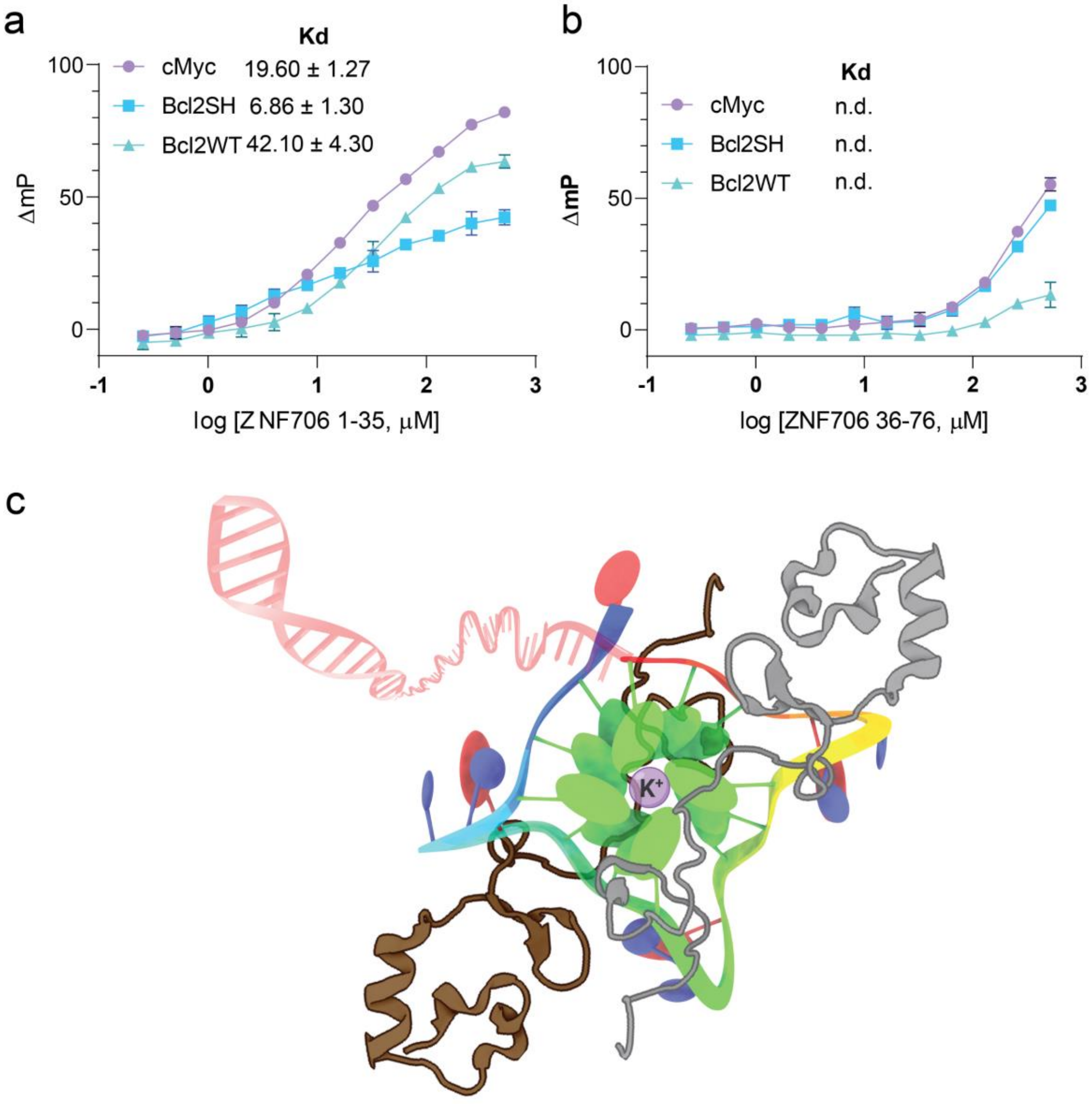


**Figure S19**. Fluorescence polarization plots measuring the binding affinities between the N- and C-terminus fragments of Znf706 and G4 quadruplexes. Binding titrations of Znf706 N-terminal residues 1-35 **(a)** and C-terminal zinc-finger residues 36-76 **(b)** and 5’ 6-FAM-labeled G-quadruplexes prepared in 20 mM NaPi, 100 mM KCl, pH 7.4. The indicated $K_d$ values were calculated by non-linear regression analysis using a one-site binding saturation model in GraphPad Prism at increasing concentrations of protein (0.25 µM to 520 µM). Error bars represent s.d derived from three replicates. n.d.: not determined. **(c)** A proposed binding model for Znf706 and G-quadruplex complex where two Znf706 molecules are shown in light and dark grey, and the G-quadruplex in green.

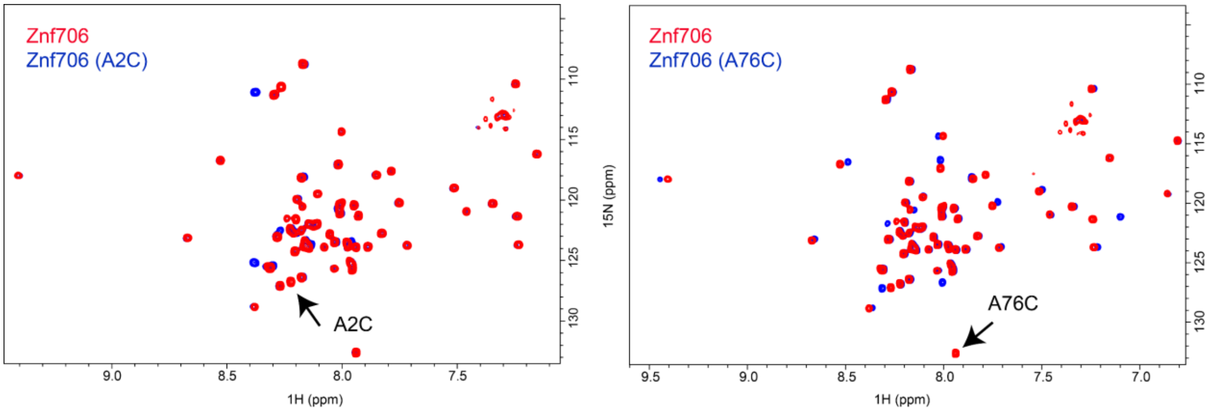


**Figure S20**. Overlay TROSY spectrum of 100 µM $^{15}N$ labeled Znf706 in red and Znf706 A2C (left) or A76C (right) in blue prepared in 20 mM NaPi, 100 mM KCl, 0.05% $NaN_3$, pH 7.4 containing 7.5% $D_2O$ recorded on a Bruker 800 MHz at 4°C.

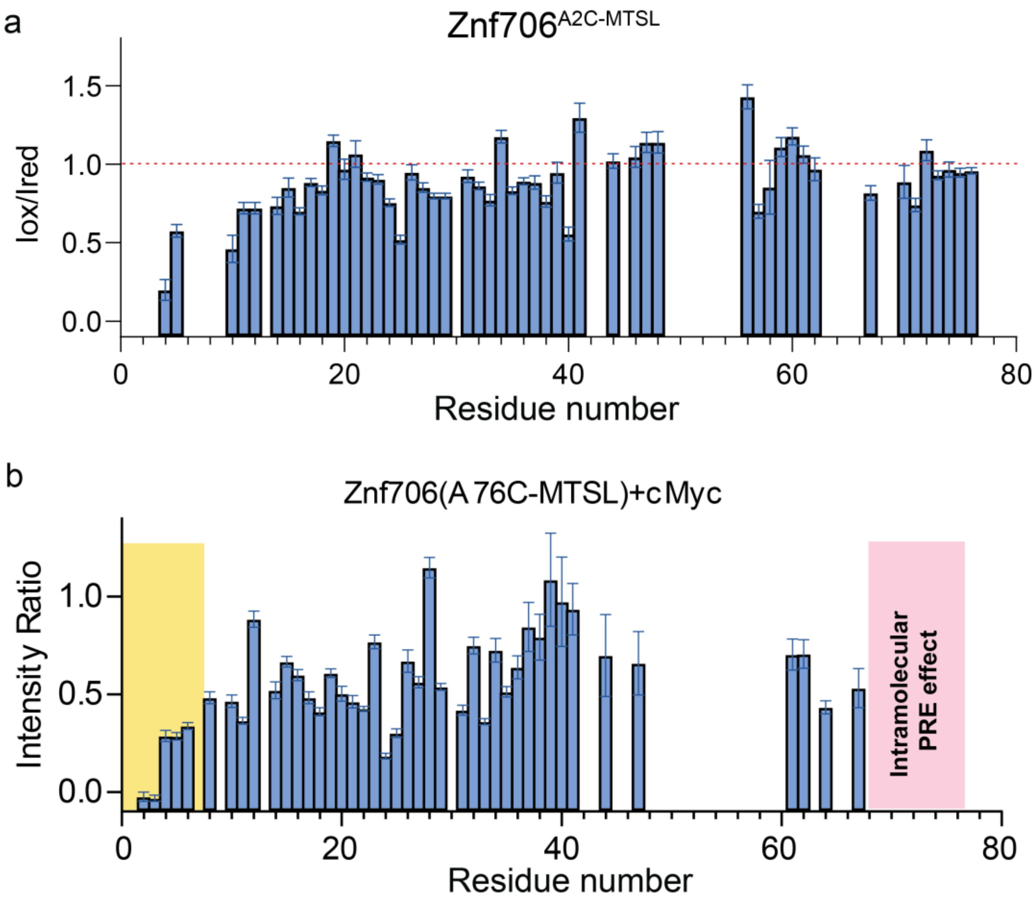


**Figure S21**. **(a)** Intensity ratio of amide protons observed for 100 µM $^{15}$N Znf706 A2C-MTSL mixed with or without 50 µM cMyc G-quadruplex as indicated in oxidized (paramagnetic state) and reduced (diamagnetic state) conditions. The Znf706 A2C-MTSL NMR sample was reduced with 5-molar excess sodium-ascorbate for ~3 hour prior to recording the diamagnetic spectrum. The NMR data were recorded on a Bruker 800 MHz spectrometer at 4 °C for samples dissolved in 20 mM NaPi, 0.05% $NaN_3$, 100 mM KCl, pH 7.4 buffer containing 7.5% $D_2O$. **(b)** Signal intensity ratio of amide protons observed for 100 µM $^{15}$N Znf706 A76C-MTSL mixed with 50 µM cMyc G-quadruplex. Standard errors are estimated from the signal-to-noise ratios. The intramolecular PRE effects at C-terminus are shaded in pink, whereas cMyc binding induced PRE effects are highlighted in yellow. NMR spectra were collected on a Bruker 800 MHz spectrometer at 4 °C for samples dissolved in 20 mM NaPi, 100 mM KCl, pH 7.4 buffer containing 7.5% $D_2O$.

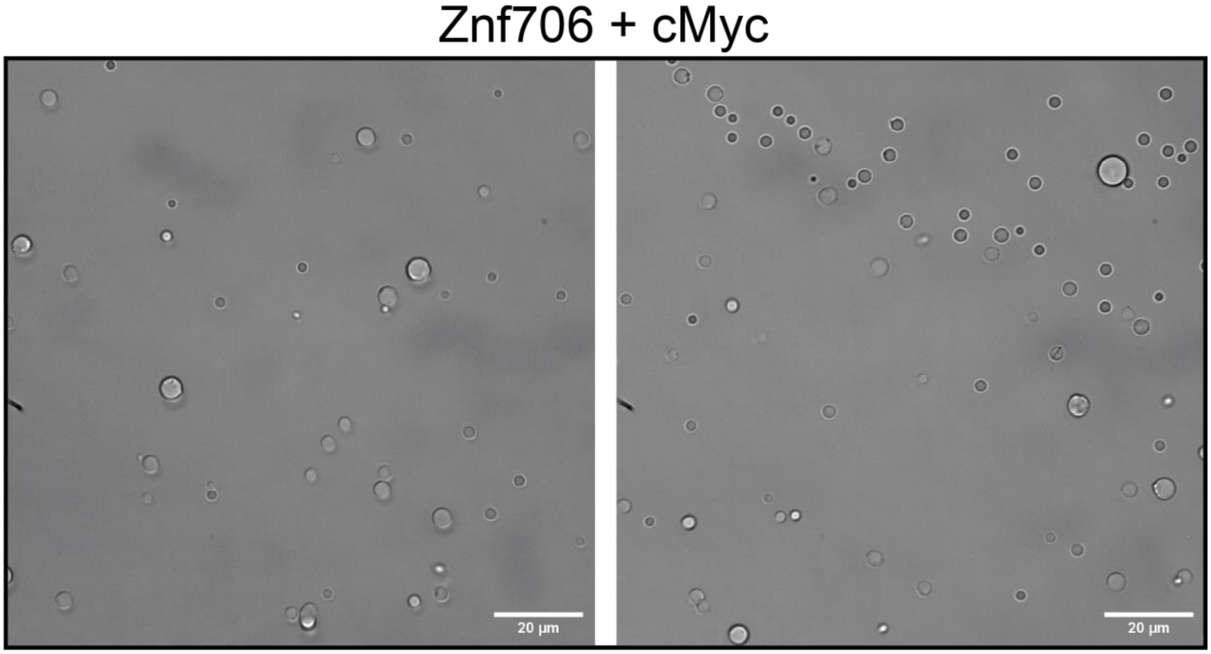


**Figure S22**. Differential interference contrast (DIC) images showing the formation of droplets in a sample mixture containing 100 µM Znf706 and 100 µM cMyc incubated overnight at room temperature. The sample mixture was dissolved in 20 mM NaPi, 100 mM KCl, pH 7.4, and 7.5% $D_2O$ inside a 0.5 mL NMR tube and transferred to a 16-well CultureWell™ before imaging.

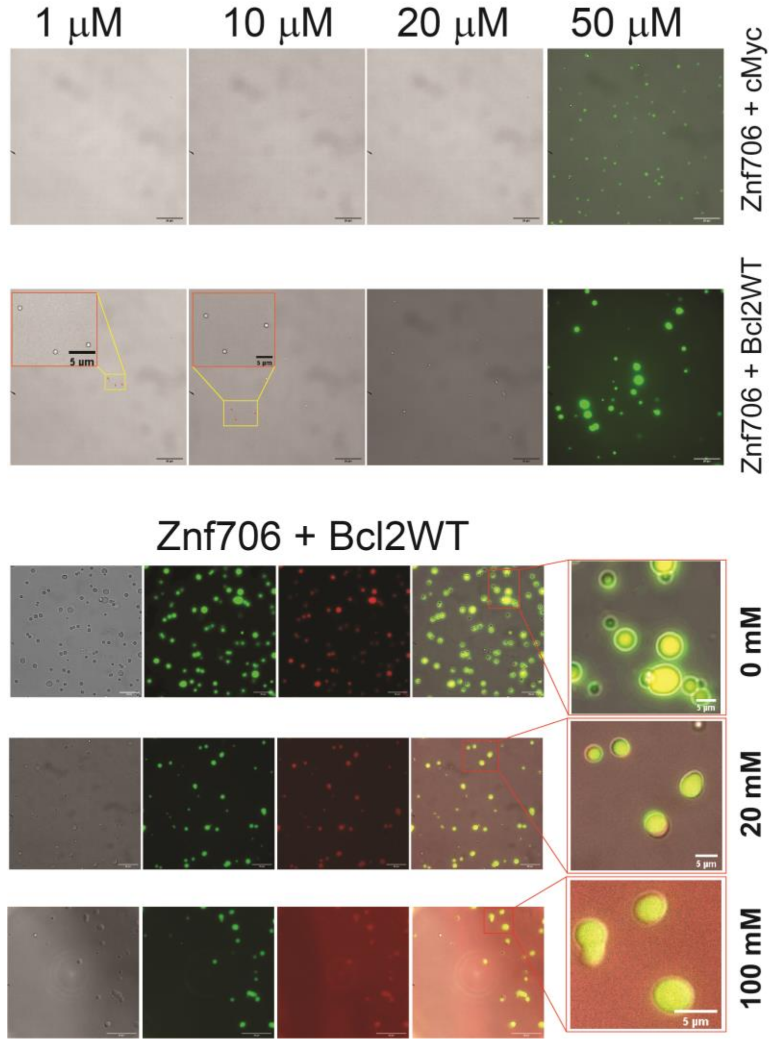


**Figure S23**. Fluorescence imaging to test the formation of LLPT in Znf706 at different concentrations (1 μM to 50 μM) mixed with an equimolar amount of cMyc or Bcl2WT (top). Monitoring the formation of LLPT in a sample mixture containing 100 μM Znf706 and an equimolar amount of Bcl2WT dissolved in 20 mM NaPi, pH 7.4 at the indicated KCl concentrations (bottom). For fluorescence imaging, a sample mixture ratio of 1:100 AF488-Znf706:unlabeled-Znf706 was used. The samples were incubated overnight at room temperature before imaging. LLPT formation was confirmed by DIC following the addition of 5 μM NMM to the sample mixture. The green and red signals correspond to AF488-Znf706 and NMM, respectively.

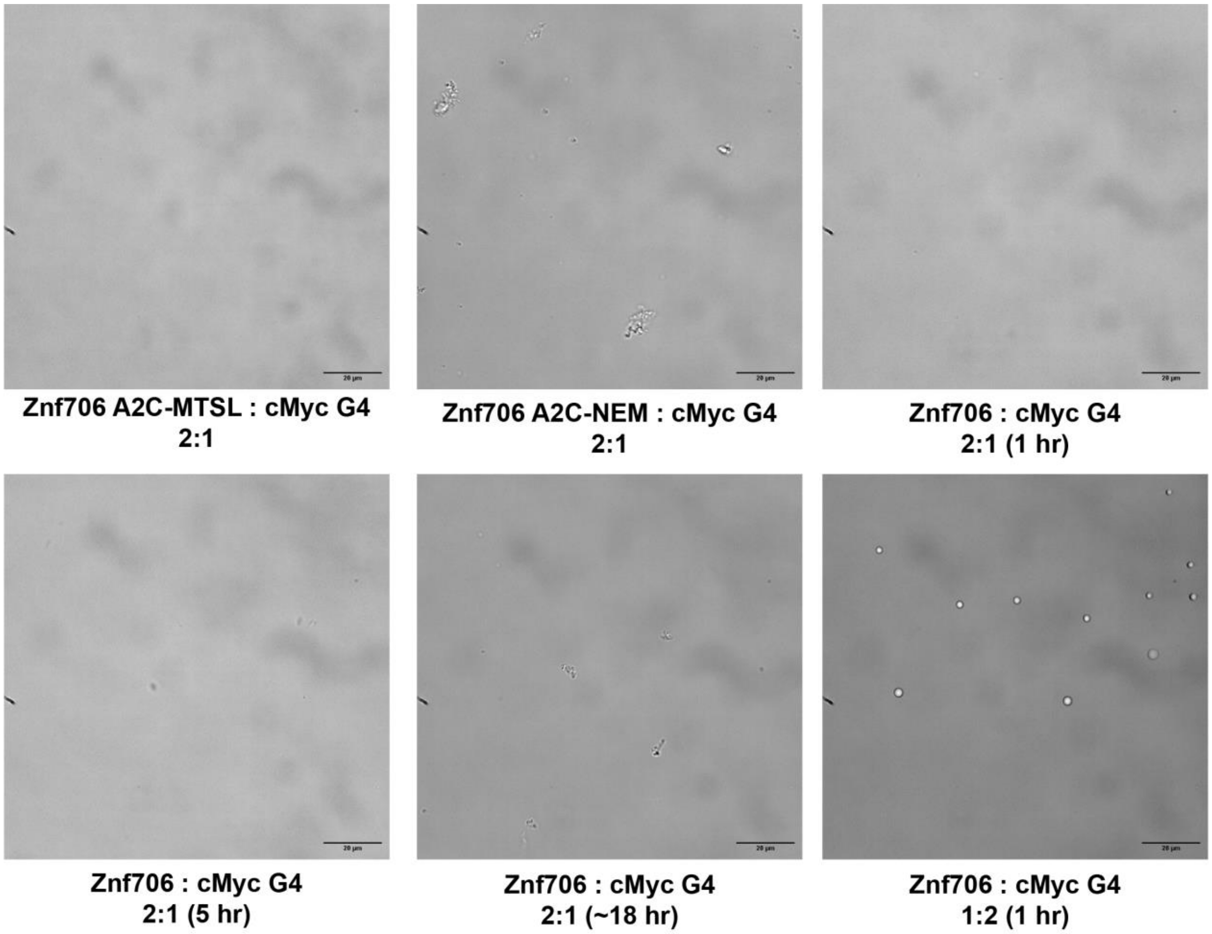


**Figure S24**. DIC images monitoring liquid-liquid phase transition in Znf706 (100 µM) or Znf706-A2C (100 µM) labelled with MTSL, or NEM incubated with 50 µM of cMyc G4 at room temperature at the indicated time points (top panel). The samples were dissolved in 20 mM NaPi, 100 mM KCl, pH 7.4, and 7.5% $D_2O$ corresponding to the NMR samples (protein:G4=2:1) used for 2D and PRE NMR measurements. The scale bar is 20 µm. Liquid droplets were observed in sample mixtures containing 50 µM Znf706 and 100 µM cMyc as indicated within ~1 hour of incubation at room temperature.

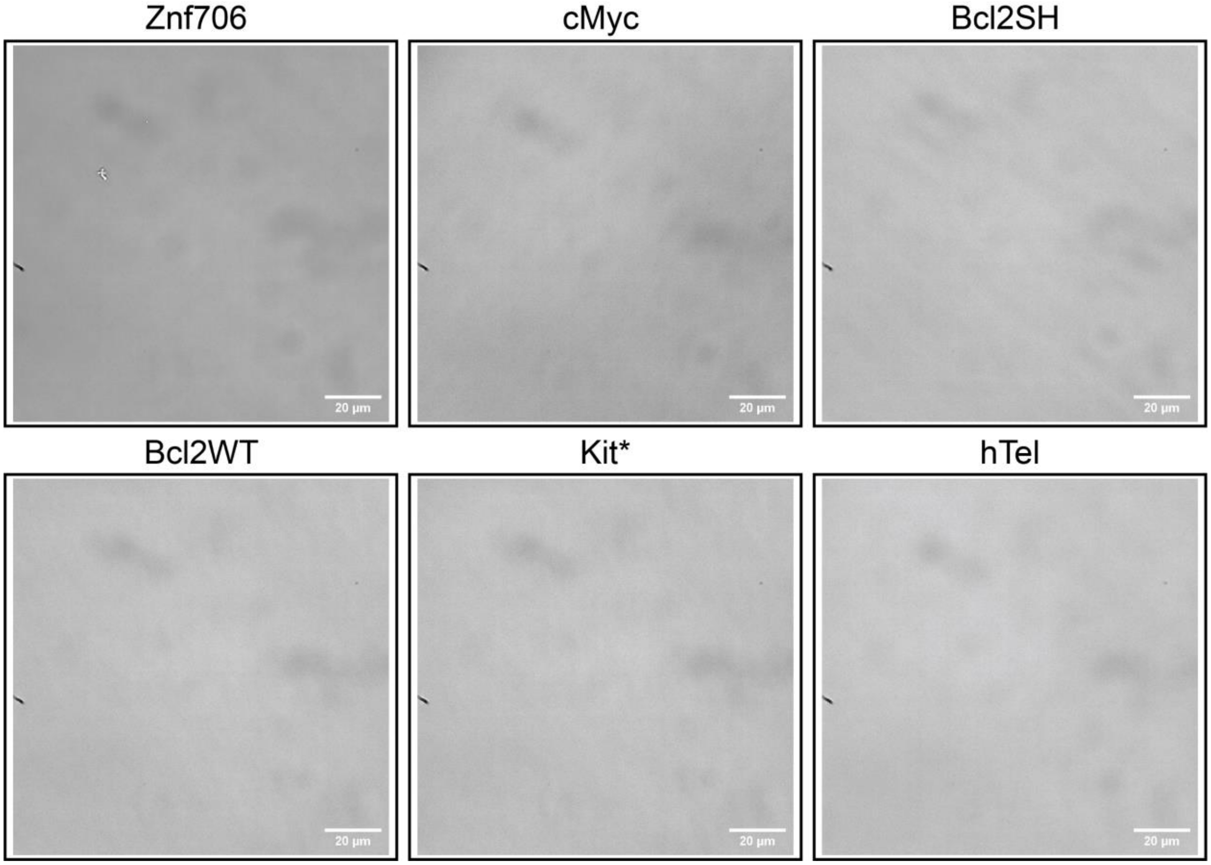


**Figure S25**. DIC images showed no LLPT formed when 100 µM Znf706 or G-quadruplexes alone were dissolved in 20 mM NaPi, 100 mM KCl, 7.5% $D_2O$, pH 7.4 and incubated overnight at room temperature before imaging.

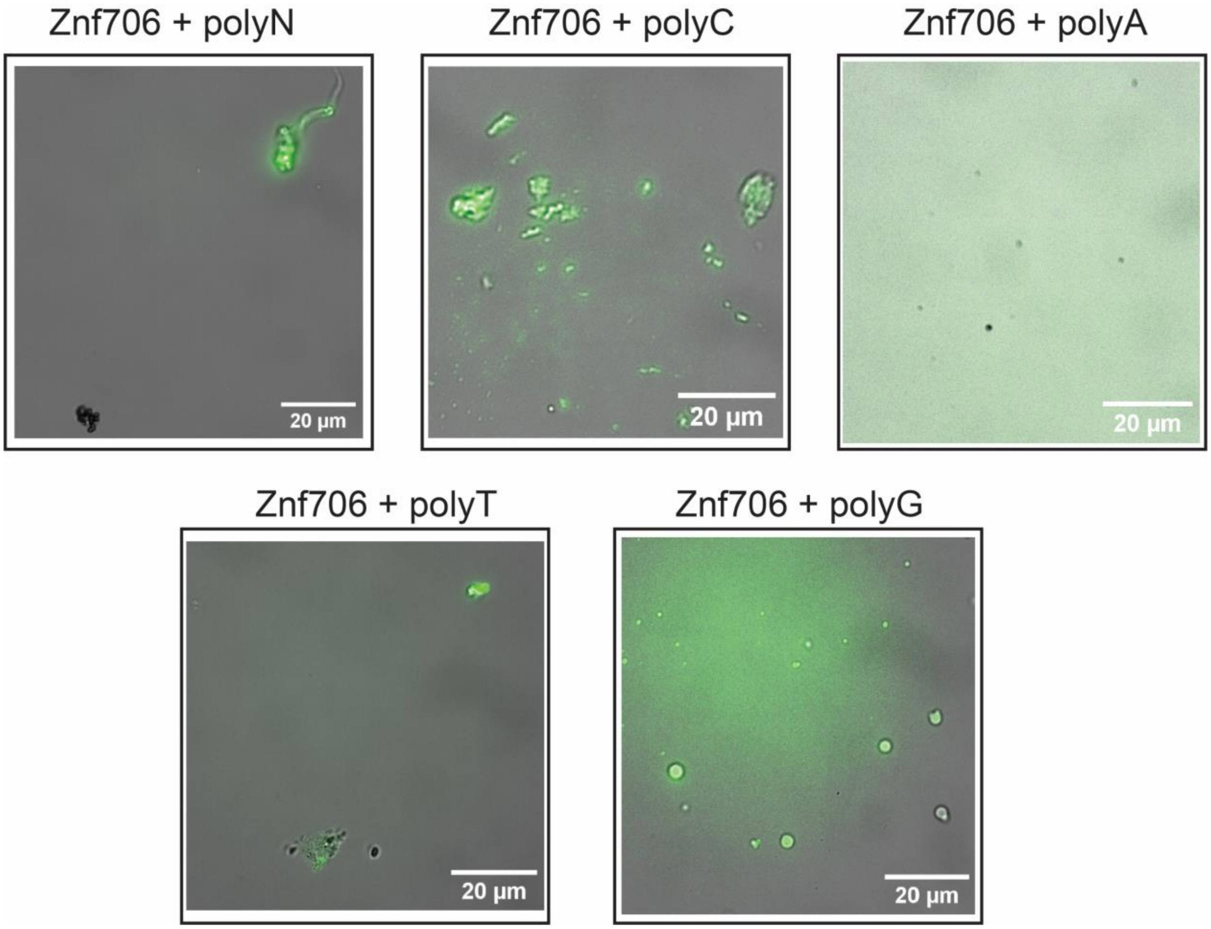


**Figure S26**. Fluorescence imaging to test the formation of LLPT for a sample mixture containing 100 μM Znf706 and equimolar DNA polynucleotides dissolved in 20 mM NaPi, 100 mM KCl, 0.05% $NaN_3$, 7.5% $D_2O$, pH 7.4. The samples were incubated overnight at room temperature before imaging.

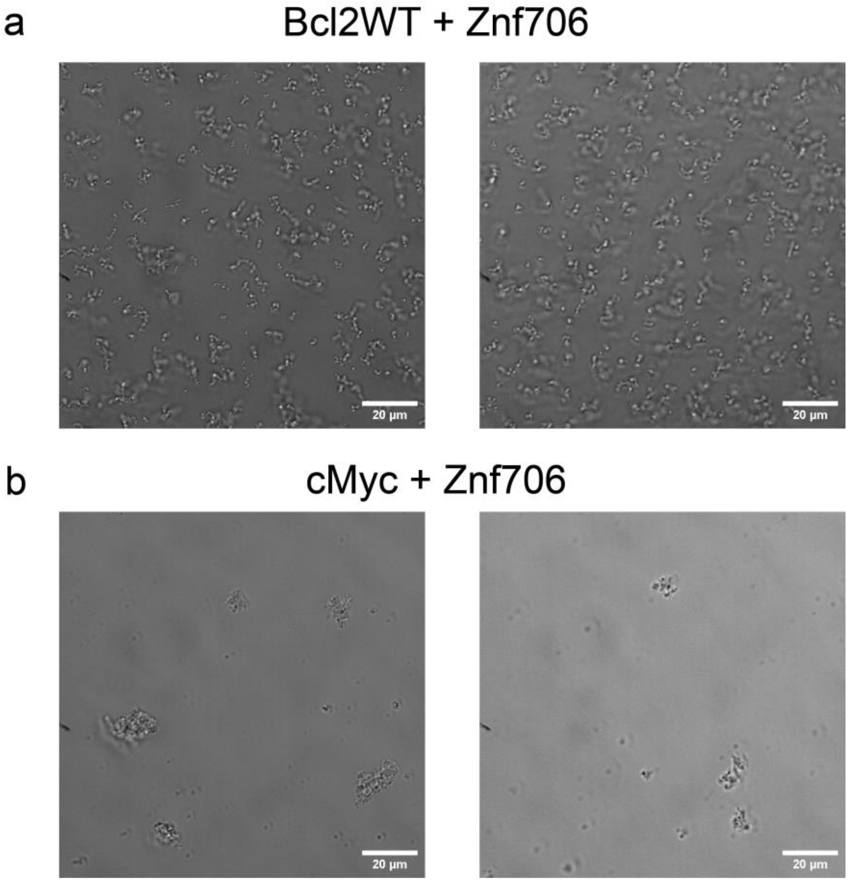


**Figure S27**. DIC images show aggregates in a sample mixture containing 50 µM Znf706 and an equivalent amount of Bcl2WT **(a)** or cMyc **(b)** G-quadruplexes prepared in 20 mM Tris, 100 mM LiCl, pH 7.4.

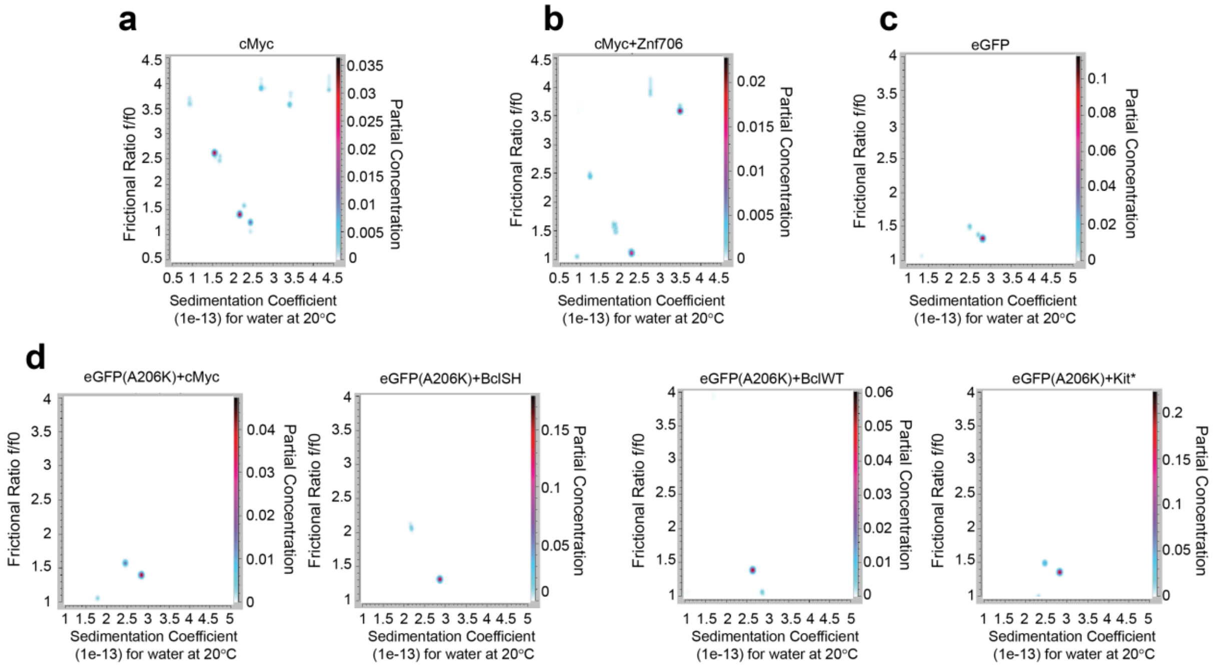


**Figure S28**. Determination of Znf706 and G-quadruplex complex sizes using analytical ultracentrifugation (AUC). cMyc (3.6 µM) dissolved in 20 mM NaPi, 100 mM KCl, pH 7.4 **(a)** was incubated with equimolar Znf706 **(b)** or for 1 hour before AUC measurement (absorbance 260 nm). AUC measurement of eGFP (12.2 µM) mixed without **(c)** or with different G-quadruplexes (12.2 µM) as indicated **(d).** Samples were incubated in the dark for ~1 hour and AUC measurements were done recording the absorbance at 488 nm. AUC data are analyzed in Ultrascan and two-dimensional plots are shown. The partial concentration shown in color in the z dimension represents the abundance of each species.

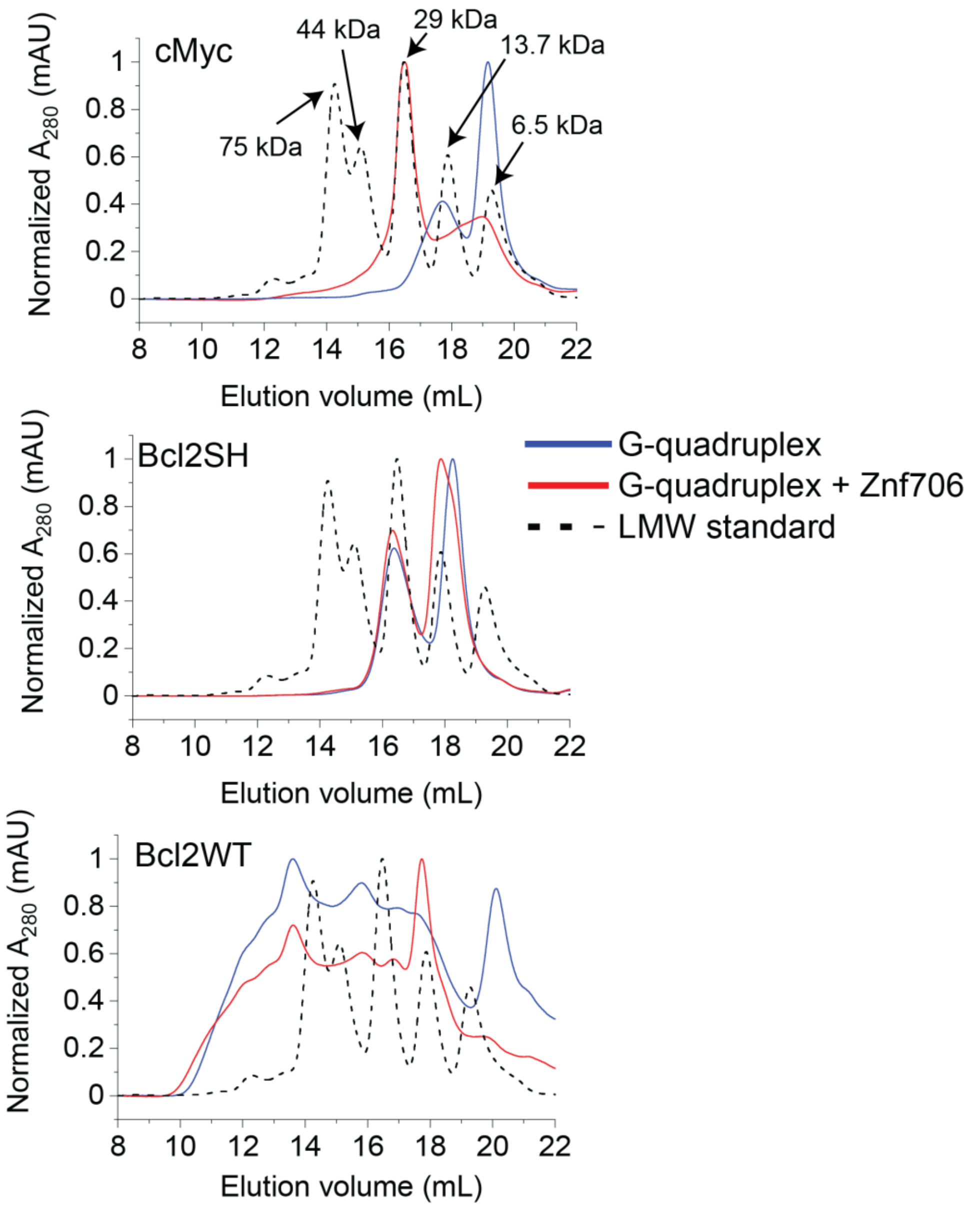


**Figure S29**. Size-exclusion chromatography profiles of three different G-quadruplexes (2.5 μM) mixed with or without an equimolar amount of Znf706. The SEC profile of a low-molecular-weight (LMW) standard is shown in dashed lines. Samples were prepared in 20 mM NaPi, 100 mM KCl, pH 7.4, and incubated ~2 hours before loading to the SEC column.

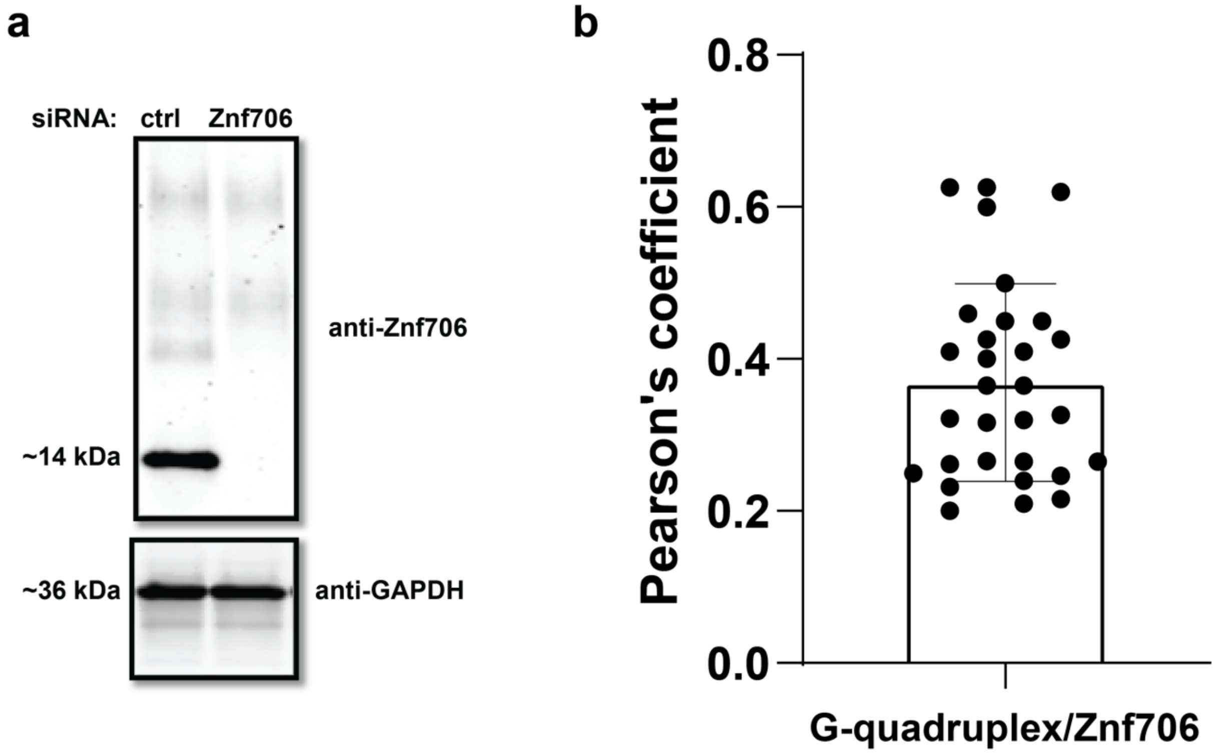


**Figure S30**. **(a)** Western blot analysis of Znf706 using a polyclonal rabbit anti-Znf706 antibody. **(b)** Pearson's coefficient analysis of colocalized Znf706 and DNA G-quadruplex signals in 30 cells was calculated using ImageJ. The fluorescence images used for the Pearson's analysis are shown in **Fig. 6a**.

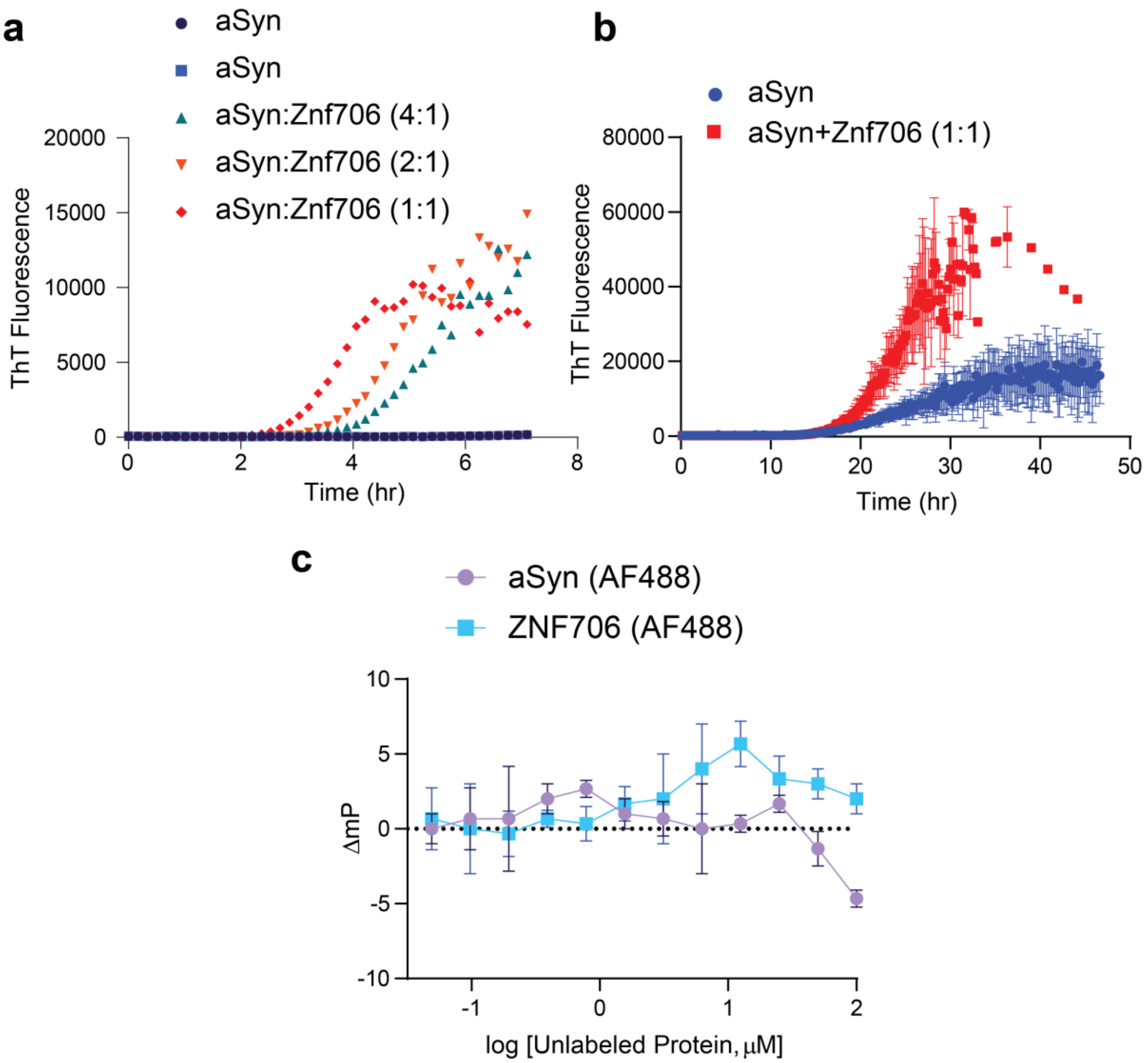


**Figure S31**. **(a)** Monitoring aggregation kinetics of 100 µM α-Synuclein (aSyn) dissolved in 20 mM NaPi, 100 mM KCl, pH 7.4 mixed with and without Znf706 at different stoichiometries as indicated by the colors analyzed by a thioflavin-T fluorescence assay. **(b)** Aggregation kinetics of 50 µM a-Synuclein mixed without (blue) or with (red) 50 µM Znf706. The standard deviation shown is obtained from four replicates. **(c)** Fluorescence polarization assay showing no interaction between Znf706 and a-Synuclein prepared in 20 mM NaPi, 100 mM KCl, pH 7.4. Fluorescence polarization binding curves are obtained for samples containing 100 nM of AF488 aSyn/Znf706 and variable concentration of Znf706/aSyn incubated at room temperature for ~15 minutes.

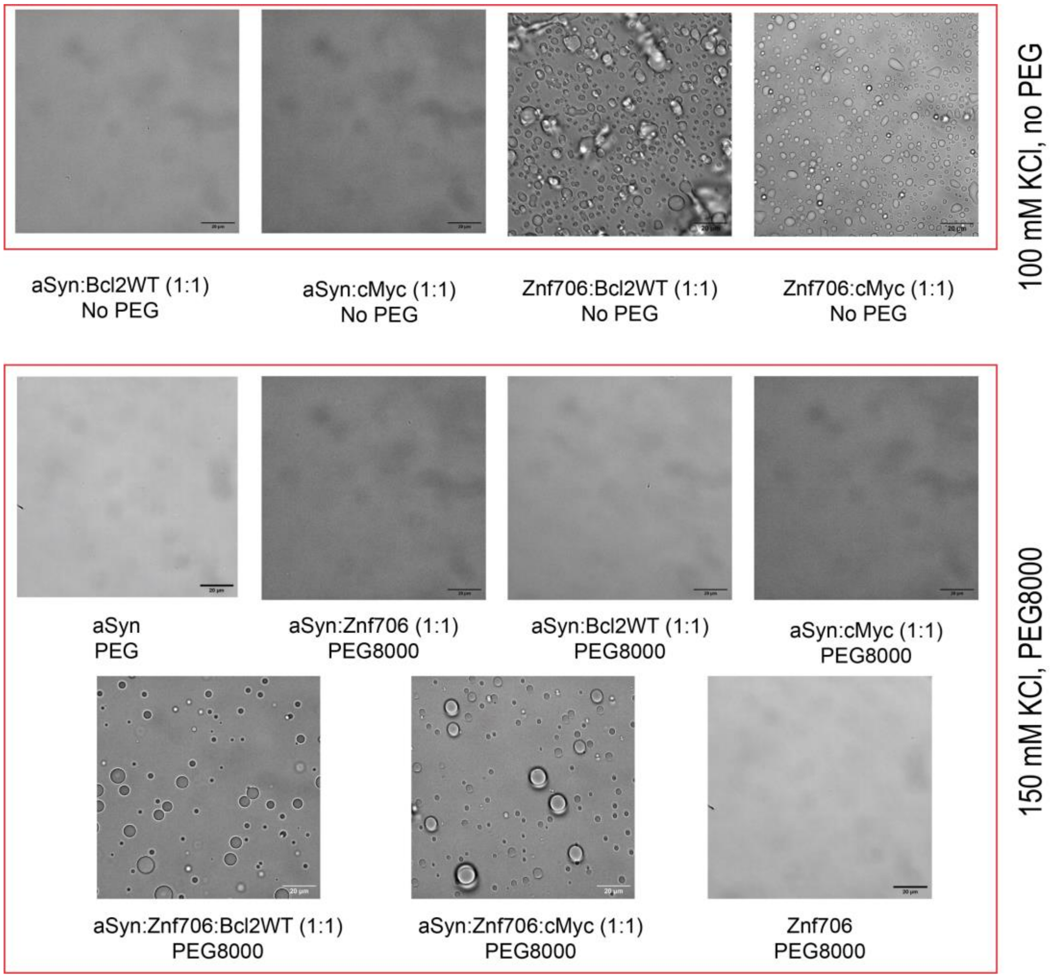


**Figure S32**. Effect of α-Synuclein (aSyn) on Znf706 and G-quadruplex liquid-liquid phase transition. DIC images showing the droplet formation in 100 μM α-Synuclein mixed with or without an equimolar concentration of Znf706 and/or G-quadruplexes. The droplet formation was tested in 20 mM NaPi sample buffer containing no PEG (100 mM KCl) or 10% PEG8000 (150 mM KCl). Scale bar is 20 μm.

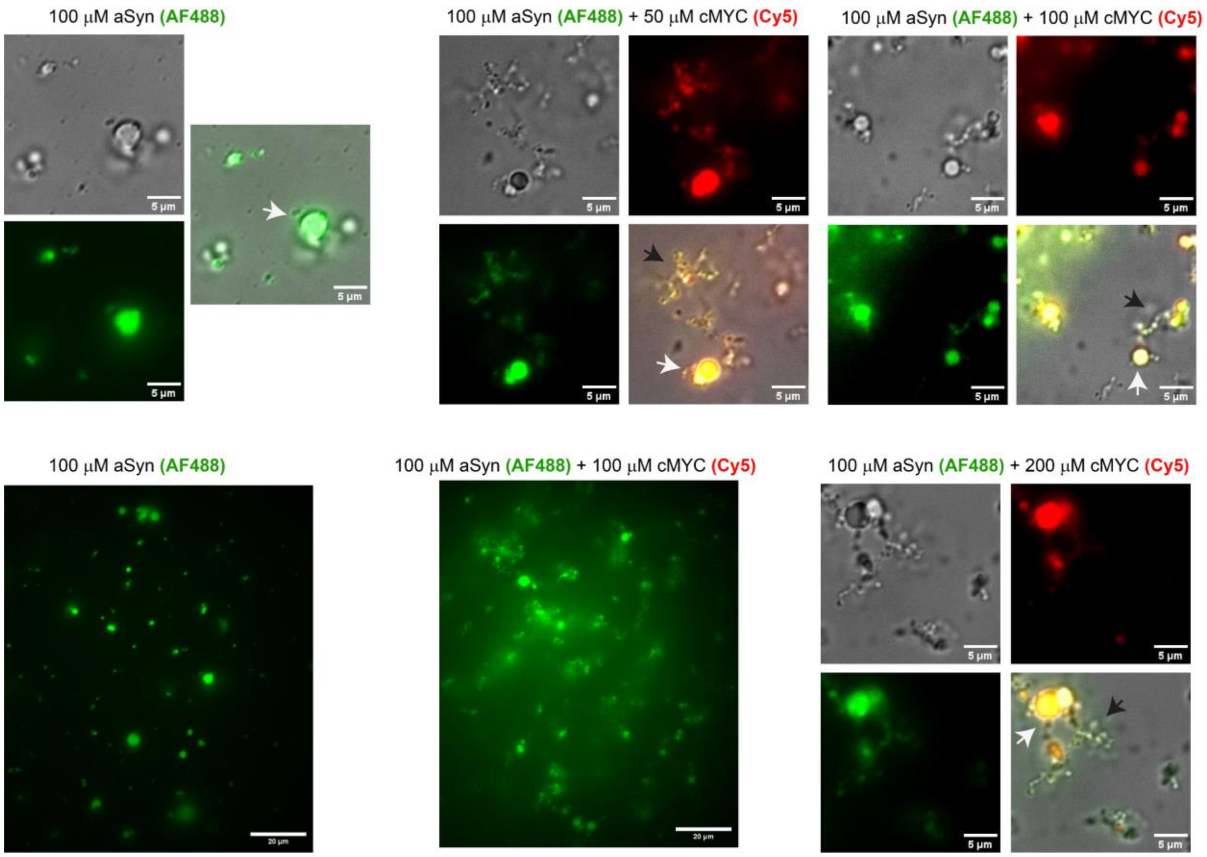


**Figure S33**. Effect of cMyc binding on 100 µM α-Synuclein (aSyn) liquid-liquid phase transition. DIC and fluorescence (aSyn-AF488, green; cMyc-Cy5, red) images showing the droplet or gel-like structure in 100 µM α-Synuclein mixed without or with variable concentrations of cMyc G-quadruplexes as indicated. The droplet formation was tested in 20 mM NaPi, 100 mM KCl, pH 7.4 containing 20 % PEG8000.

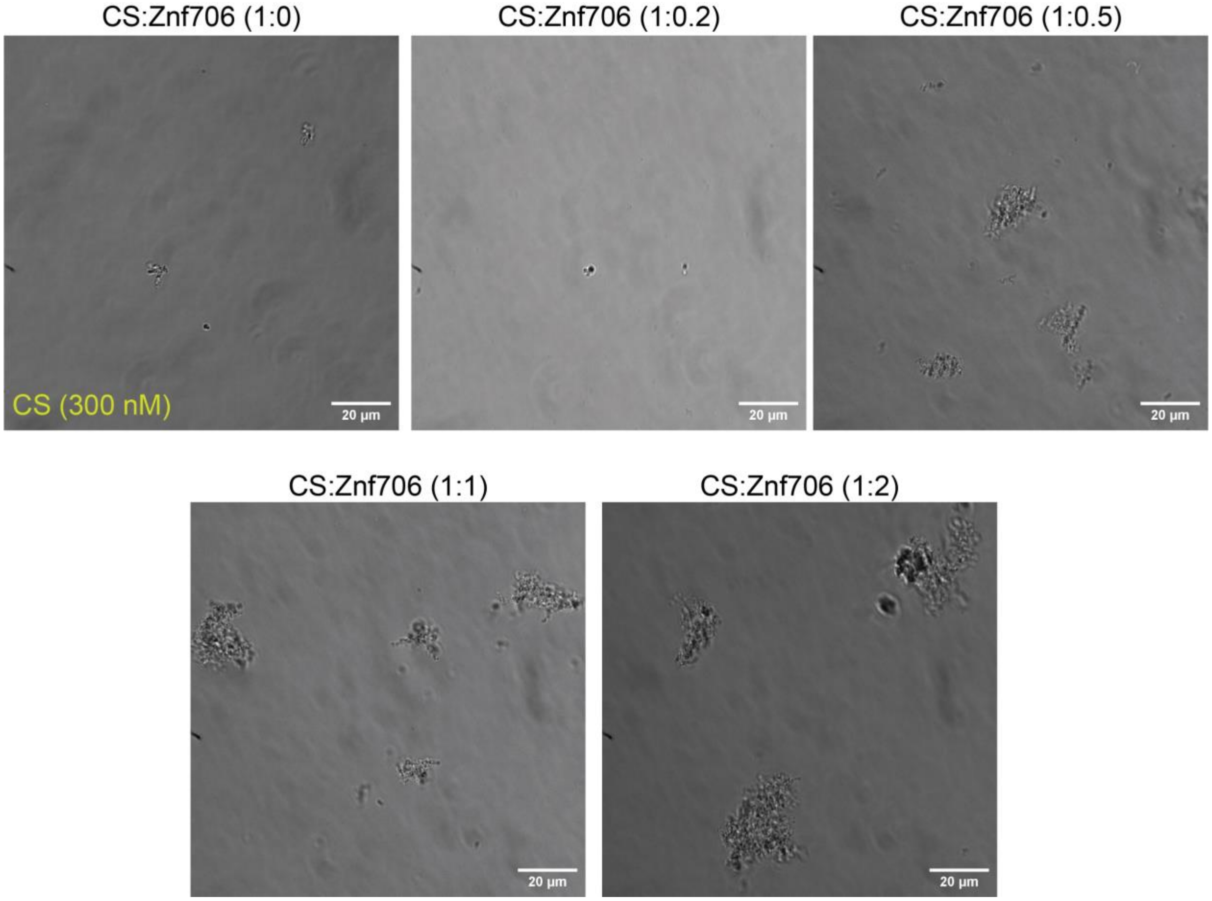


**Figure S34**. DIC images of citrate synthase (CS) amorphous aggregates obtained from thermal aggregation experiments. Thermal aggregation of 300 nM CS mixed without or with the indicated Znf706 concentrations was done at 48 °C.

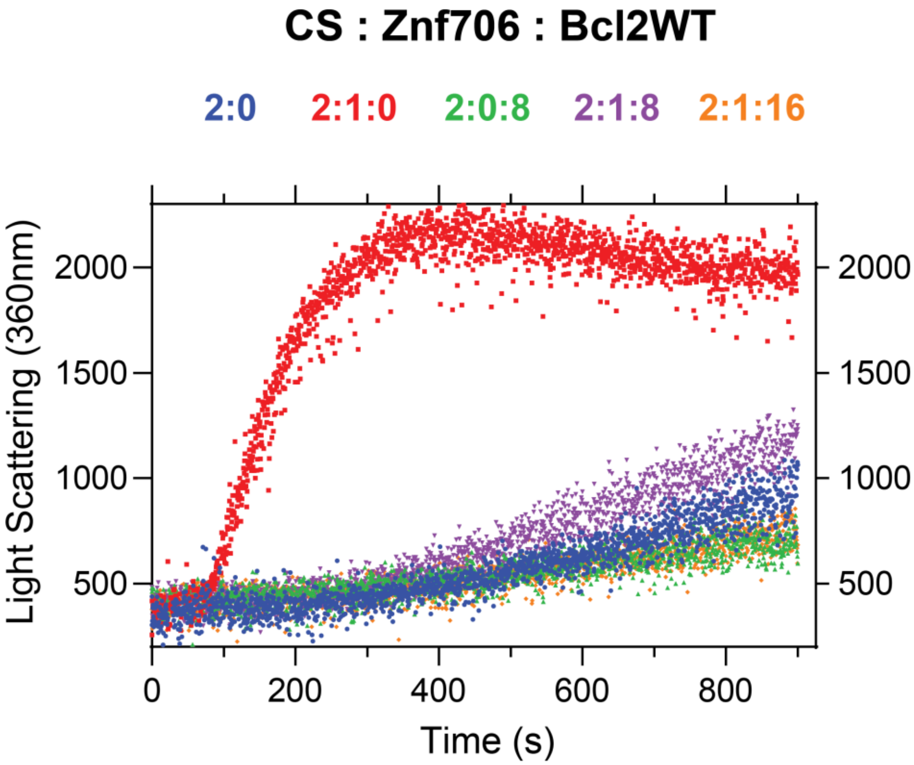


**Figure S35**. Thermal aggregation curves of 300 nM CS mixed without (blue) or with the indicated Znf706 and Bcl2WT G-quadruplex concentrations. The aggregation kinetics of CS was done at 48 °C in 40 mM HEPES-KOH, pH 7.5.

**Video SV1**. LLPT droplets formed by mixing 100 µM of Bcl2WT G-quadruplex and 100 µM of Znf706. The imaging was done post-incubation of the sample mixture overnight at room temperature.

**Video SV2**. LLPT droplets formed by mixing 100 µM of cMyc G-quadruplex and 100 µM of Znf706. The imaging was done post-incubation of the sample mixture overnight at room temperature.
